# The *Caenorhabditis elegans* pangenome elucidates hyper-divergent region gene content to reveal extensive gene copy-number variation and a novel anthelmintic resistance locus

**DOI:** 10.64898/2026.11.17.767430

**Authors:** Lance M. O’Connor, Nicolas D. Moya, Robyn E. Tanny, Amanda O. Shaver, Ayeh Khorshidian, Ryan Pellow, Ofer Rog, Stefan Zdraljevic, Leonid Kruglyak, Erik C. Andersen

## Abstract

*Caenorhabditis elegans* has been a keystone model in biological research since the early 1970s, and it was the first metazoan to have its genome sequenced and assembled in 1998. Over the last few decades, the genetic and genomic resources developed for *C. elegans* made it one of the most powerful models for understanding metazoan biology. Recently, the *C. elegans* community has collected over 2,000 globally distributed wild strains, which after whole-genome sequencing, were found to contain punctuated regions of extreme genetic variation, called hyper-divergent regions (HDRs). HDRs have been described in organisms across the tree of life, including plants and humans, and genes in HDRs appear to be involved in adaptation, such as environmental response genes, parasitism genes, or immunity genes. However, the extreme genetic divergence at these loci complicates short-read alignments, and the genomic content of these regions in non-reference strains is largely uncharacterized. To define the gene content in HDRs, we *de novo* assembled and annotated genomes of 140 genetically diverse strains. These data enabled the construction of the first gene-based pangenome for the species. Among these strains, we characterized the landscape of structural variation and developed novel methods to quantify gene content diversity. We found that both variable copy-number and stable-copy environmental response genes were enriched in HDRs. These results further suggest that these regions have an important evolutionary role in adaptation to heterogeneous environments. We leveraged this pangenome to elucidate gene content in previously identified quantitative trait loci associated with adaptive traits. In one HDR, we found large deletions, spanning multiple genes, that are correlated with resistance to the anthelmintic drug abamectin. The *C. elegans* pangenome provides novel insights into genome and gene family evolution in a genetically diverse animal species, and empowers deeper investigations of genotype-phenotype relationships underlying adaptive traits, particularly in the context of structural and gene copy-number variation.

## Introduction

Reference genomes provide a single representation of the gene content in a species, and do not capture the genetic diversity among individuals. Comparative genomics has historically been performed in relation to a single reference genome, and the completeness and identity of the reference genome can skew estimates of genetic variation among individuals in a species^1–4^. To mitigate biases associated with using a single reference genome, such as the inability to reliably identify non-reference gene content, assembling genomes from multiple individuals of a species enables the comprehensive quantification of gene content variation in a pangenomics framework. Pangenomes represent all unique DNA content in a species and can provide a comparative framework for identifying large sources of genetic variation that cannot reliably be detected using a single reference genome, such as gene copy-number variants, novel genes, and large structural variants (SVs) or genome rearrangements^5^.

With the advent of highly accurate long-read DNA sequencing and advances in *de novo* assembly algorithms, pangenomics studies have leveraged these advances to construct catalogs of genetic diversity from structural variation to gene family evolution and have pioneered recent discoveries of adaptation, domestication, and disease using reference genomes for multiple individuals in a population^5–12^. Over the last five years, most pangenome studies have been constructed in plants^5,13–17^. These studies identified lineage-specific expansions of genes that regulate fruit production^18^ and flowering time^19^, and SVs that regulate resistance to environmental stressors such as ultraviolet radiation, drought, salt, and mold in various ornamental and agricultural plant genera^20–24^. Since the completion of the human genome in 2022^25^, The Human Pangenome Reference Consortium^26^ has made rapid advances in characterizing human genetic variation^6,27–29^, quantifying reference genome biases^1,2,7,8^, and expanding genomic resources for underrepresented human populations^7,9–12^. These studies have revealed important insights into ancestry, population structure, and allelic variation in disease-relevant genes. However, the causality of genetic variants cannot extend beyond statistical association because of ethical constraints. Pangenome studies in other animal systems are limited^30–35^, and many of these animals are constrained by similar ethical issues, low-throughput phenotyping, and a lack of genome editing tools. *C. elegans* provides a unique opportunity to study genome and gene family evolution in an animal system that is equipped with experimental tools, such as high-throughput phenotyping, CRISPR-Cas genome editing, and a short generation time for efficient, controlled genetic crosses, to identify and validate alleles that underlie adaptation.

*Caenorhabditis elegans* is a foundational model species, pioneering discoveries of signal transduction, programmed cell death, gene regulation by microRNAs, and gene silencing by RNA interference^36–40^. However, much of this research uses the laboratory-adapted strain N2, which is a single representation of the *C. elegans* gene content. Recently, the *C. elegans* community has collected over 2,000 wild strains from around the globe, 684 of which are genetically distinct^41^. These strains have been whole-genome sequenced and found to contain punctuated regions of extreme genetic variation, referred to as hyper-divergent regions (HDRs)^42,43^. HDRs have been described in organisms across the tree of life, including other free-living nematodes^44,45^, parasitic nematodes^46–48^, humans^49^, and plants^50,51^. Across all these taxa, HDRs are enriched for reference genes that appear to be involved in adaptation, such as environmental sensing and response genes, parasitism genes, or immunity genes. However, the extreme genetic divergence at these loci complicates short-read sequence alignments, making the content of HDRs in non-reference individuals a mystery. To define the gene content in HDRs, genomes for multiple individuals of a species are needed.

In this study, we *de novo* assembled and annotated genomes of 140 genetically diverse *C. elegans* strains collected from across the globe to construct the first gene-based pangenome for this species. Among these strains, we describe the catalog of structural variation and its correlation with geography, quantified gene family conservation, and identified thousands of strain-specific genes. We developed novel methods to quantify gene content diversity in HDRs and identified both variable copy-number and stable-copy genes enriched in HDRs, further suggesting that these regions might have important evolutionary roles in adaptation to heterogeneous environments. Using pangenomics, we defined HDR gene content in a QTL associated with resistance to the anthelmintic drug abamectin. We found large, multi-gene deletions in wild strains correlated with resistance. This *C. elegans* pangenome is the first to characterize HDR gene content across genetically diverse individuals and provides novel insights into genome and gene family evolution that can empower deeper investigations of genotype-phenotype relationships underlying adaptation.

## Results

### High-quality genome assemblies of genetically diverse *C. elegans* strains

*C. elegans* are globally distributed, and previous studies quantified genetic variation among wild strains, revealing population structure as well as large chromosome-level selective sweeps^42,52^. Wild strains largely cluster into three genetically distinct groups, referred to as the Hawaii group, the Pacific group, and the Global group^42^. The Hawaii group contains some of the most genetically diverged strains from the N2 reference strain, and the largest group, the Global group, contains strains that have chromosome-level selective sweeps. The immense genetic variation across *C. elegans* strains has motivated the community to amass a collection of over 2,000 wild *C. elegans* strains, and these strains cluster into 684 isotypes (genome-wide haplotypes, strains that are >99.97% identical to each other)^41^. We chose 140 isotype reference strains from different clusters based on sharing a high proportion of SNVs for PacBio HiFi long-read sequencing to best capture the genetic variation in the species (Figure 1a,b, Supplementary Figure 1). Among these 140 wild strains, we obtained a mean fold-coverage of 54.56x and an average read length of 13.56 kb (Supplementary Table 1). We used hifiasm^53^ in a custom Nextflow^54^ pipeline to generate *de novo* assemblies to ensure reproducibility (see *Methods*). The pipeline produced highly contiguous assemblies with an average N50 of 8.58 Mb (min 1.16 Mb, max 18.00 Mb) and an average N90 of 2.8 Mb (min 200 kb, max 13.09 Mb) (Figure 1c, Supplementary Table 1, Supplementary Figure 2). Many of these wild strain assemblies exhibit near chromosome-level contiguity, including 20 assemblies that have an N50 greater than 14 Mb, comparable to the canonical N2 reference genome (scaffold N50 of 15.2 Mb)^55–57^. Assembly sizes ranged from 102.35 Mb to 112.19 Mb (average 106.58 Mb), similar to both the N2 reference genome (100.2 Mb) and the updated gap-free reference genome, CGC1 (106.4 Mb) (Figure 1c, Supplementary Table 1). To assess genome completeness based on the presence of conserved single-copy orthologs among nematode species, we ran genome BUSCO^58^ and found that all genome assemblies have a BUSCO completeness score of 99.2% or greater (Supplementary Table 1).

**Figure 1.**
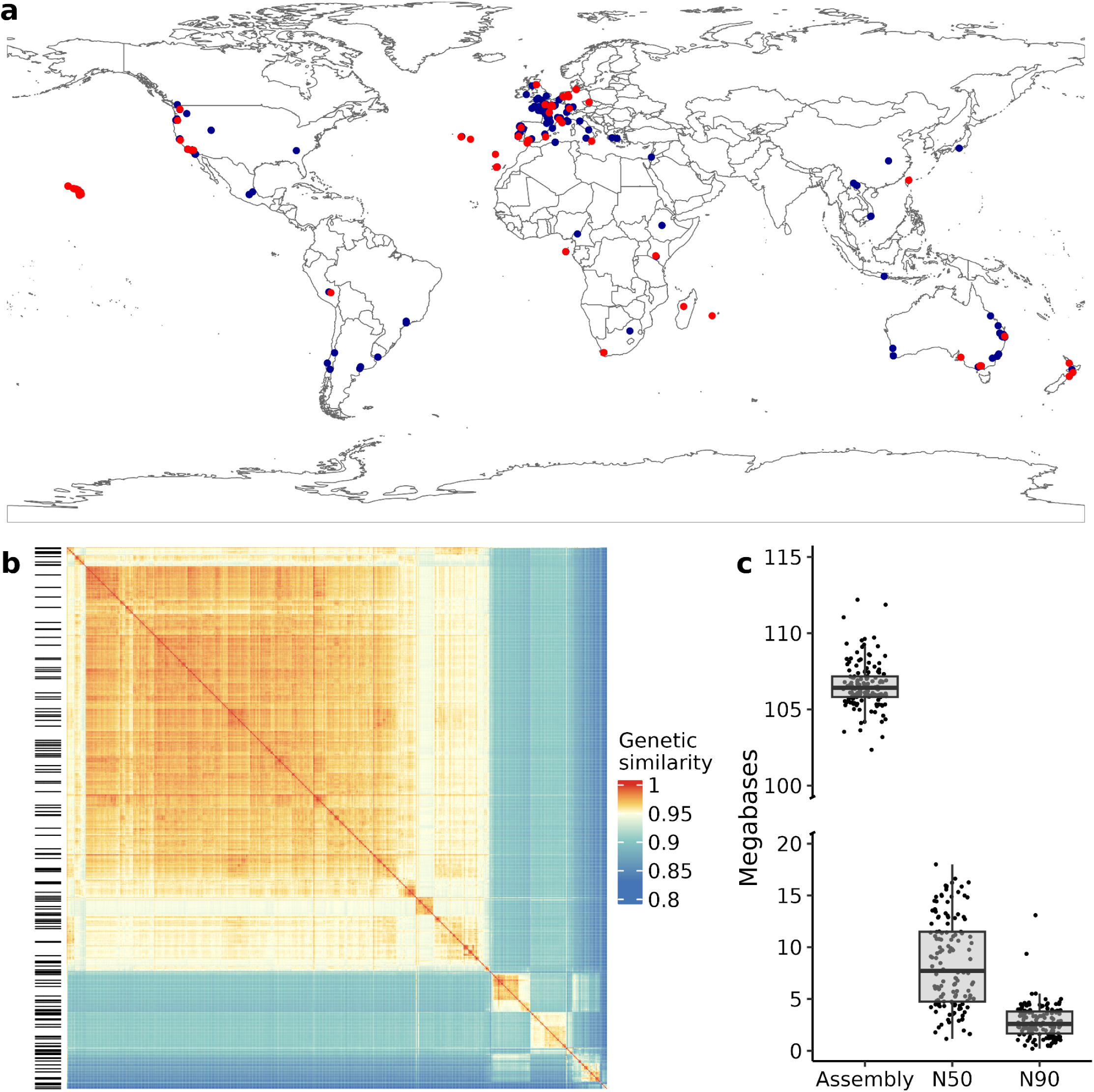
PacBio HiFi sequencing and *de novo* assemblies of 140 wild *C. elegans* strains selected to maximize captured genetic variation. **a)** The geographic collection sites of 684 *C. elegans* isotype reference strains. Each point represents a strain, with strains selected for PacBio HiFi sequencing in red. Six strains that do not have geographic information are not displayed. b) A genetic relatedness matrix displaying the proportion of shared, alternative single-nucleotide variants between any pair of the 684 isotype reference strains. Strains selected for PacBio HiFi sequencing are indicated with black, horizontal lines on the left y-axis. c) Genome assembly statistics for 140 wild *C. elegans* strains. Box and whisker plots display the median value for each metric on the x-axis. The upper and lower whiskers correspond to the first and third quartiles, respectively.

### Wild *C. elegans* strains exhibit extensive structural variation

Using the 140 genome assemblies, we sought to understand how genetically diverse *C. elegans* strains structurally differ from the N2 reference genome. Previous studies found that self-fertilizing *Caenorhabditis* species, such as *C. elegans* and *Caenorhabditis briggsae*, have undergone extensive intra-chromosomal rearrangements of single-copy orthologs shared between the species in comparison to their outcrossing relatives^59,60^. Yet, comprehensive estimates of conspecific structural variation in *C. elegans* have not been well documented. We used the Phased Assembly Variant caller (PAV)^61^ (see *Methods*) to identify SVs among 141 strains (140 wild strains and CGC1) in relation to the N2 reference genome. Among all 141 strains, each strain has 3,523 deletions, 60 inversions, and 4,700 insertions greater than 50 bp in relation to the N2 reference strain on average (Supplementary Figure 3a). Across all strains, insertions account for over 50% of SVs identified in relation to the N2 reference genome (Supplementary Figure 3b). The updated N2 reference genome, CGC1, has over 6 Mb of sequence that is absent from the N2 reference genome^62^, which likely indicates the wild strain assemblies have captured some of the added sequence in CGC1 that is absent in N2 and might explain the large proportions of insertions. The variability in the number of SVs identified among wild strains also correlates with total span of structural variation. The extent of structural divergence in relation to the 100.2 Mb N2 reference genome ranges from less than 4 Mb to over 12 Mb (Figure 2a). Historically, genetic diversity in a population is quantified using SNV counts^42^. Population-scale comparative genomics studies have revealed punctuated loci that are enriched for SNVs, known as HDRs. We found that SNV counts highly correlate (R^2^ = 0.972) with SV counts among wild strains (Supplementary Figure 4), and we were curious if SVs were also enriched in HDRs. We found that deletions (2.97 fold enrichment), inversions (2.23 fold enrichment), and insertions (2.58 fold enrichment) were all enriched in HDRs (Supplementary Figure 5). These results indicate that not only are HDRs enriched for allelic variation sourced from SNVs but also structural changes that might result in gene copy-number differences, intra-chromosomal rearrangements, and non-reference gene content.

**Figure 2.**
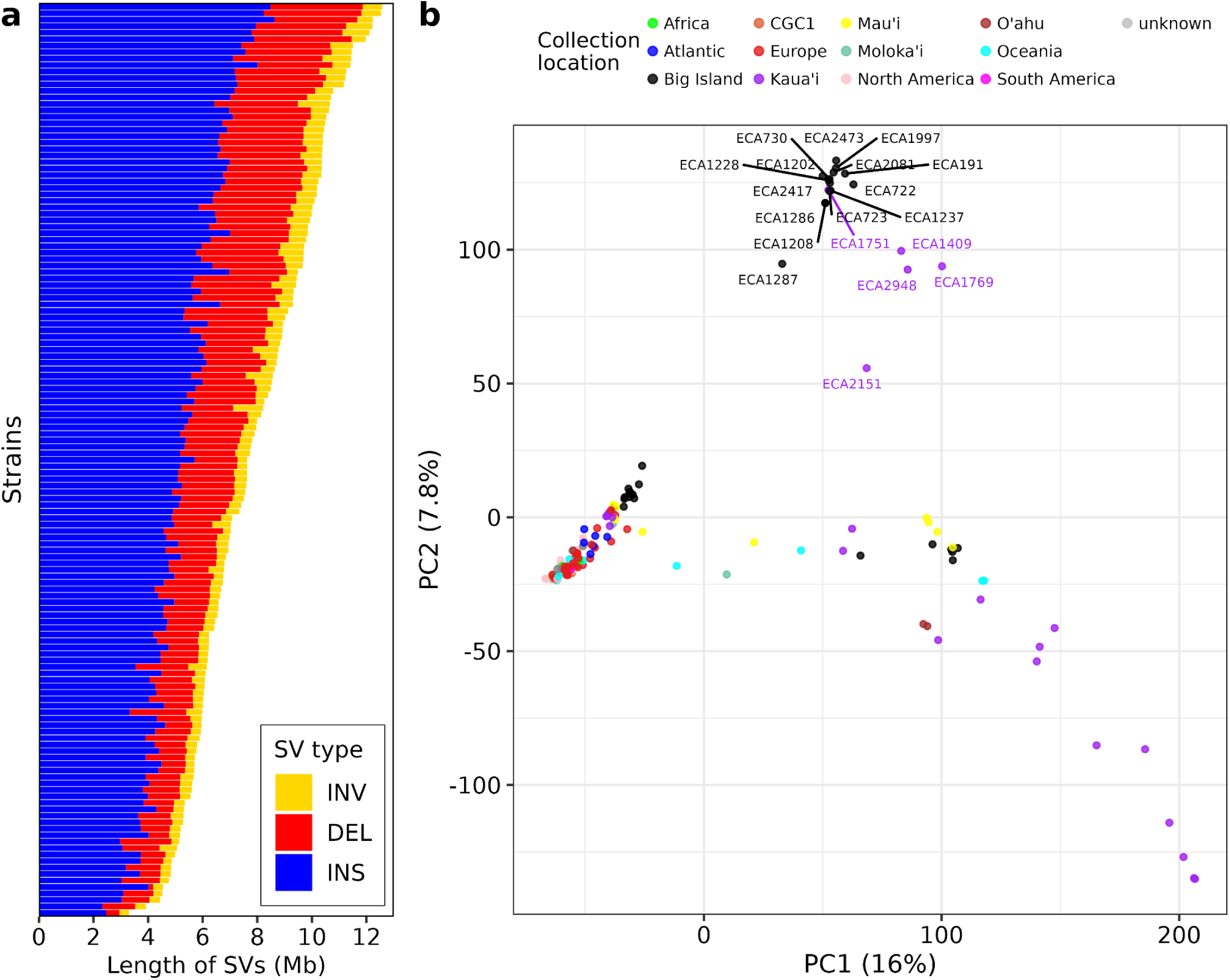
*C. elegans* wild strains are structurally diverged from the reference genome. **a)** Cumulative length of structural variants for 141 strains of *C. elegans*. **b)** Principal components (PCs) of all structural variants (MAF >= 0.05) with strains colored by the sampling location.

Although SV count and span vary among wild strains, SVs are mostly found on chromosome arm domains where recombination is most frequent (Supplementary Figure 6)^63^. Across all SVs identified, 18.6% percent of the N2 reference genome is covered by at least one deletion, 6.8% percent is covered by at least one inversion, and 43.1% percent of the N2 reference genome length is contained in unique insertions. We found SVs that are common among wild strains overlap with many N2 protein-coding genes (Supplementary Figure 7). The SVs that overlap within two kb of N2 reference genes account for 79.0% of deletions, 85.0% of inversions, and 79.1% of insertions that are common among wild strains. We found that these common SVs were primarily in GPCRs, F-box genes, protein kinases, C-type lectins, and nuclear hormone receptors (Supplementary Table 3), similar to findings in a recent study^64^. These gene classes are some of the largest in *C. elegans*^65^, and we found that N2 reference genes with overlapping SVs have lower expression levels among wild strains than N2 reference genes that are structurally conserved (Supplementary Figure 8, see *Methods*)^66^. Because most SVs overlap with protein-coding genes involved in environmental response and these genes have lower expression levels, these SVs might have a functional role in shaping gene content diversity and regulation in large, redundant gene classes among wild strains.

Strains collected from the Hawaiian islands have higher nucleotide diversity than non-Hawaiian strains^42,67^, which correlates with Hawaiian strains harboring the most SVs (Figure 2a, Supplementary Figure 3a, Supplementary Table 2) Previous studies revealed that strains collected from Hawaii display population structure using SNVs^42,67^. To investigate the population structure of strains in our pangenome strain set through the lens of structural variation, we performed principal component analysis (PCA) on all common SVs (MAF greater than or equal to 0.05). PCA revealed some Hawaiian strains collected from the Big Island and from Kaua’i have greater genetic similarity to one another than to all other strains (Figure 2b). The observed structure is consistent across principal components for each SV type (*i.e.*, deletions, inversions, and insertions) (Supplementary Figure 9). The 19 Hawaiian strains that cluster in PCA share 61 unique SVs (47 deletions, 12 insertions, and one inversions) that are not found in other wild strains (Supplementary Table 4). However, not all Big Island and Kaua’i strains cluster with each other, which is reflective of intra- and inter-island relatedness patterns, where strains from different islands might be more related to each other than strains from the same island. These results are consistent with previous findings of population structure among Hawaiian strains and spatial patterns of relatedness^42,67^. These patterns might reflect human influence on the global dispersal of *C. elegans* or shared niche-specific adaptations found across islands^67^. Alternatively, these shared SVs might be from shared ancestry and inter-island dispersal via hitchhiking off human travel.

### *C. elegans* pangenome reveals extreme gene content variation among wild strains

The *C. elegans* N2 reference strain has approximately 20,000 protein-coding genes and has served as the representative gene content for the species. However, HDRs and the extensive structural variation among wild strains reveal that these strains have gene content differences from the N2 reference strain. Here, we sought to understand the level of gene content conservation among wild strains. To identify protein-coding genes, we used BRAKER3^68^ to predict a total of 3,215,467 protein-coding gene models in the 141 (140 wild strains and CGC1) assemblies (see *Methods*). Next, we clustered genes into groups of orthologous genes (orthogroups) using OrthoFinder (see *Methods*)^69^. OrthoFinder identified 55,159 orthogroups among the 142 *C. elegans* strains (N2, CGC1, 140 wild strains) (Supplementary Table 5). Of these orthogroups, 12,315 (22.32%) are single-copy orthologs and are found among all 142 strains. The other 78.68% of orthogroups are found in some but not all strains or are multi-gene orthogroups comprising likely paralogs. To quantify gene conservation, all orthogroups were classified into pangenome gene sets based on the proportion of strains that contribute an ortholog. A total of 2,131,084 genes were classified as core (found in all 142 strains), 1,067,222 as accessory (found in some but not all strains), and 17,161 as private (found in a single strain) (Figure 3a). On average, each wild strain has 15,008 genes that contribute to the core pangenome, 7,516 that contribute to the accessory pangenome, and 121 genes that contribute to the private genome (Supplementary Figure 10). We observed consistency in the proportion of genes that contribute to the core pangenome per strain, with an average of 66.29% (min 65.0%, max 75.1%) (Supplementary Figure 10), indicating a lack of widespread gene model errors or DNA contamination. Despite the slight variability in the number of genes for each strain of *C. elegans*, we found that wild strains with more total genes do not necessarily have more private genes (Supplementary Figure 11). The mean number of genes predicted among all strains is 22,644, with a minimum of 19,984 (the N2 strain, WS283 WormBase) and a maximum of 23,090 (the ECA2377 strain). The highest count of private genes (540) in the N2 strain is likely because those gene models have undergone decades of manual curation and some of these private genes might have non-canonical or difficult-to-predict gene structures missed by BRAKER3. Only 14,998 of the 19,972 canonical N2 reference genes (excluding mitochondrial genes) are conserved among all wild strains, the other 4,974 genes are found at variable frequencies in the population or they are unique to the N2 reference strain.

**Figure 3.**
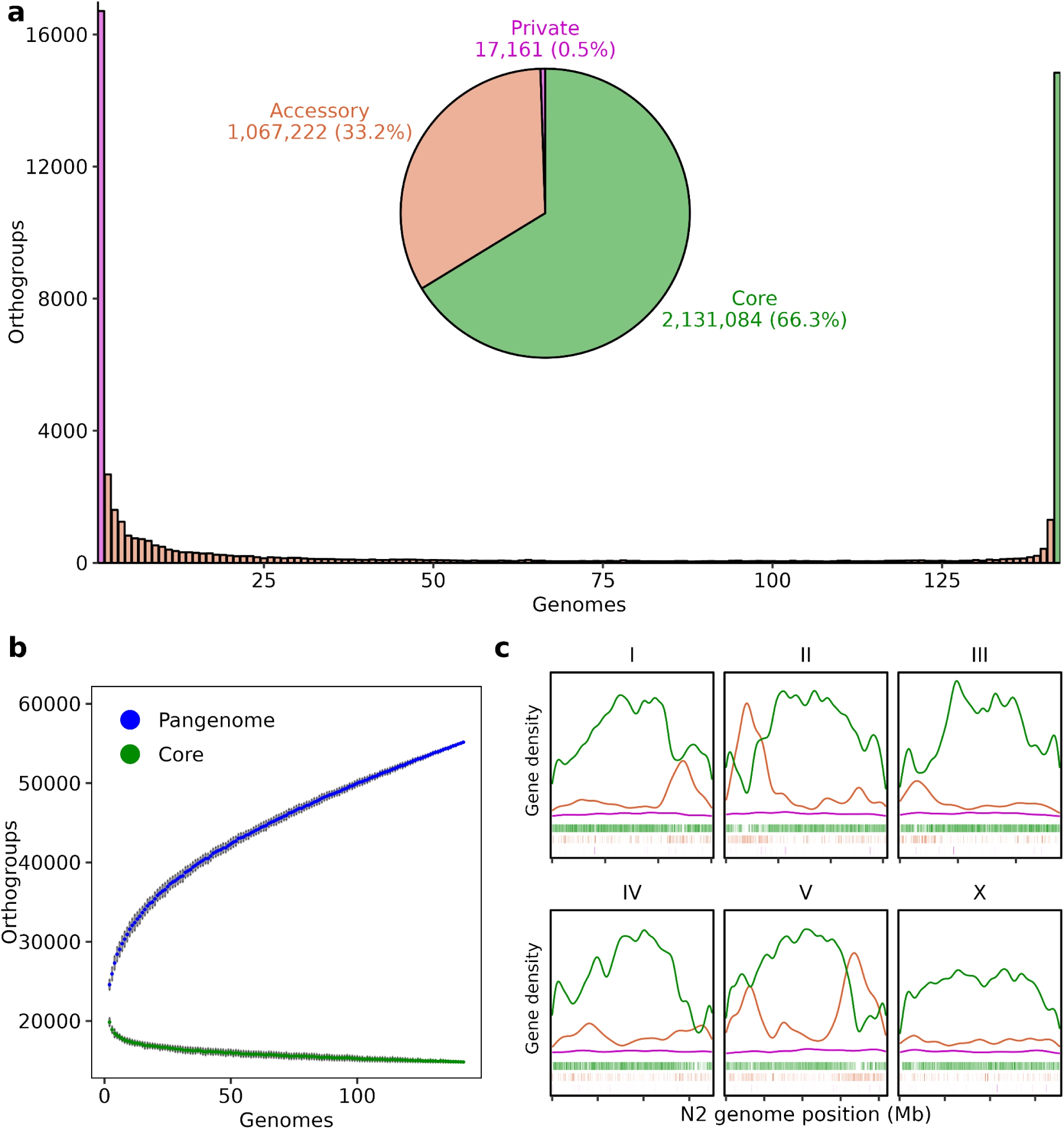
*C. elegans* pangenome gene set classification. **a)** Histogram of pangenome gene sets. Pie chart inset displays the proportion of genes contributing to each gene set, with the number of genes shown in text. **b)** Rarefaction curves of all orthogroups (pangenome) and core orthogroups. Colored points display the mean value for each subsample of genomes (between 2 - 142) after 100 iterations, and error bars represent one standard deviation from the mean. **c)** Smoothed kernel-density plot of N2 reference genes colored by gene set and faceted by N2 chromosome. Tick marks on the x-axis increase every five megabases starting from zero. N2 reference genes are plotted as rectangles, colored by their gene set classification.

To determine if this set of 142 strains captures the gene content diversity of the *C. elegans* species, we used rarefaction to quantify the saturation of orthogroup discovery. We found that the pangenome contains most (if not all) core orthogroups, because new core orthogroups are not readily discovered with the addition of new wild strains (Figure 3b). However, many new orthogroups, agnostic to gene set classification, are discovered as new strains are added (Figure 3b, Supplementary Figure 12), indicating that wild strains are diverse and a single reference genome cannot capture the genetic diversity in the species. The observed growth of orthogroups as new strains are added to the pangenome is primarily driven by the private pangenome (Supplementary Figure 12). Because rarefaction curves are built using orthogroups, and not genes, the private pangenome will continuously grow at a steady rate if newly added strains are genetically diverse from the pangenome strain set. We purposefully selected strains that represent diverse relatedness clusters in the species, so new private orthogroups are readily discovered with the addition of each wild strain. We anticipate that if we add strains to the pangenome that are closely related to the current strains, then many of the private orthogroups would become accessory orthogroups.

Because of the observed population structure in our pangenome strain set (Figure 2b), we clustered orthogroups based on BUSCO-inferred relatedness to identify relatedness-group-specific accessory orthogroups (Supplementary Figure 13). We found that 18 of the 19 Hawaiian strains that cluster in SV PCA also cluster based on shared accessory orthogroups (Supplementary Figure 14). These data revealed the loss of *sra-2*, *sra-9*, *T09F3.7*, and *W02B12.16* specific to these 19 strains. The loss of *sra-2* and *sra-9* is caused by a 6.9 kb deletion found on chromosome II that is shared among these strains (Supplementary Figure 15). Both *sra-2* and *sra-9* are seven-transmembrane serpentine receptors predicted to be involved in the detection of chemical stimuli, and *sra-9* has been shown to modulate EGF/MAPK signaling required for proper vulval development^70^. This lineage-specific loss of *sra-9* involved in vulval cell fate determination might reveal alternative mechanisms of cell-fate specification during development in these 19 Hawaiian strains.

These pangenomic analyses revealed that only 75% of the N2 reference genes are conserved across wild strains in the pangenome strain set. The N2 reference genes that contribute to the core pangenome are more often found on chromosome centers, and N2 reference genes that contribute to the accessory pangenome are more often found on chromosome arms, overlapping with HDRs (Figure 3c). This observed distribution is consistent with previous studies that have found chromosome centers are more gene-dense and harbor many essential genes, and chromosome arms are hotspots for recombination, repetitive content, less gene-dense, and harbor large, redundant gene families^63,71–73^. Using the pangenome strain set, we can define the gene classes that are in HDRs among all wild strains and quantify their conservation with N2 reference HDR genes to understand their evolution.

### Hyper-divergent regions are enriched for environmental response genes found at intermediate frequencies in the pangenome

We cannot use short-read data to reliably define the genomic content in HDRs among wild strains because the extreme divergence in these regions complicates alignments. However, we can define HDR genomic content for 140 wild strains using the *de novo* genome assemblies. HDR boundaries are in the N2 reference genome coordinate system and can be lifted over to wild strain genome coordinates using genome-genome alignment (see *Methods*, Supplementary Figure 16). Upon lifting over HDRs to wild strain genomic coordinates, we observed that wild strain HDRs were correlated with, but rarely identical to, their respective sizes in N2 coordinate space (Supplementary Figure 17). This discrepancy in size is consistent with HDRs harboring many insertions and deletions found in HDRs in relation to the reference genome (Supplementary Figure 15). We found that the ECA2199 Hawaiian strain has the most HDRs (319) and the ECA2565 Australian strain has the fewest HDRs (15), and the average number among all 140 wild strains is 139 (Supplementary Table 6). In addition, the span of HDRs is also highly variable. We found the average HDR size is 33.59 kb and the largest HDR is 1.22 Mb (ECA1769 strain) (Supplementary Table 7). The average cumulative span of HDRs is 4.74 Mb, the shortest span is 0.29 Mb (NIC2 strain), and the longest span is 15.46 Mb (ECA2199 strain) (Supplementary Figure 18), correlated with the extent of structural variation among these wild strains (Figure 2a, Supplementary Figure 3). Genes in the wild strains were classified as present in HDRs or not based on their genomic locations, which revealed that 206,224 out of 3,173,110, or 6.5% (min 0.4%, max 19.1%) of all wild strain genes are in HDRs (Supplementary Figure 18, Supplemental Table 6). Furthermore, 70,320 out of 2,101,058 (3.35%) core genes, 133,445 out of 1,055,492 (12.64%) accessory genes, and 2,459 out of 16,560 (14.85%) private genes are found in wild strain HDRs (Figure 4ab, Supplementary Table 8). These data reveal that the majority of wild strain HDR genes are found at variable frequency in the population and contribute to the accessory pangenome (Figure 4ab).

It is important to identify if specific gene classes are enriched in HDRs because it might provide insights into the functional importance of these regions and their potential roles in environmental adaptation. We know that environmental response genes in the N2 reference genome are enriched in HDRs. However, because of the immense structural variation in HDRs, it is likely that wild strains contain genes that are not found in the N2 reference genome. To identify the types of genes in wild strains that lack orthology with the N2 reference strain, we used InterProScan (IPR)^74^ to annotate functional protein domains (Supplementary Table 9). We found that the most abundant IPR terms among wild strain HDR genes are G protein-coupled receptors (GPCRs), F-box proteins, C-type lectins, and nuclear hormone receptors (NHRs) (Supplementary Figure 19). Next, we sought to identify if these genes are enriched in HDRs in relation to the rest of the pangenome. We performed functional enrichment analyses (see *Methods*) on core, accessory, and private genes in wild strain HDRs in relation to the entire pangenome and isolated the 35 most enriched IPR terms in HDRs among all three gene sets (Figure 4c, Supplementary Table 10). The elevated nucleotide diversity and enrichment for environmental response genes in HDRs relative to the rest of the genome suggests that these regions are under different selective pressures^42^. Because these regions contain alleles that are found at intermediate frequencies in the population, these haplotypes might be maintained by balancing selection, whereas genes outside of HDRs are more conserved in the species and likely experience stronger purifying selection to maintain the function of genes essential for survival. We expected to find the majority of gene classes enriched in HDRs to contribute to the accessory pangenome, and few, if any, to be enriched in the core pangenome. However, despite being in HDRs, we found NHRs and cytochrome P450s that contribute to the core pangenome are enriched in HDRs (Figure 4c). These results indicate that not only do HDRs contain genes that are core to the species, but NHRs and cytochrome P450s are in HDRs more frequently than expected by chance (see *Methods*). These core genes might be single-copy orthologs that have been conserved, despite being in regions of extreme divergence or they might be an artifact of ortholog grouping, where multi-gene orthogroups have some members in HDRs and some outside of HDRs. Additionally, we found wild strain HDR genes that contribute to the accessory and private pangenomes are enriched for F-box genes, GPCRs, and C-type lectins (Figure 4c, Supplementary Figure 20). These gene classes have many members and are involved in environmental sensing and response. The conservation (NHRs and cytochrome P450s) and plasticity (F-box genes, GPCRs, C-type lectins) of HDR genes might reflect complex selection regimes that act to diversify or conserve different environmental sensing and response gene classes in different environments. We found that the pangenome strain set exhausted novel discovery of these gene classes (Supplementary Figure 21). These five gene classes are the same types of genes that are enriched in HDRs in the N2 reference genome^42^. However, given that GPCRs, F-box genes, and C-type lectins are enriched in the accessory pangenome, many of these genes might not be orthologous to N2 reference strain HDR genes.

**Figure 4.**
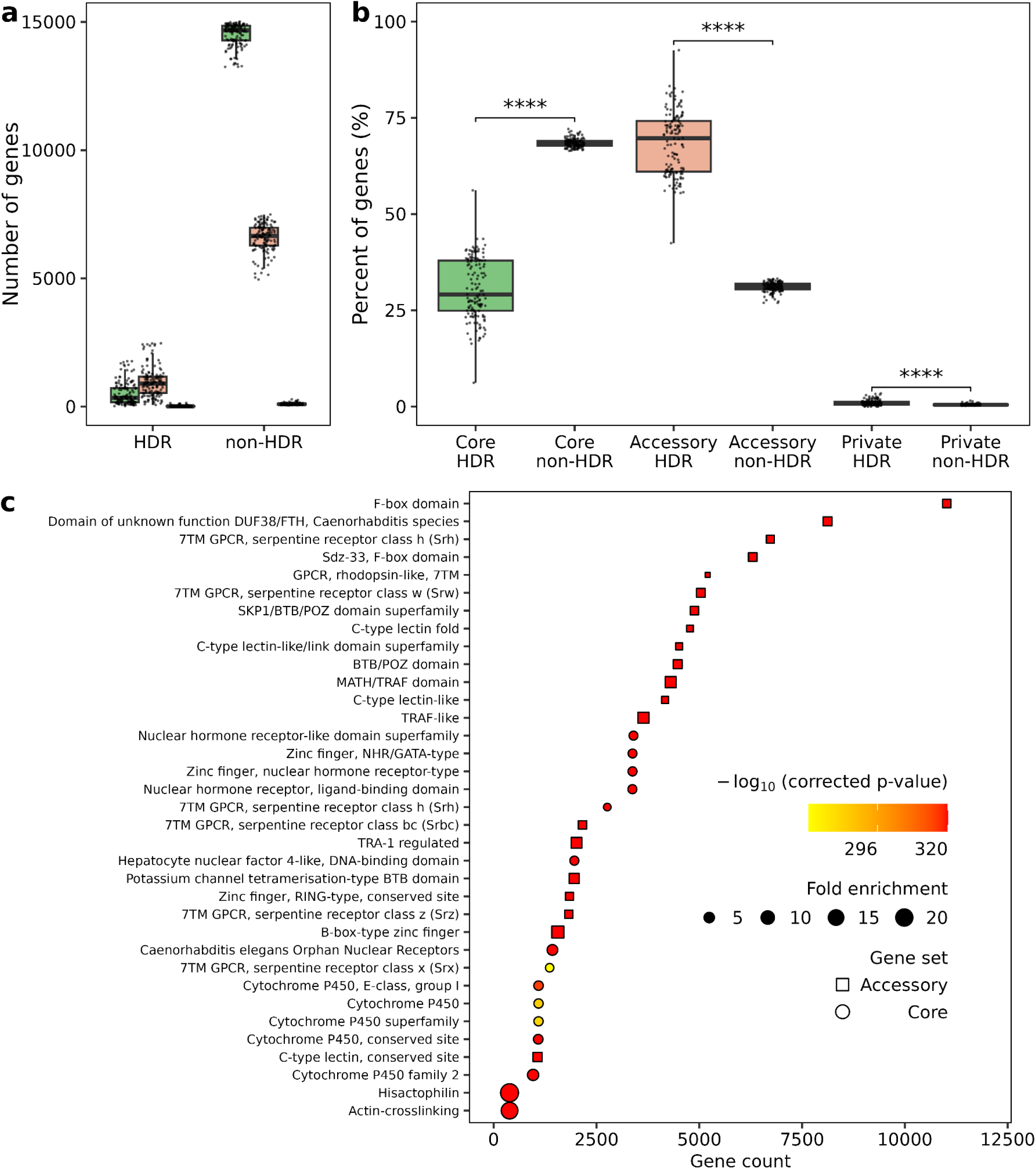
Wild strain HDRs are enriched for genes involved in environmental sensing and response. **a)** The number of genes in HDRs or outside of HDRs (non-HDR) categorized by gene set (core = green; accessory = orange; private = purple). **b)** Of genes in or outside of HDRs for each strain, the percent composition of each gene set is displayed. Box and whisker plots display the median value for each metric on the x-axis. The upper and lower whiskers correspond to the first and third quartiles, respectively. Statistical significance was determined using the Wilcoxon signed-rank test with Benjamini-Hochberg correction. \*\*\*\**p* < 0.0001. **c)** IPR functional enrichment analysis using a hypergeometric statistical test of wild strain genes in HDRs using all genes in the pangenome as the background set. Corrected *p*-values are FDR–adjusted using Benjamini-Hochberg. Displayed are the top 35 most significantly enriched IPR annotations in HDRs across all pangenome gene sets.

### Variable copy-number and stable-copy environmental genes are enriched in hyper-divergent regions

To understand how well conserved wild strain gene copy numbers in HDRs are with N2 reference genes, we quantified the proportion of orthogroups that contained gene copy-number variants (CNVs), presences/absence variants (PAVs, non-reference genes), or are in stable copy (*n*-to-*n*) with N2 reference genes inside and outside of HDRs for each wild strain. When the N2 reference strain and a wild strain have a different number of orthologs that contribute to an orthogroup, those orthogroups were defined as CNVs. PAVs were defined when a wild strain has an ortholog that contributes to an orthogroup but the N2 reference strain does not, and stable-copy orthogroups were defined when the N2 reference strain and a wild strain have the same number of orthologs that contribute to an orthogroup. We found that gene CNV and PAV orthogroups were statistically enriched in HDRs compared to outside of HDRs, and stable-copy orthogroups and single-copy orthologs (conserved among all 142 strains) were enriched outside of HDRs (Supplementary Figure 22). When when we considered GPCRs, F-box proteins, C-type lectins, cytochrome P450s, or nuclear hormone receptors (see *Methods*, Supplementary Table 11), we found similar trends of CNV and PAV orthogroups enriched in HDRs and stable-copy orthogroups enriched outside of HDRs for most of these gene classes (Figure 5). However, we found cytochrome P450s were enriched for PAV orthogroups outside of HDRs, and HDRs were enriched for stable-copy orthogroups. Additionally, NHRs appear to be well conserved with the N2 reference strain among wild strains, where nearly 90% of NHR orthogroups outside of HDRs are in stable-copy with the N2 reference strain, and nearly 75% of NHR orthogroups in HDRs are in stable-copy with the N2 reference strain (Figure 5). These data reveal that not only do NHRs and cytochrome P450s contribute to the core pangenome and are enriched in HDRs, but many of these genes are in stable-copy with the N2 reference strain. Alternatively, we find that F-box orthogroups are not well conserved with the N2 reference strain, where nearly 75% of F-box orthogroups in HDRs are PAVs and nearly 50% of F-box orthogroups outside of HDRs are also PAVs (Figure 5). F-box gene families are much larger in *C. elegans* in comparison to other animal species^75,76^, and they have also been shown to be variable in gene family size between *Caenorhabditis* species^77^. Because this analysis method fails to identify PAVs that are present in the N2 reference strain and absent in a wild strain, we classified N2 reference genes as inside or outside of HDRs for each wild strain, and performed the same enrichment analysis explained above for N2 gene CNVs, PAVs, and stable-copy orthogroups for each wild strain. This analysis revealed similar results of CNV and PAV orthogroups enriched in HDRs and stable-copy and single-copy orthogroups enriched outside of HDRs (Supplementary Figure 23). Although the same gene classes are enriched in HDRs among wild strains and in the N2 reference strain, both wild strain HDRs in relation to the N2 reference strain, and N2 reference genes in HDRs among all wild strains are enriched for gene CNVs and PAVs, indicating that these polymorphic regions contain immense gene content diversity in environmental sensing and response genes that might be niche-specific and could help these nematodes survive in different environments.

**Figure 5.**
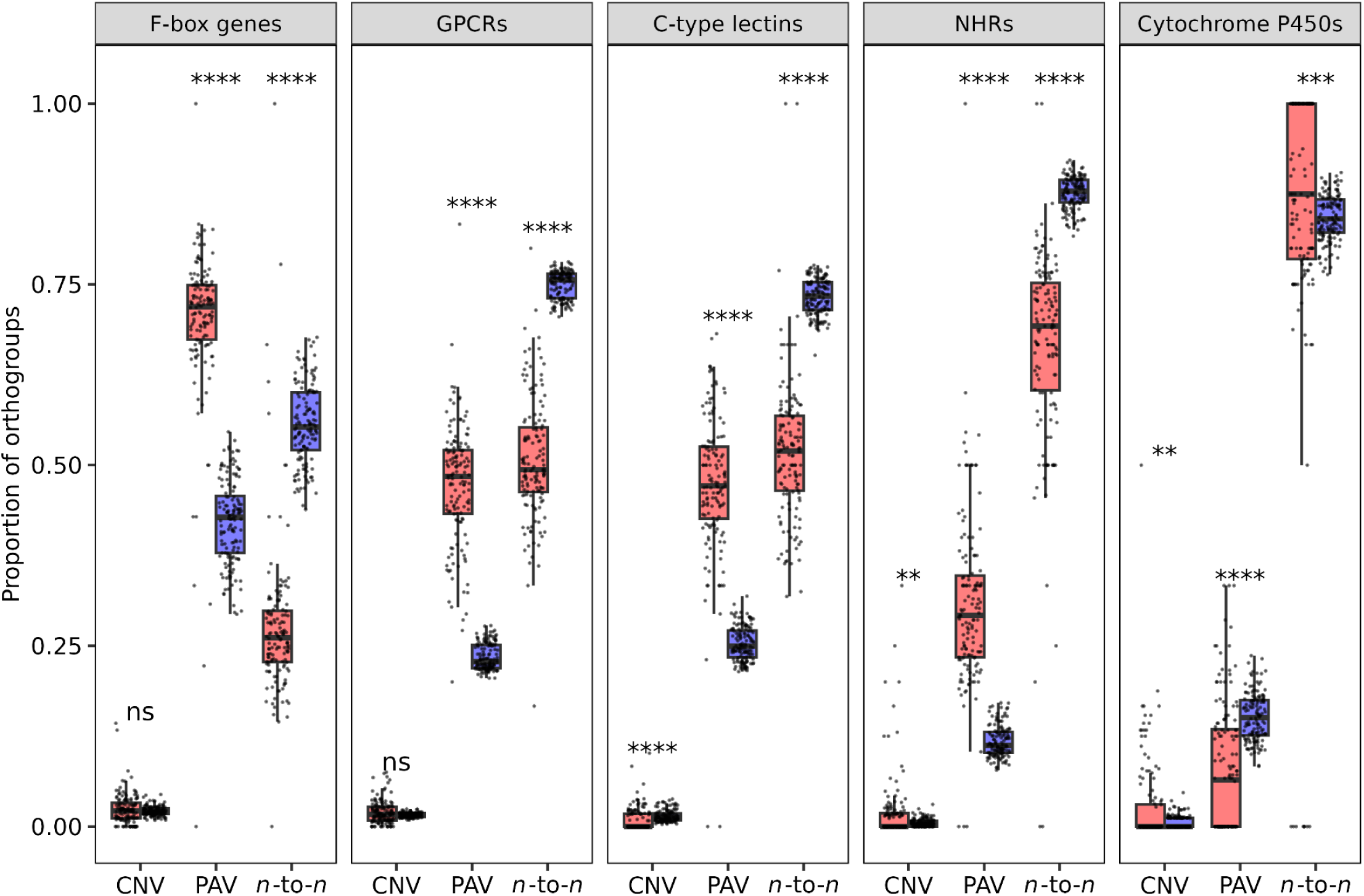
Wild strain HDR environmental sensing and response genes are found at variable copy-number in relation to the reference genome. Enrichment of gene CNV, PAV, or stable-copy (*n*-to-*n*) orthogroups inside (red) or outside (blue) of HDRs where each point represents a wild strain. Gene classes (facets) are defined by IPR annotations (see *Methods*, Supplementary Table 11). Box and whisker plots display the median value for each metric on the x-axis. The upper and lower whiskers correspond to the first and third quartiles, respectively. \*\**p* < 0.01, \*\*\**p* < 0.001, \*\*\*\**p* < 0.0001, ns = not significant.

Although stable-copy cytochrome P450s are enriched in HDRs, the number of cytochrome P450 orthogroups is the fewest out of the enriched gene classes in HDRs (Supplementary Figure 24). To mitigate estimates of enrichment based on total number of orthogroups inside or outside of HDRs and because of the large proportion of orthogroups in HDRs that are in stable copy with the N2 reference strain, we looked at the proportion of single-copy orthogroups that are found in HDRs for the same five genes classes and found that cytochrome P450s have the largest proportion of single-copy orthogroups that are found in HDRs among these gene classes (10.7% on average) (Supplementary Figure 25). Single-copy orthologs in HDRs are likely under different selection regimes than those outside of HDRs, causing differences in sequence conservation. To quantify allelic diversity among single-copy orthologs inside versus outside of HDRs, we calculated synonymous divergence (K_s_) and non-synonymous divergence (K_a_) among all single-copy orthologs. Of the 11,562 single-copy orthogroups used for divergence calculations (see *Methods*), 82 were found in HDRs for at least 71 wild strains (50% of strains). We found that K_s_ and K_a_ were elevated among single-copy orthologs found in HDRs compared to single-copy orthologs outside HDRs, and K_a_ / K_s_ was the same (Supplementary Figure 26). These data are consistent with the observed elevated nucleotide diversity in HDRs and reveal that conserved single-copy genes in these regions might have diverged longer ago than non-HDR single-copy orthologs, but they do not appear to be under different selective constraints.

### Hyper-divergent region gene copy-number differences are associated with abamectin resistance

The macrocyclic lactone (ML) class of anthelmintics are drugs used to treat parasitic nematode infections of humans and livestock^78^. Frequent and widespread use has caused resistance to all known ML drugs across nearly all parasitic nematode species^78,79^. Genetic selection experiments using *C. elegans* identified the heteromeric glutamate-gated chloride (GluCl) channels as targets of ML drugs with laboratory-derived mutations in *avr-14*, *avr-15*, and *glc-1*^80,81^. This drug class is derived from natural products produced by the bacteria *Streptomyces avermitilis* found worldwide in soil environments^82^. *C. elegans* wild strains live adjacent to soil environments and also harbor natural ML resistance^83,84^. This resistance has been mapped to a QTL containing *glc-1*^84–86^. Because GLC-1 is a part of the GluCl channel, the assumption was that variation in this gene caused ML resistance, but no specific variants or genes were validated. A recent study demonstrated that loss of *glc-1* alone is not sufficient to cause strong ML resistance^87^, and across different nematode species, the QTL for ML resistance has been broad, likely containing multiple causal genes^84–86,88^. We found that the ML response QTL (abamectin) overlapped with an HDR, making the gene content underlying this drug response variation poorly defined (Supplementary Figure 27, Supplementary Tables 12 and 14). We leveraged the *C. elegans* pangenome to elucidate the gene content in this QTL containing *glc-1* and revealed distinct haplotypes among strains that were susceptible or resistant to abamectin (Figure 6, Supplementary Figure 28). Strains resistant to abamectin have deletions of four genes (with the exception of the ECA36 strain) upstream of *glc-1*, and all susceptible strains have these four genes (Figure 6). When we looked downstream of *glc-1*, we found no other CNVs or PAVs that were specific to resistant strains (Supplementary Figure 29). These four gene deletions upstream of *glc-1* include *F11A5.18*, *irld-26*, *oac-15*, and *F11A5.16*, and we found no orthologs of these genes in the rest of the genome for any resistant strain (aside from strain ECA36). Although the resistant strain ECA36 does not harbor these gene deletions, the amino acid divergence of these genes is substantial in relation to N2 reference genes, ranging from 52.8% (F11A5.18) to 94.5% (OAC-15) identity (Figure 6). The protein IRLD-26 in the resistant strain ECA36 has 86.5% identity with the susceptible N2 reference strain and is predicted to be involved in insulin / EGF signaling. This gene might have indirect effects on abamectin resistance by modulating drug metabolism or cellular uptake^89,90^. We also identified 109 SNVs that are shared among all resistant strains and absent in susceptible strains in the region of interest (Supplementary Figure 30, Figure 6). Of these SNVs, 68 were found in coding regions, and 23 were found in *F11A5.15* and 14 each in *glc-1* and *F11A5.5* (Supplementary Figure 30, Supplementary Table 15). To understand how these variants might affect function of GLC-1 and F11A5.5, which are present in all resistant strains, we looked at the predicted functional consequences of codon or splice site region changes using BCFtools CSQ^91^, and the predicted deleteriousness of missense variants using BLOSUM62^92^ and Grantham^93^. We classified missense variants as “high impact” if they had a BLOSUM62 score less than zero or a Grantham score greater than 100. Notably, GLC-1 had a high impact missense variant (T163M) located in the ligand-binding domain, and a missense variant (V367A) in the transmembrane domain, which might alter abamectin binding efficacy (Supplementary Table 15)^94,95^. For F11A5.5, which is predicted to be involved in macromolecule glycosylation at the cell membrane, four splice region variants and two high-impact missense variants located in the glycosyl transferase, family 11 domain are shared among resistant strains (Supplementary Table 15). Although the four genes in the ECA369 strain that are absent in the resistant strains, aside from ECA36, have low percent identity with N2 reference genes, the missense variants in the ECA369 strain are different than the ECA36 strain for *F11A5.18*, *irld-26*, and *oac-15*. ECA36 has two, 12, and six high impact missense variants in F11A5.18, IRLD-26, and OAC-15, respectively, and these high impact variants in IRLD-26 and OAC-15 are in receptor or enzymatic domains, which might alter enzymatic function and have larger modulatory roles in abamectin susceptibility (Supplementary Table 15). Even though the ECA36 strain has F11A5.18, IRLD-26, OAC-15, which are absent in other resistant strains, the genetic divergence of these proteins is extreme in relation to the susceptible N2 reference strain. We found no SVs that were shared among all abamectin resistant strains, and we did not find a difference in *glc-1* or *F11A5.5* RNA fold-coverage levels between susceptible and resistant strains (Supplementary Figure 31)^66^. Additionally, we looked to see if resistant strains shared any variants in *avr-14* and *avr-15* that might cause partial or full loss of function because a previous study showed that deletions of *avr-14* and *avr-15* were sufficient to cause abamectin resistance^87^. We found no variants in *avr-14* or *avr-15* that were shared among resistant strains (Supplementary Table 15). The mechanism of resistance might be a combination of altered GLC-1 function with the partial or full loss of function of metabolic proteins such as F11A5.5, IRLD-26, and OAC-15.

**Figure 6.**
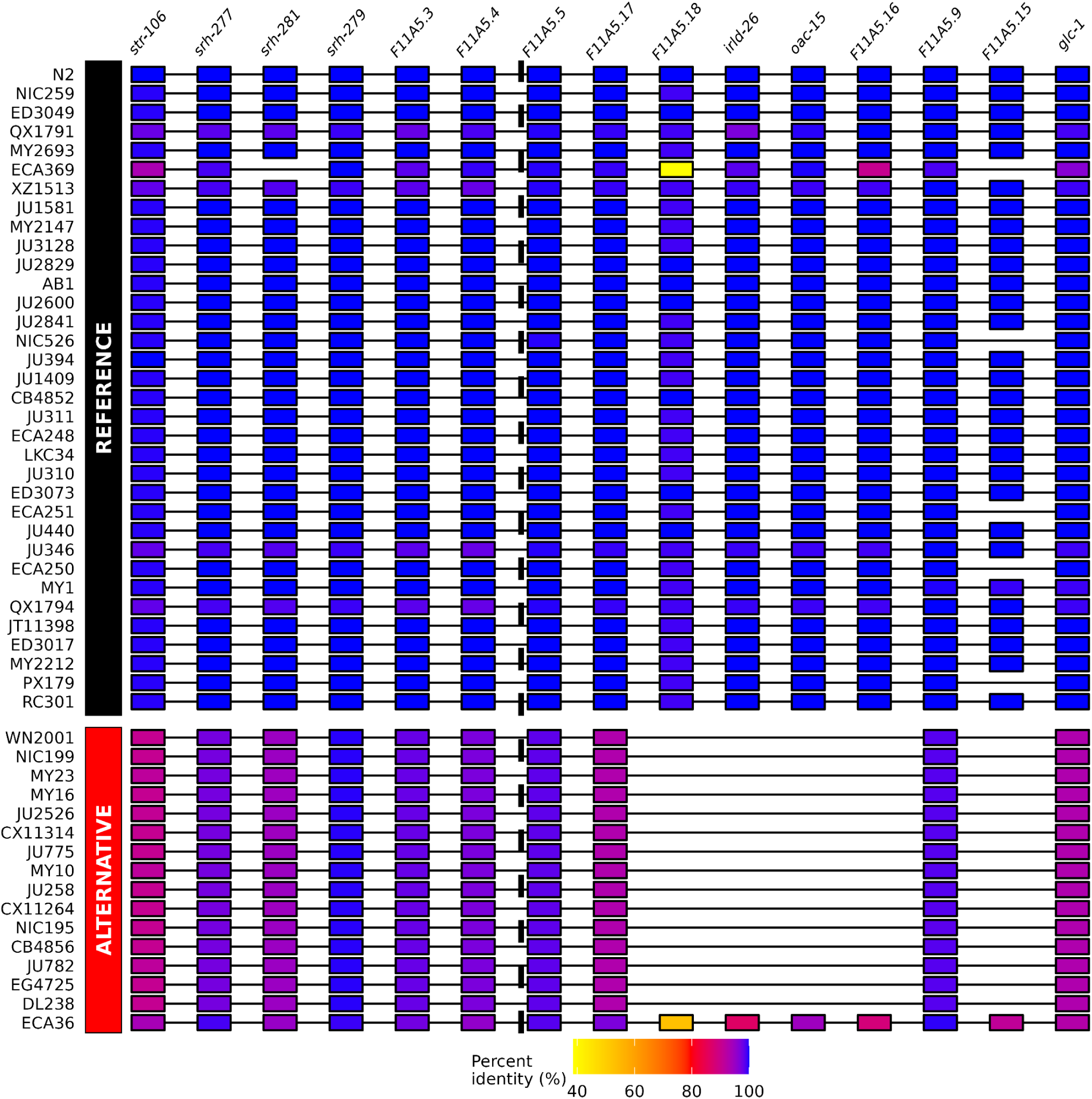
Haplotype associated with abamectin resistance has multi-gene deletions. Strains are faceted by their genotype for the peak SNV (vertical dashed line) associated with abamectin resistance (Supplementary Figures 27 and 28). Orthologous genes are indicated by gene position and colored by percent amino acid identity to the N2 reference ortholog.

Recently, large effect QTL associated with resistance to MLs have been identified in the parasitic nematode of small rudiments *Haemonchus contortus*^79,88,96^. The QTL for MLs in *H. contortus* are also on chromosome V. The chromosomal gene content between *C. elegans* and *H. contortus* is highly conserved^97^, and orthologs shared in the QTL of both species could suggest a conserved genetic mechanism of resistance. A previous study identified one-to-one orthologs across different *C. elegans* ML QTL with *H. contortus* ML QTL and did not find shared orthology of genes implicated in ML resistance^85^. We performed a similar analysis to look for orthologs of the 21 *C. elegans* genes surrounding our peak marker in the recently narrowed ML QTL in *H. contortus* (V:37.25-37.55 Mb) and a broader locus surrounding this QTL (V:36-45 Mb), which is genetically differentiated after sub-therapeutic treatment with MLs^79,88^. Three of the 21 N2 reference genes had orthologs outside of the *H. contortus* QTL, and *F11A5.5* and *F11A5.15* were the only genes that had orthologs in the broader *H. contortus* QTL (Supplementary Table 16). *F11A5.5* had three orthologs in *H. contortus* and *F11A5.15* had two. For each of these orthogroups, one of the orthologs in *H. contortus* is contained in the broader locus associated with ML resistance. These data suggest that variation in *F11A5.5* and/or *F11A5.15* might be a shared mechanism of resistance in *C. elegans* and *H. contortus*. The shared genetic variation in *F11A5.5*, most proximal to the peak marker associated with abamectin resistance, *glc-1*, and the loss or diverged sequences of *irld-26* and *oac-15* among resistant strains might all contribute to abamectin resistance in *C. elegans*.

## Discussion

The *C. elegans* reference genome has undergone multiple updates with extensive manual curation of gene annotations since its initial assembly in 1998^57,62,98,99^. Despite decades of curation, comparative genomics revealed punctuated regions of the *C. elegans* N2 reference genome that are extremely polymorphic in the population (HDRs)^42^. These HDRs are found in numerous organisms across the tree of life, and their genomic composition in non-reference individuals has not been characterized because of the need for *de novo* genome assemblies of non-reference individuals. In recent years, advances in massively parallel sequencing, specifically long-read sequencing, and *de novo* assembly algorithms have made reference genome construction more straightforward. In this study, we created reference genomes and gene annotations for 140 strains and constructed the first *C. elegans* pangenome to define HDR gene content and quantify structural and gene content variation.

### The functional role of structural variation

*Caenorhabditis* species show high rates of intrachromosomal rearrangements and extreme intraspecies structural variation^59,60^, and the functional role of this structural variation has yet to be realized. The geographic origin of the *C. elegans* species is believed to be the Hawaiian islands or somewhere in the Pacific region because of the elevated genetic diversity and spatial patterns of relatedness^42,67^. Consistent with strains collected from the Hawaiian islands displaying elevated nucleotide diversity, we found that they also display much more structural variation than non-Hawaiian strains. The sharing of SVs among Hawaiian strains might be reflective of common origins or adaptive evolution, where specific SVs provide a fitness advantage in certain ecological niches. Alternatively, these SVs could be the result of genetic drift or relaxed selection, and might not have a pertinent, functional role. If adaptive, these SVs could arise from negative frequency-dependent selection or condition-specific selection, which might explain their rare-to-intermediate frequency in the population and elevated nucleotide diversity at loci that harbor SVs^42,100,101^. For example, we identified a large inversion (438 kb) that is shared among three strains collected years apart from different geographic locations on the island of Kaua’i, and a 4.03 Mb inversion in the wild strain ECA3088 from O’ahu (Supplementary Figures 32 and 33). These SVs might be lineage-specific and maintained by condition-specific selection. This catalog of SVs can be further explored and used for SV-based genome-wide association studies to understand how structural differences might influence adaptive traits, which has been validated in many plant species including maize^102^, *Arabidopsis*^20^, soybeans^103^, roses^104^, buckwheat^17^, and rice^105,106^.

### The adaptive potential of hyper-divergent region gene content diversity

This pangenome has revealed the extreme extent of gene content variation among wild strains, where only 14,998 of the 19,984 N2 reference genes are conserved among all wild strains in the pangenome strain set. The selected 142 strains used for pangenome construction appear to capture genes that are core to the species, and these strains reveal the immense gene diversity among wild strains, with each strain possessing their own repertoire of private genes (Figure 3b, Supplementary Figure 11). The gene content of HDRs was defined among all wild strains, which revealed the primary contribution of HDR genes to the accessory pangenome. Although wild strain HDRs were enriched for the same classes of environmental sensing and response genes as the N2 reference genome^42^, we found substantial gene CNVs and PAVs in wild strain HDRs. The diversification of these gene classes in wild strains might reflect condition-specific selection, caused by local environmental conditions that favor certain receptor, immune, or regulatory genes. F-box proteins are largely expanded in *Caenorhabditis* species in comparison to other animals^75,76^, and these large families have been observed in tandem segments with rapid diversification^77,107,108^, which could lead to novel functions that might be selected in specific environments. Surprisingly, NHRs and cytochrome P450s that contribute to the core pangenome were also enriched in wild strain HDRs. Furthermore, approximately 10% of NHR and cytochrome P450 single-copy orthologs that are conserved across all 142 strains are found in HDRs (Supplementary Figure 25). We found that single-copy orthologs in HDRs have elevated K_s_ and K_a_ in comparison to single-copy orthologs found outside HDRs (Supplementary Figure 26). HDRs appear to have experienced multiple forms of selection, where large gene classes are enriched for CNVs and PAVs (F-box genes, GPCRs, C-type lectins) that diversify the number of environmental and immune recognition genes, and conserved single-copy genes have undergone allelic diversification among wild strains while conserving copy number (NHRs and cytochrome P450s). The observed elevated K_s_ and K_a_ in HDR single-copy orthologs is not likely explained by increased mutational load in HDRs because a previous study found mutations to be uniformly distributed across *C. elegans* chromosomes^109^. Rather, the allelic diversification might be condition-specific and reflect selection for specific receptor configurations in immune and xenobiotic stress responses genes for defense in different environments.

### Sources of strain-specific genes in C. elegans

The private pangenome of *C. elegans* is quite large, where each wild strain has, on average, 121 genes that are unique. The pangenome strain set was specifically selected to maximize the capture of genetic variation, and if we add strains that are closely related to strains in the current pangenome strain set, many lineage-specific private orthogroups might become accessory and the discovery rate of new orthogroups would decrease, leading to the saturation of orthogroup rarefaction curves (Figure 3b, Supplementary Figure 12). The observed private genes might stem from duplication divergence, which appears to be the case for F-box genes and has been shown in other studies in *Caenorhabditis*^77^ (Supplementary Figures 20 and 21). Other sources include horizontal gene transfer^110,111^, as well as ancient introgression prior to the transition to androdioecy. It would be important to validate these private genes using expression support from RNA sequencing. Lastly, an unavoidable source of private genes come from gene model error. On average, each wild strain has 4,457 non-N2 reference genes, and the N2 reference strain has, on average, 1,797 genes that are not found in a wild strain (Supplementary Figures 34 and 35). The majority of these genes contribute to the accessory pangenome, likely indicative of some consistent gene model error among wild strains, such as pseudogenes that have been manually removed from N2 gene annotations but are predicted in wild strains. However, it is not feasible to manually curate over three million genes, and it is difficult to systematically differentiate non-N2 reference genes as gene model errors or true gene content diversity without manual curation. Accurate *ab initio* gene model prediction has historically been quite difficult, with substantial efforts for its improvement in recent years^112–114^. To our benefit, *C. elegans* has one of the most curated gene annotations of any organism, which enables us to quantitatively benchmark gene model prediction software that cannot be done in other organisms.

### Pangenomes to understand the genetics of quantitative traits

One of the greatest strengths of pangenomes is the resolution of gene content in polymorphic genomic regions. Pangenomics has expanded canonical SNV-based GWAS to implement larger sources of genetic variation such as structural and gene content variation to understand the genetic basis of quantitative traits. GWAS have been powerful to identify genomic variants that confer quantitative traits in *C. elegans*^115–117^. However, sometimes SNVs that are found to be strongly associated with a trait are in HDRs and the genomic content of the QTL is unknown. Because these SNVs are used as markers to identify QTL, they are not always causal in the quantitative trait difference, and other sources of genetic variation such as SVs or CNVs might be causal but cannot be reliably identified using a single reference genome. Using pangenomics, we defined the HDR genes in a previously identified QTL associated with resistance to the anthelmintic drug abamectin and found multi-gene deletions in wild strains correlated with resistance. A target of abamectin, GLC-1, contained three missense variants that were shared among all resistant strains (Supplementary Table 15). A previous study showed that the loss of *glc-1* is not sufficient for abamectin resistance^87^, and QTL identified across nematode species have been broad, suggesting that multiple genes might play a role in the mechanism of resistance^84–86,88^. We hypothesize the mechanism of abamectin resistance in *C. elegans* is a combination of altered GLC-1 functional in conjunction with altered or loss of function of upstream genes, such as *F11A5.5*, *irld-26*, and *oac-15*, which are either lost or extremely diverged in resistant strains. These proteins are involved in broad functions such as macromolecule glycosylation, acylation, and insulin / EGF binding where they might have accessory functions in abamectin metabolism. However, functional validation of this locus in abamectin resistance is necessary. Identifying a genetic mechanism of resistance to ML anthelmintic drugs will provide potential insights into the genetic mechanisms of resistance in parasitic nematodes that are similar to *C. elegans*, such as hookworms, which cause huge global health and economic issues. We believe this *C. elegans* pangenome will aid the *Caenorhabditis* community in unraveling long-standing genotype-phenotype relationships of quantitative traits.

## Methods

### Strain maintenance

All *C. elegans* strains used in this study were maintained at 20°C on a modified nematode growth medium (NGMA) (consisting of 1% agar and 0.7% agarose)^118^ on 6 cm plates using *Escherichia coli* (strain OP50) as the food source.

### Long-read DNA sequencing

Nematodes were grown on four 10 cm NGMA plates seeded with OP50. When the plates were close to starvation, the animals were washed off using cold (4°C) M9 buffer (3 g of KH2PO4, 6 g of Na2HPO4, and 5 g of NaCl in 1 L Milli-Q water) into a 50 mL conical tube and centrifuged at 2,000 rpm for eight minutes with a break of three for a slow stop of the centrifuge. After this and subsequent spins, samples were settled on ice for two minutes before the supernatant was aspirated. The pellet was washed three times using cold M9 buffer + 0.01% Tween 20. A final wash was performed using cold M9 buffer. Most of the supernatant was removed, and the pellet was transferred to a 1.7 mL microfuge tube using a glass pasteur pipette. If smaller pellets were needed, the resuspended material was divided into multiple tubes. The animals were centrifuged at 2,500 rpm for five minutes. Supernatant was removed, and the pellets were weighed. The pellet was flash frozen in liquid nitrogen and stored at −80°C.

For PacBio HiFi sequencing of 138 wild *C. elegans* strains, 60 mg of animal pellet was used to make High Molecular Weight (HMW) DNA using the Monarch HMW DNA Extraction Kit for Tissue (New England Biolabs, Catalog #T3060L). The following modifications were made to the protocol. Before using the pestle, 50 µL of the lysis buffer was added to the sample. The remaining 600 µL was added after homogenization. For the lysis at 56°C after homogenization, we incubated in a Thermomixer for 15 minutes at 750 rpm and then at 550 rpm for 30 minutes or longer. We used cold (4°C) Protein Separation Solution. After the inversion step, samples were incubated on ice for 10 minutes. We used three Capture beads. For elution, we added 200 µL Elution Buffer, incubated for five minutes at 56°C at 300 rpm. We then added another 50 µL of the Elution Buffer before separating the DNA from the beads. The final solubilization was done at 37°C for 30 minutes at 300 rpm. The final samples were sent to Maryland Genomics at the University of Maryland for library generation and sequencing. ONT sequencing for strains JU3226 and ECA1493 was received from Drs. Ryan Pellow and Ofer Rog’s group at University of Utah. PacBio HiFi sequencing for strain ECA741 was received from Drs. Stefan Zdraljevic and Leonid Kruglyak at the University of California, Los Angeles.

### Genome assembly and quality control

Sequenced PacBio HiFi genomic reads were demultiplexed and adapters were removed by the University of Maryland sequencing core using *lima* v2.12.0 (https://lima.how). PCR duplicates were removed using *pbmarkdup* v1.1.1 (https://github.com/PacificBiosciences/pbmarkdup). Primary *de novo* assemblies were generated using *hifiasm* v0.24.0-r702^53^ with the initial bloom filter *-f39*, which is recommended to save memory on k-mer counting for large amounts of sequencing data, haplotig purging disabled (*-l0*, recommended for homozygous genomes). If ONT reads were available for a strain, primary *de novo* assemblies were generated using HiFi and ONT reads using *hifiasm* with the same parameters as HiFi-exclusive assemblies but ONT reads were provided to the *-ul* argument. After assembly, contigs were taxonomically classified using *DIAMOND* 2.1.13^119^ with parameters *--faster* and an e-value of 1e-10 against the UniProt database (Release 2025_03)^120^. Contigs that were annotated as non-*Nematoda* were purged using *BlobToolKit* v4.4.5^121^ with *--param bestsumorder_phylum--Inv=Nematoda*. We only used *do novo* genome assemblies with an N50 of one megabase or larger for analyses. *pbmarkdup*, *hifiasm*, and *BlobToolKit* executions were carried out using the Nextflow pipeline available at https://github.com/AndersenLab/assembly-nf.

If PacBio HiFi sequencing for a strain had been collected from multiple runs, the sequencing data was pooled together and genome quality metrics of N50, L90, and genome BUSCO score were used to determine if individual HiFi sequencing or pooled HiFi sequencing yielded a higher-quality genome assembly (Supplementary Table 1). For strain JU3226, *de novo* assembly using ONT reads was performed using *hifiasm* with parameters *-l0* and reads were supplied to parameter *--ont*. Contigs sourced from non-*Nematoda* DNA were removed using the same methods previously described. For strain ECA2555, DNA preparation was contaminated with *C. nigoni*, resulting in an assembly with six large contigs (> 10 Mb) and thousands of small contigs (<100 kb). *C. nigoni* contigs were removed from the primary assembly by removing contigs with a mean HiFi fold-coverage of less than 10x, resulting in a highly contiguous assembly. The updated *C. elegans* N2 reference genome assembly, CGC1, was retrieved from NCBI PRJDB19205.

### Structural variant identification

SVs among 140 wild strains and CGC1 were identified in relation to the N2 (WormBase, WS283) reference genome using Phased Assembly Variant caller (PAV) v2.4.6^61^ with parameters *--latency-wait 900* and *--nt* supplied to the Docker container. SVs were filtered for those that passed all filters and were larger than or equal to 50 bps using *bcftools* v1.20^122^. Structural variants identified among all 141 strains were then merged using *jasmine*^123^ v1.1.5 with parameter *--output_genotypes*. Principal component analysis of SVs was performed using the *prcomp* function from the *stats* package in base R v4.2.1 with parameters of *center* and *scale. = TRUE)*. SVs were filtered for a MAF of 0.05 prior to PCA analysis.

### Protein-coding gene annotation

Repetitive sequences were masked in *de novo* genome assemblies using *RepeatMasker* v4.2.1^124^ and a custom *C. elegans* repeat library created using methods described in a previous study^125^. *BRAKER3* v3.0.8^68^ was used to predict protein-coding gene models on soft-masked genomes using the N2 (WormBase, WS283) proteome to guide gene annotation. Proteomes of the longest-isoform for gene annotations were created using *agat_sp_keep_longest_isoform.pl* from AGAT v1.4.1^126^ and *gffread* v0.12.7^127^. Gene model biological completeness of each gene annotation was estimated using protein *BUSCO* v5.8.3^58^ with the *nematoda_odb10* database (Supplementary Table 1). *RepeatMasker*, *BRAKER3*, *agat_sp_keep_longest_isoform.pl, gffread*, and *BUSCO* executions were carried out using a Nextflow pipeline available at https://github.com/AndersenLab/geneAnno-nf.

### Pangenome gene set analysis

To identify orthogroups among the 142 *C. elegans* pangenome strain set we used *OrthoFinder* v3.1.0^69^. *OrthoFinder* identifies orthogroups among a set of 64 “core” proteomes, and then “additional” proteomes are used to update “core” orthogroups as well create new orthogroups that are not present in the 64 “core” proteomes. *OrthoFinder* authors suggest selecting proteomes that capture a broad range of evolutionary diversity for the 64 “core” set, so we selected strains by maximizing the phylogenetic spread across the *C. elegans* species tree. To do this, we calculated pairwise patristic distance from the species tree and used Partitioning Around Medoids (PAM) clustering to select 64 medoids (strains) that minimize the total patristic distance to all other cluster members. The 64 medoid strains serve as representatives that collectively maximize phylogenetic spread. Default parameters of *OrthoFinder* were used for “core” orthogroup creation, and then the additional 78 proteomes were provided using *--assign* to the “core” orthogroups results directory using *--core*. The *Orthogroups.tsv* and *Orthogroups_UnassignedGenes.tsv* were concatenated together and orthogroups were classified as core (found in all strains), accessory (found in some but not all strains), or private (found in a single strain) based on the number of strains that had contributing orthologs (Figure 3). Mitochondrial genes were removed from gene set analyses (orthogroups OG0042586, OG0042630, OG0043448, OG0044504, OG0044948, OG0045081, OG0045344, OG0046458, OG0047997, OG0049864, OG0053290, and OG0054049). To cluster orthogroups into a presence/absence supermatrix, a maximum-likelihood tree was created from protein BUSCOs using *mafft* v7.526^128^, *trimal* v1.5.1^129^, and *iqtree* v3.0.1^130^ using the GTR20+G4 model using 1000 bootstraps (Supplementary Figure 13). All software executions to build the BUSCO phylogenetic tree were carried out using a Nextflow pipeline available at https://github.com/lstevens17/busco2phylo-nf.

### Hyper-divergent region lift-over

To lift over HDRs from N2 reference genome coordinates to wild strain genome coordinates we used *NUCmer* v3.1^131^ to perform whole genome-genome alignment of every wild strain to the reference N2 genome (WormBase, WS283). To lift over hyper-divergent region coordinates to wild strain genome coordinates, we first filtered wild strain genome alignments for those that overlapped with HDRs for each wild strain. We then filtered alignments for the wild strain contig that spanned the N2 reference genome hyper-divergent region the most to ensure the removal of multi-alignments from other contigs. To remove distal intra-contig alignments, we removed alignments that were >75 kb distance away from the most proximal upstream and downstream alignments. Lastly, when wild strain contig alignments did not span one or both boundaries of an HDR, we extended the region to the most proximal contig alignment (up to 50 kb distance) outside of the HDR boundary or used the end of the alignment contig alignment span when no proximal alignment was found outside of the HDR boundary (Supplementary Figure 16). HDRs that spanned less than 5 kb were removed, and HDRs that were 5 kb distance or less apart were merged.

### Functional enrichment analysis

*InterProScan* (IPR) v5.75^74^ was used to identify functional protein domains and their associated Gene Ontology IDs that are enriched in wild strain HDRs. IPR was run using options *--goterms*, *--iprlookup*, and *--disable-precalc* on the longest-isoform proteome of all 142 strains of *C. elegans*. IPR annotations were filtered to only unique IPR terms for each protein. A total of 2,360,448 / 3,173,110 (74.39%) proteins were annotated with IPR functional protein domains in the wild strain pangenome (Supplementary Table 8). The 2,360,448 annotated proteins served as the background set and proteins found in wild strain HDRs were assessed for enrichment categorized by pangenome gene set. A total of 57,444 / 70,320 (81.69%) proteins in wild strain HDRs that contribute to the core pangenome gene set received IPR annotations, 99,973 / 133,445 (74.92%) in the accessory pangenome gene set, and 1,774 / 2,459 (72.14%) proteins in the private pangenome gene set. For IPR domain enrichment analysis, a one-sided hypergeometric test was conducted in R, with a false discovery rate adjusted p-value using the R package *stats* v4.2.1. IPR functional protein domains were classified as significantly enriched if they had an adjusted p-value less than 0.05. Fold enrichment is defined as: (Total number of wild strain HDR proteins with IPR annotation *x* / Total number of proteins in pangenome with IPR annotation *x*) / (Total number of wild strain HDR proteins with an IPR annotation in pangenome gene set / Total number of proteins in pangenome with an IPR annotation) (Supplementary Table 9). The IPR description of ‘Repeat of unknown function XGLTT” was removed from the enrichment analysis because proteins with this annotation appear to stem from gene model error.

### Enrichment of gene copy-number variation in HDRs

After HDR coordinate lift-over to wild strains, all wild strain genes were categorized as inside or outside of HDRs and the orthogroups that they contribute to. Orthogroups were then classified as CNV, PAV, or stable-copy (*n*-to-*n*) for all identified orthogroups between N2 and a wild strain. Orthogroup counts for CNVs, PAVs, and stable copy were normalized by the total number of orthogroups found in HDRs or outside HDRs for each wild strain (Figure X). To identify statistical enrichment of each orthogroup class inside or outside of HDRs, we used a paired Wilcoxon signed-rank test using the R package *rstatix* v0.7.2 with Benjamini-Hochberg HDR-adjusted correction. Wild strain genes with IPR annotations containing the string “F-box” were classified as F-box proteins, proteins with IPR annotations containing “7TM” were classified as GPCRs, proteins with IPR annotations containing “C-type lectin” were classified as C-type lectins, proteins with IPR annotations containing “Cytochrome P450” were classified as cytochrome P450s, and proteins containing “uclear hormone receptor” were classified as nuclear hormone receptors (Supplementary Table 11, Figure 5). “Domain of unknown function DUF38/FTH, *Caenorhabditis* species” likely represent F-box genes, since over half of the predicted N2 F-box genes also have this IPR annotation^75,132^. However, we chose to be conservative in how we classified F-box proteins and did not include the IPR annotation of “Domain of unknown function DUF38/FTH, *Caenorhabditis* species” as an F-box protein classifier (Supplementary Table 11). Because this analysis method fails to identify PAVs that are present in N2 and missing in a wild strain, we classified N2 reference genes as inside or outside of HDRs for each wild strain, and performed the same enrichment analysis explained above for N2 CNVs, PAVs, and stable-copy orthogroups for each wild strain (Supplementary Figure 23).

### K_A_ / K_S_ estimates of single-copy orthologs

Single-copy orthologs shared among all 142 *C. elegans* strains were identified using *OrthoFinder*^69^ (see *Pangenome gene set analysis*). Protein sequence alignments for each single-copy orthogroup was reverse-translated into codon-aware coding sequence alignments using *PAL2NAL* v14^133^. Synonymous and non-synonymous site divergence was calculated using the *kaks()* function from *seqinr* R package v4.2.44^134^ with parameter *rmgap = TRUE* for single-copy orthogroups that passed the criteria of possessing at least 127 orthologs (90% orthogroup occupancy) in the multiple coding sequence alignment after filtering orthologs that had 70 percent codon occupancy or less and had less than 100 codons in the coding sequence alignment. After filtering, 11,562 / 12,315 (93.9%) of single-copy orthologs were retained. Single-copy orthogroups were classified as HDR or non-HDR based on the criterion of orthologs found in HDRs for at least 71 wild strains (50% of strains). To identify statistical differences in synonymous and non-synonymous rates in HDR single-copy orthologs versus non-HDR single-copy orthologs, we used the Wilcoxon rank-sum statistical analysis using the R package *rstatix* v0.7.2 with Benjamini-Hochberg FDR-adjusted correction.

### Genome-wide association mapping of abamectin resistance

Using a previously described high-throughput fitness assay^135,136^, 210 *C. elegans* wild strains were phenotyped in both abamectin (7.5 nM) and dimethyl sulfoxide (DMSO) (control) conditions (Supplementary Tables 12, 13, and 14). Briefly, strains were passaged for four generations on NGMA plates, bleach-synchronized, and 25-50 embryos were aliquoted into 96-well microtiter plates at a final volume of 50 μL of K medium. After 12 hours, arrested L1 larvae were fed *Escherichia coli* HB101 bacterial lysate (Pennsylvania State University Shared Fermentation Facility, State College, PA) at a final concentration of 5 mg/mL in K medium. Animals were grown for 48 hours at 20°C while shaking at 180 rpm. Three L4 larvae were sorted into a 96-well microtiter plate with 10 mg/mL of HB101 bacterial lysate, 50 μM kanamycin, and either 1% DMSO or abamectin dissolved in 1% DMSO using a large-particle flow cytometer (COPAS BIOSORT, Union Biometrica; Holliston, MA). Animals grew for 96 hours at 20°C while shaking at 180 rpm. The animals and their offspring were treated with 50 mM sodium azide in M9 Buffer to paralyze and straighten nematodes to obtain accurate length measurements with the COPAS BIOSORT. Measurements collected in the high-throughput fitness assay were processed with the *R/easysorter* v 4.0.3 package^136^. Analyses were performed on a strain-specific basis. A genome-wide association (GWAS) mapping was performed using the differences in responses of strains exposed to abamectin and DMSO (control) conditions. The regressed animal length (q90.TOF) trait data were analyzed using the *NemaScan* mapping pipeline (https://github.com/AndersenLab/nemascan-nf)^137^ with the default parameters. The association mapping was conducted using biallelic SNVs from the *C. elegans* hard-filtered variant call format file on the *Caenorhabditis* Natural Diversity Resource (CaeNDR) (Release: 20250625)^41^. Significance was determined using a Bonferroni threshold for all tested markers.

### Wild strain RNA alignment

Wild strain RNA sequencing data publicly available on NCBI (PRJNA669810) was used for alignment to their respective wild strain genomes. RNA reads were aligned using *STAR* v2.7.11b^138^ in basic two-pass mode with a maximum intron size of 10 kilobases (*--alignIntronMax 10000*). RNA alignments were used to validate non-N2 reference gene model predictions in wild strains (Supplementary Figure 29). Gene model predictions in wild strains between genes *srh-277* and *srh-281* did not have RNA support, suggesting these gene models are predictions of the N2 reference pseudogene *srh-280* given conserved flanking gene order. Additionally, for overlapping N2 reference gene models *srz-13*, *srh-277*, and *srz-14*, *BRAKER3* was only able to make a single gene prediction at this locus in wild strains for *srh-277*, and RNA seq was not able to reliably validate the presence of *srz-13* and *srz-14*. For these reasons, non-N2 reference genes for *srh-280* and N2 reference genes *srz-13* and *srz-14* were removed from the QTL visualization (Figure 6). For expression analysis of N2 reference genes among wild strains, we used the publicly available expression dataset for 207 *C. elegans* wild strains^66^. Of the 207 strains, 48 overlapped with our pangenome strain set and were used for expression analyses (Supplementary Figures 8 and 31).

### Haplotype visualization

Wild strain whole-genome alignments to the N2 reference genome was performed using *NUCmer* v3.1^131^. Wild strain genome alignments were filtered for those that overlapped with the specified N2 genome region of interest for each wild strain. Alignments were then filtered for the wild strain contig that spanned the N2 genome region of interest the most to ensure the removal of multi-alignments from other contigs. To remove distal intra-contig alignments, we removed alignments that were >150 kb away from the most proximal upstream and downstream alignments. Alignments were then trimmed to be within 20 kb of the boundaries of the region of interest. Orthologous genes and strain-specific genes among all specified wild strains and the N2 reference strain were extracted for the region of interest. Genomic coordinates of all genes were left-adjusted to be in relation to the first syntenic gene among all specified strains. Colored polygons were added to connect orthologous genes, and wild strain-specific genes were colored in grey (Supplementary Figure 29). For simplistic display, all identified orthologous and strain-specific genes received psuedo-coordinates to represent gene order among all strains (Figure 6).

## Data Availability

The raw PacBio HiFi and Oxford Nanopore Technologies long-read sequencing data are publicly available from NCBI accession PRJNA1558913. Wild strain *de novo* genome assemblies are publicly available on NCBI under accession PRJNA1558933 and at CaeNDR. The SNVs used for analyses are publicly available at CaeNDR (Release: 20250625)^41^. Scripts and related raw and processed data are available at GitHub (https://github.com/AndersenLab/Ce_pangenome_MS/).

## Acknowledgements

We thank members of the Andersen Laboratory for their feedback on this manuscript. We thank CaeNDR, which is supported by the National Science Foundation Capacity grant (2224885), and we thank the citizen scientists who continue to contribute wild strains of *C. elegans* to CaeNDR.

## Author Contributions

L.M.O., N.D.M., and E.C.A. conceived and designed the study.

L.M.O., N.D.M., and A.O.S analyzed the data.

L.M.O. and E.C.A. wrote the manuscript.

N.D.M., R.E.T., A.O.S., R.P., O.R., and S.Z. edited the manuscript.

R.P., O.R., S.Z., and L.K. provided long-read DNA sequencing for two *C. elegans* wild strains.

R.E.T. and A.K. collected DNA from *C. elegans* wild strains for long-read DNA sequencing.

