## Supplementary Figures for "The *Caenorhabditis elegans* pangenome elucidates hyper-divergent region gene content to reveal extensive gene copy-number variation and a novel anthelmintic resistance locus"

### SUPPLEMENTARY DATA

**Supplementary tables:**

Supplementary Table 1: *C. elegans* wild strain genome assembly statistics.
Supplementary Table 2: Cumulative length of wild strain structural variants.
Supplementary Table 3: Most abundant IPR terms for genes that overlap with SVs.
Supplementary Table 4: SVs that are unique to 19 Hawaiian strains that cluster in PCA.
Supplementary Table 5: Orthogroups among 142 strains of *C. elegans*.
Supplementary Table 6: Wild strain HDR coordinates after lift-over.
Supplementary Table 7: Wild strain genes in HDRs.
Supplementary Table 8: Statistics of wild strain HDRs and the genes contained in them.
Supplementary Table 9: IPR annotations of 142 *C. elegans* strains.
Supplementary Table 10: Enriched IPR terms in wild strain HDRs.
Supplementary Table 11: Gene family classifications from IPR descriptions.
Supplementary Table 12: Genotype matrix of strains used in genome-wide association mapping.
Supplementary Table 13: Macrocyclic lactone resistance trait file used for genome-wide association mapping.
Supplementary Table 14: Genome-wide association mapping of abamectin q90 TOF.
Supplementary Table 15: Annotation of shared SNVs across resistant strains.
Supplementary Table 16: Orthology of *C. elegans* genes associated with abamectin resistance with *H. contortus*.

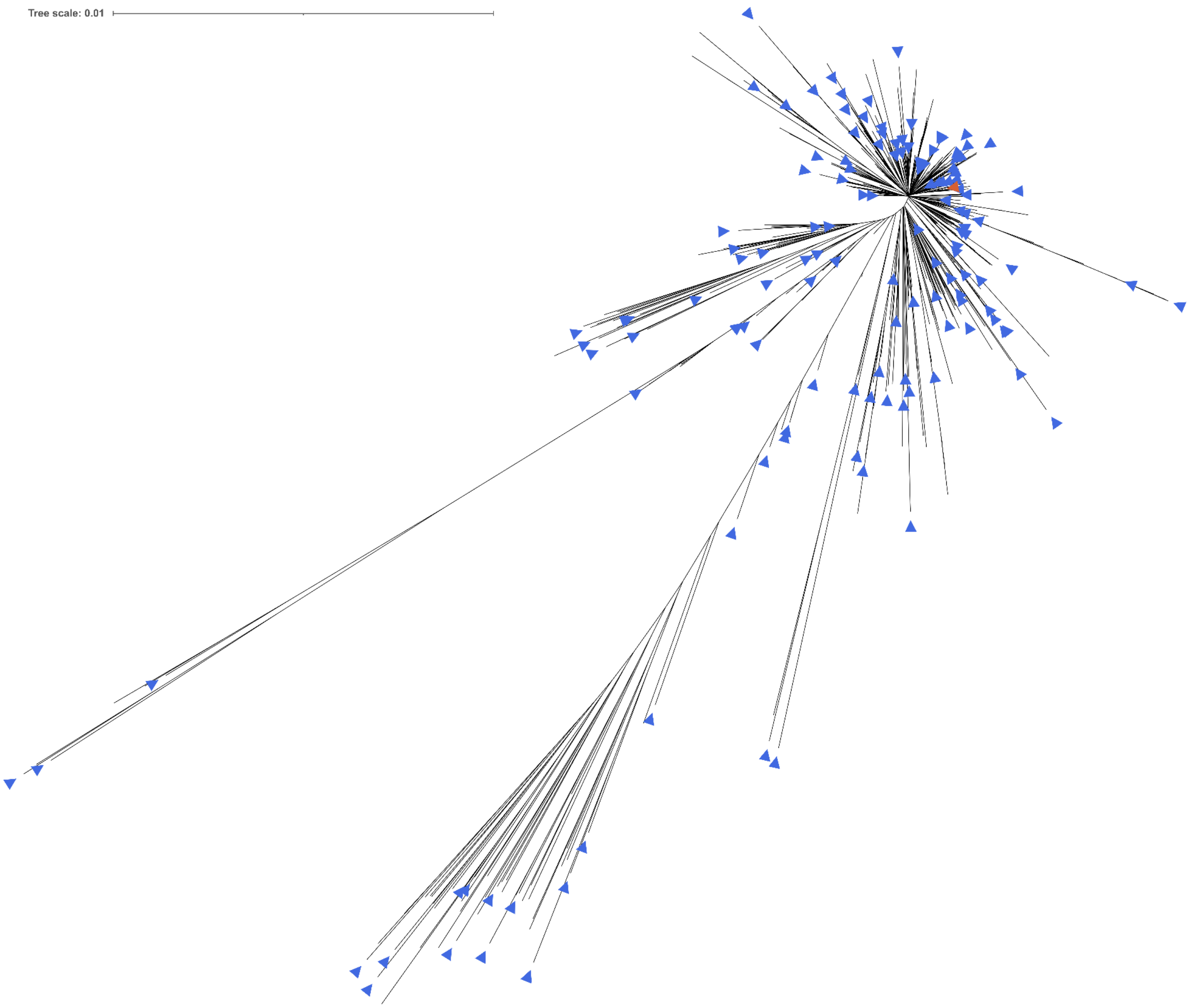


**Supplementary Figure 1. Maximum-likelihood tree of 684 genetically distinct *C. elegans* strains.** Tree constructed using linkage-disequilibrium-pruned (removed $r^2 > 0.9$) biallelic SNVs and IQ-TREE[1] with the GTR model. The 140 selected strains for PacBio HiFi sequencing are indicated with blue triangles, and the N2 reference strain with an orange triangle. The scale bar represents a 1% substitution rate of non-missing SNVs.

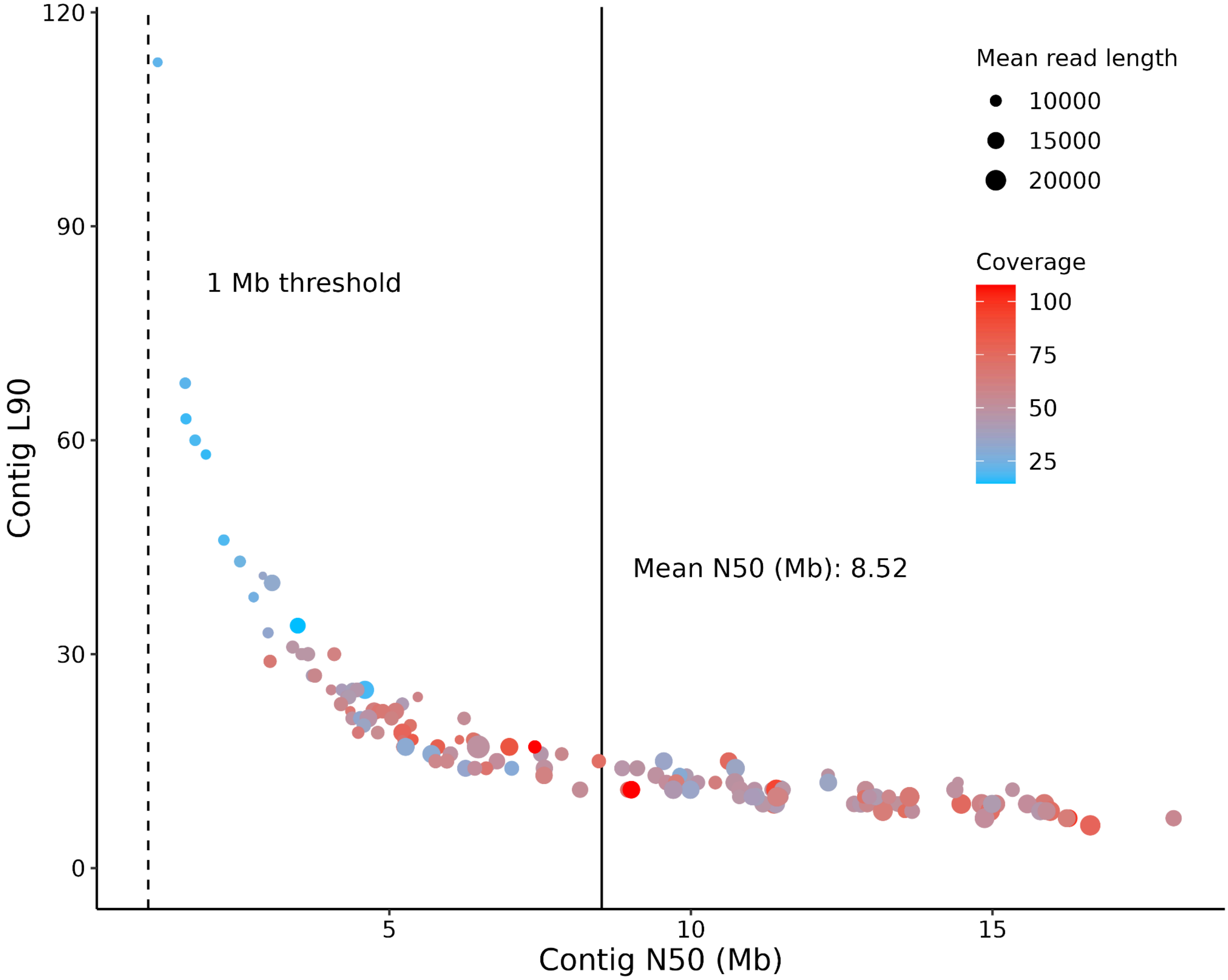


**Supplementary Figure 2. Genome assembly statistics of wild *C. elegans* strains.** Each point represents a wild strain genome and is sized by the mean HiFi read length and colored by fold-coverage. The vertical dashed line represents the 1 Mb threshold used for downstream genome assemblies, and the vertical solid line represents the mean N50 of genome assemblies with an N50 > 1 Mb (140 assemblies). ECA741 has a coverage of 406.82 (Supplementary Table 1) and was artificially given a coverage of 100x for plotting purposes.

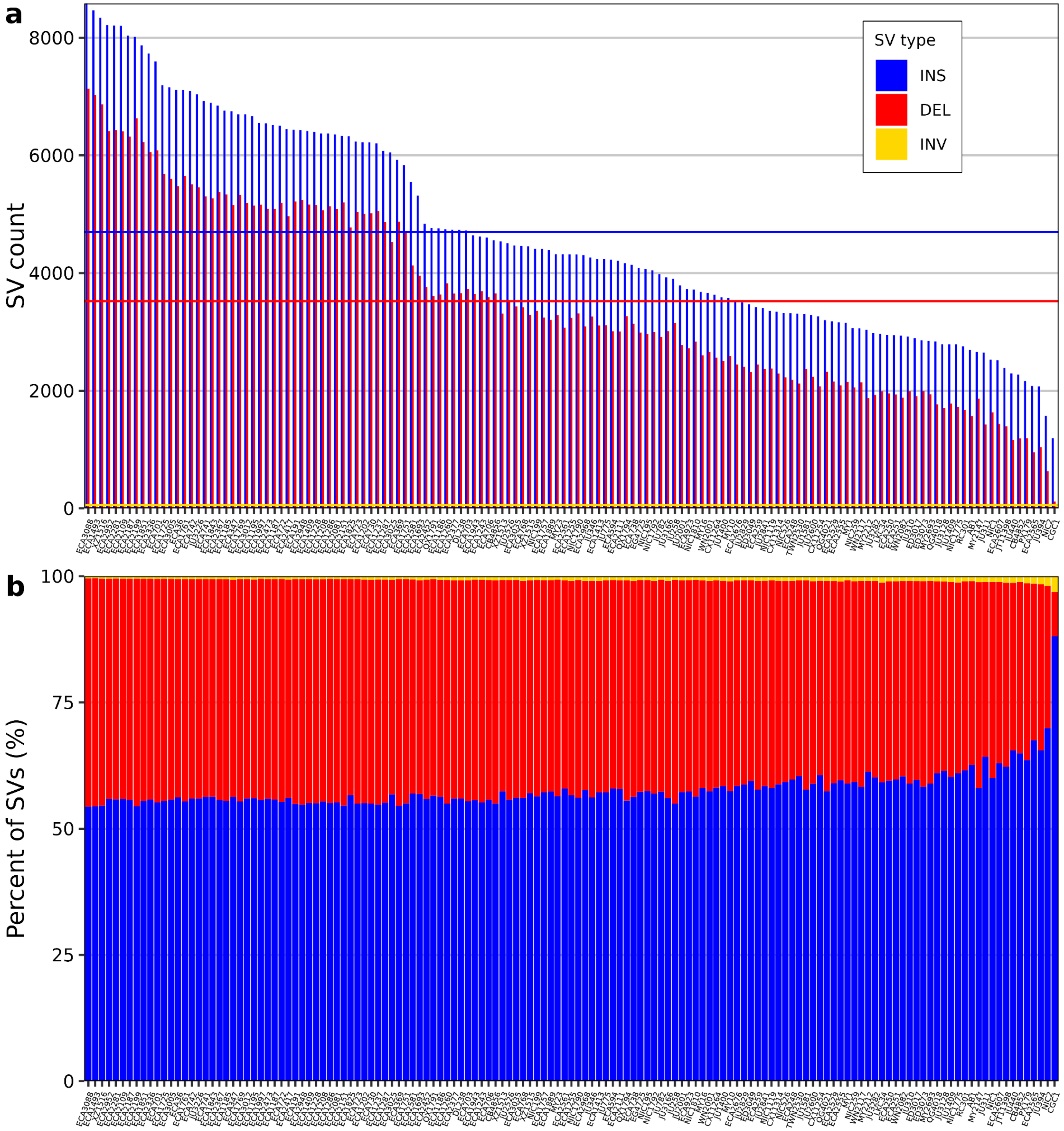


**Supplementary Figure 3. SV counts among 141 *C. elegans* strains. a)** The count of SVs, colored by SV type, for all *C. elegans* strains in the pangenome (x-axis). The horizontal lines indicate the average SV count (insertions: 4,700, deletions: 3,523, inversions: 60), colored by SV type. **b)** The percent of each SV type for all SV calls for every strain.

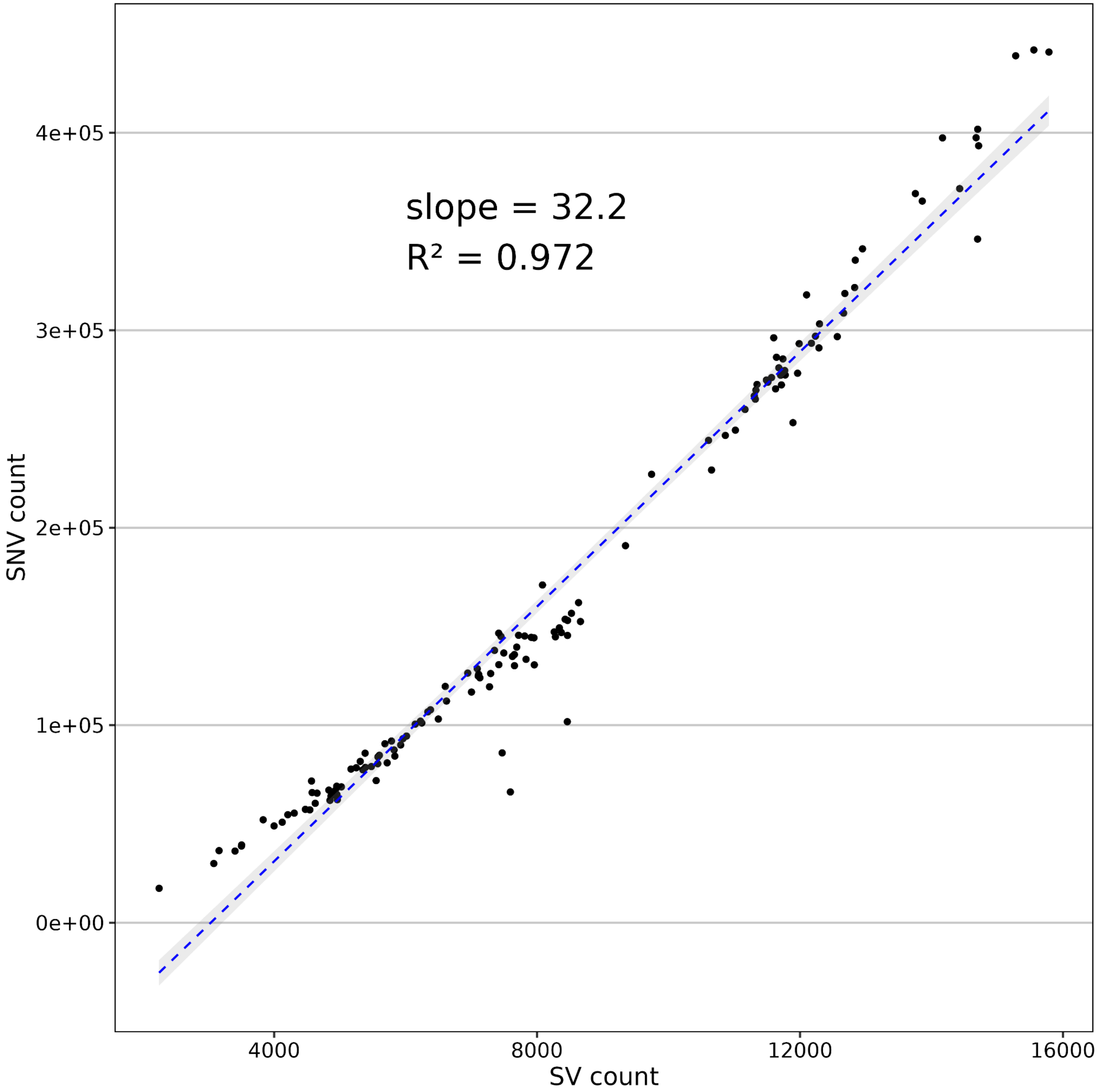


**Supplementary Figure 4. SVs correlate with SNVs among 140 *C. elegans* wild strains.** Structural variant counts (x-axis) and SNV (y-axis) counts correlation estimated using linear regression. The dashed blue line represents an ordinary least squares regression, and gray shading represents 95% confidence of the model fit.

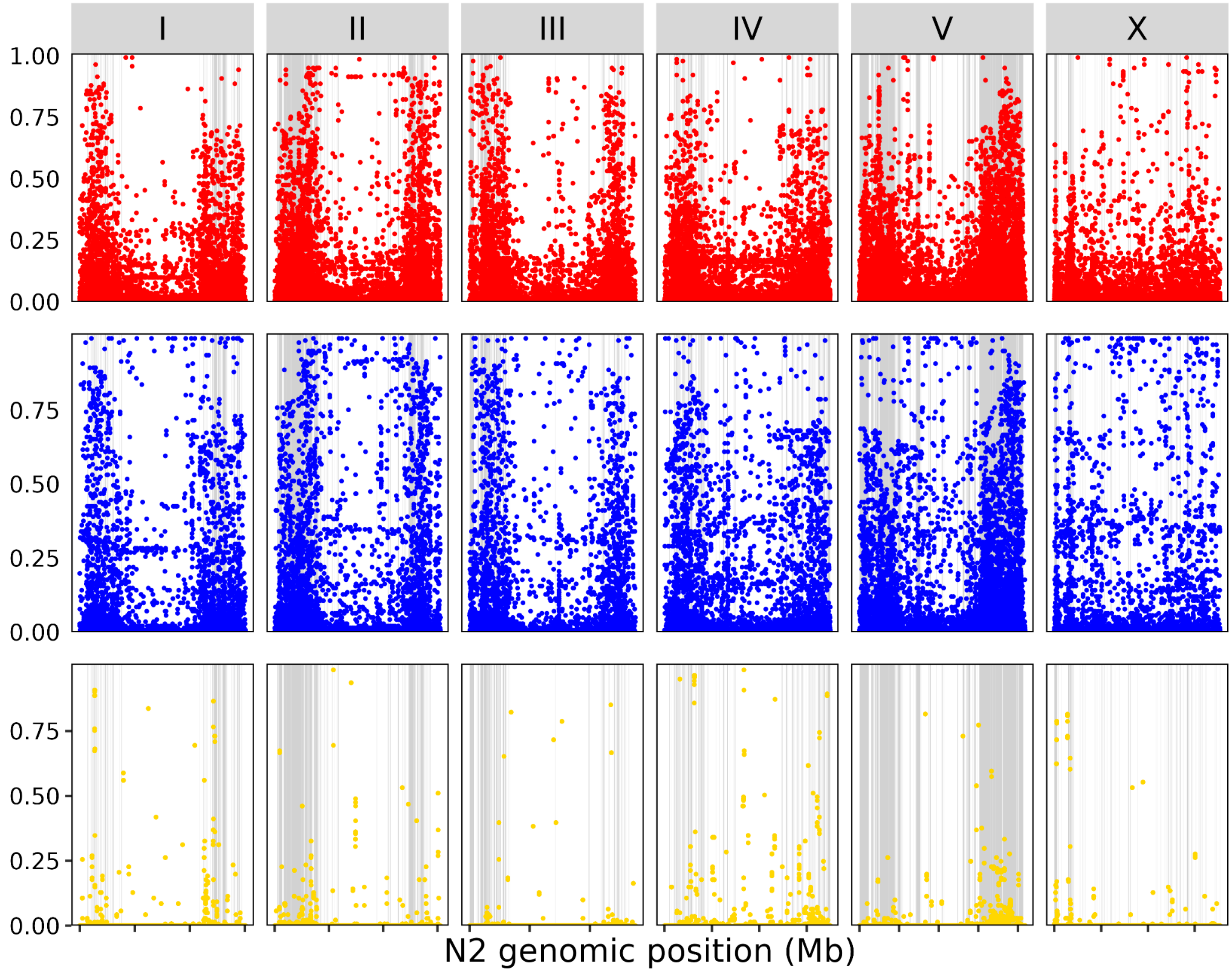


**Supplementary Figure 5. SVs are enriched in HDRs.** The frequency that SVs among 140 wild strains of *C. elegans* (y-axis) are identified per 10 kb (x-axis). The N2 genome coordinates are displayed in megabases, with each x-axis tick representing 5 Mb. HDRs identified among the 140 pangenome strain set are collapsed and represented with light grey rectangles. Deletions are represented in red, insertions in blue, and inversions in gold. Enrichment was calculated as: ((number of unique SVs in HDRs / number of base pairs in HDRs) / (number of unique SVs outside HDRs / number of base pairs outside HDRs)).

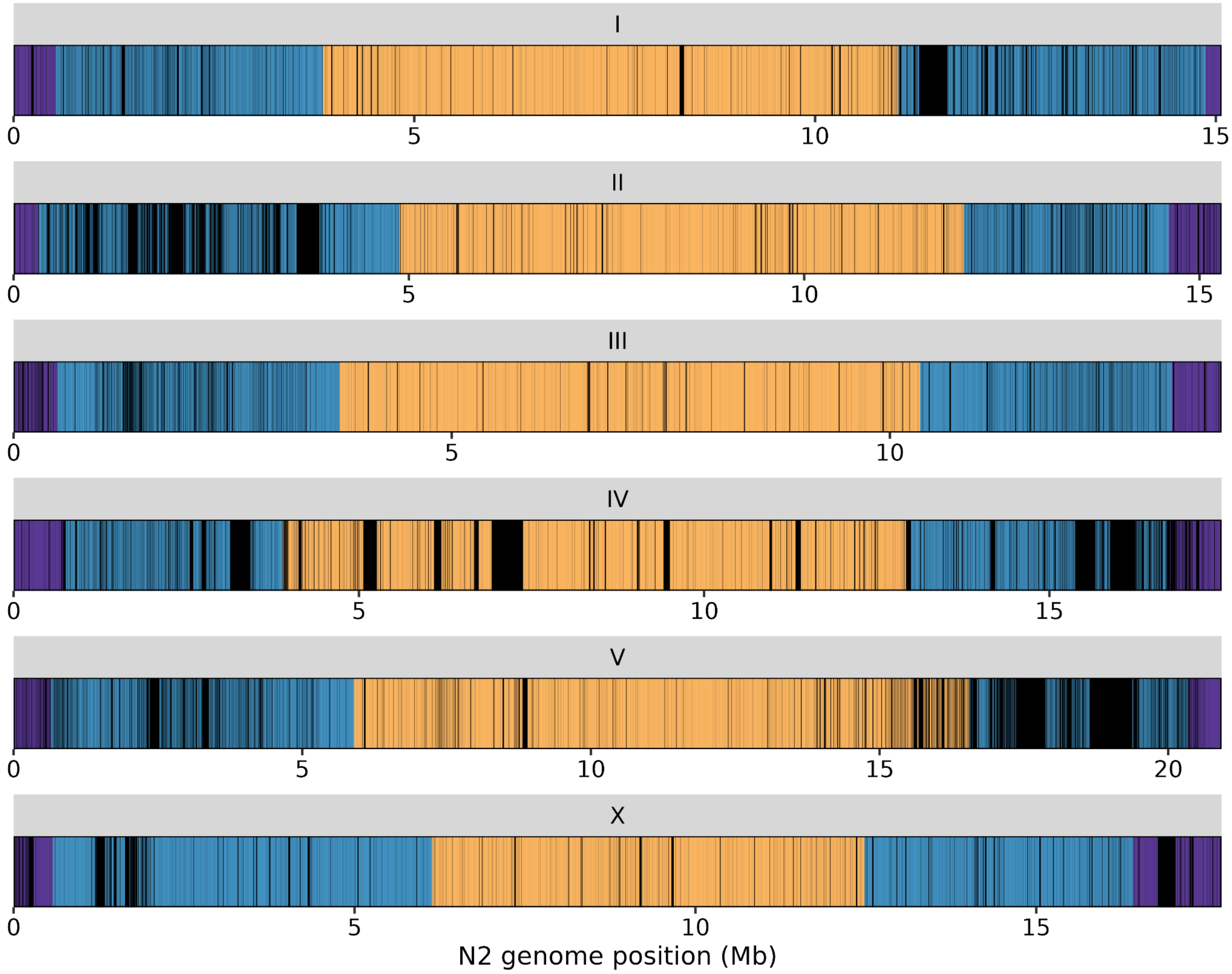


**Supplementary Figure 6. SVs are densely populated on chromosome arms.** Collapsed deletion and inversion structural variants identified among 141 strains of *C. elegans* are represented by black rectangles. N2 chromosomes (facets) are shaded by domains (purple = tips, blue = arms, gold = centers)[2].

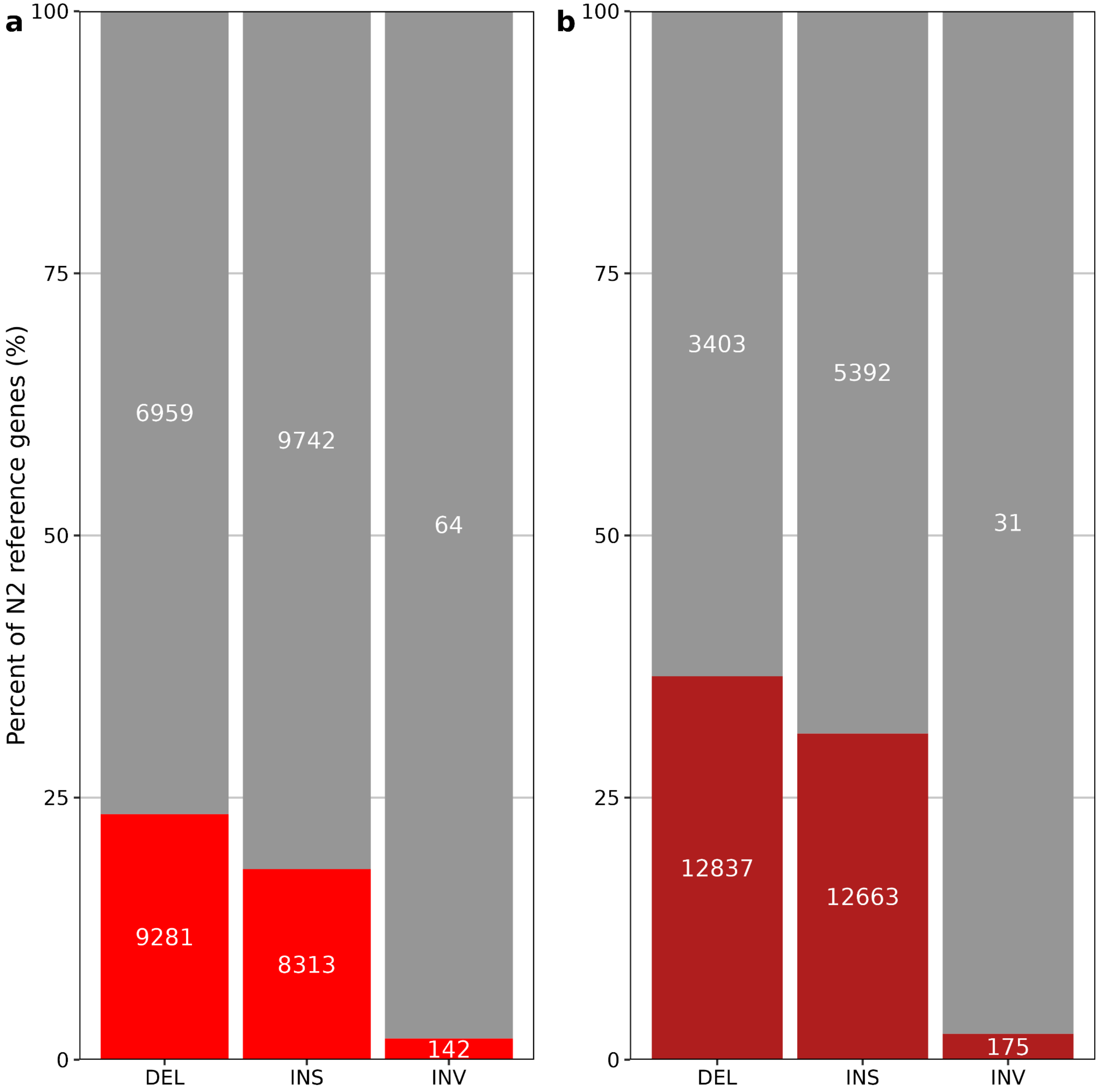


**Supplementary Figure 7. SVs overlap with N2 reference genes. a)** The red bars represent the percent of N2 reference genes that have at least one SV overlapping, faceted by SV type. The white numbers inside the bars display the number of SVs that overlap with N2 genes (red) or do not overlap with N2 genes (grey). **b)** The same as panel **a)**, but SVs overlapping with an N2 gene or within 2 kb of the coding region.

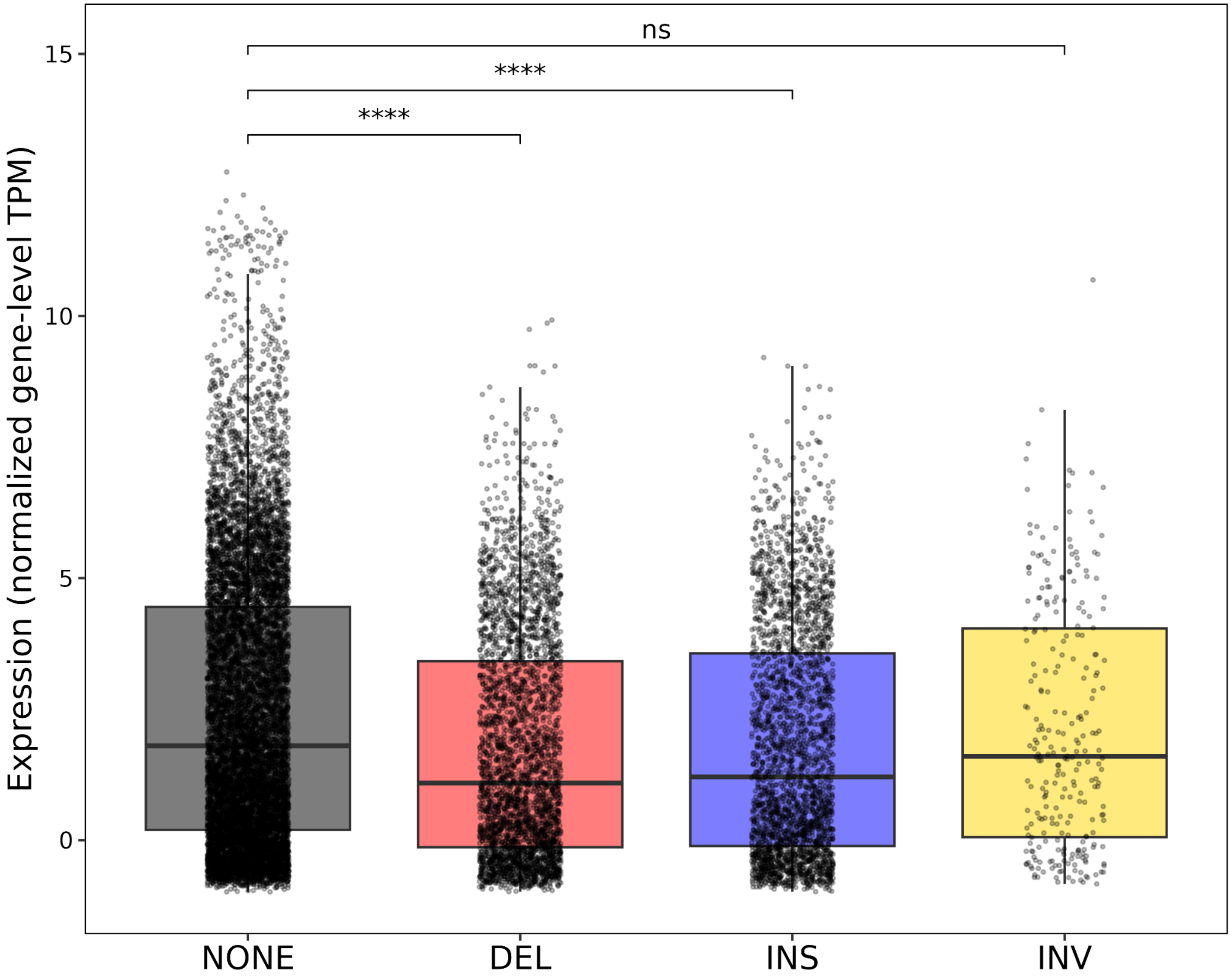


**Supplementary Figure 8. N2 reference genes that are structurally divergent across wild strains have lower expression.** N2 reference genes are classified based on SV overlap, where N2 reference genes with no overlapping SVs are displayed in gray (NONE), genes with overlapping deletions in red (DEL), genes with overlapping insertions in blue (INS), and genes with overlapping inversions in gold (INV). Statistical significance was determined using the Wilcoxon rank-sum test. *$p < 0.05$, **$p < 0.01$, ***$p < 0.001$, ****$p < 0.0001$, ns = not significant.

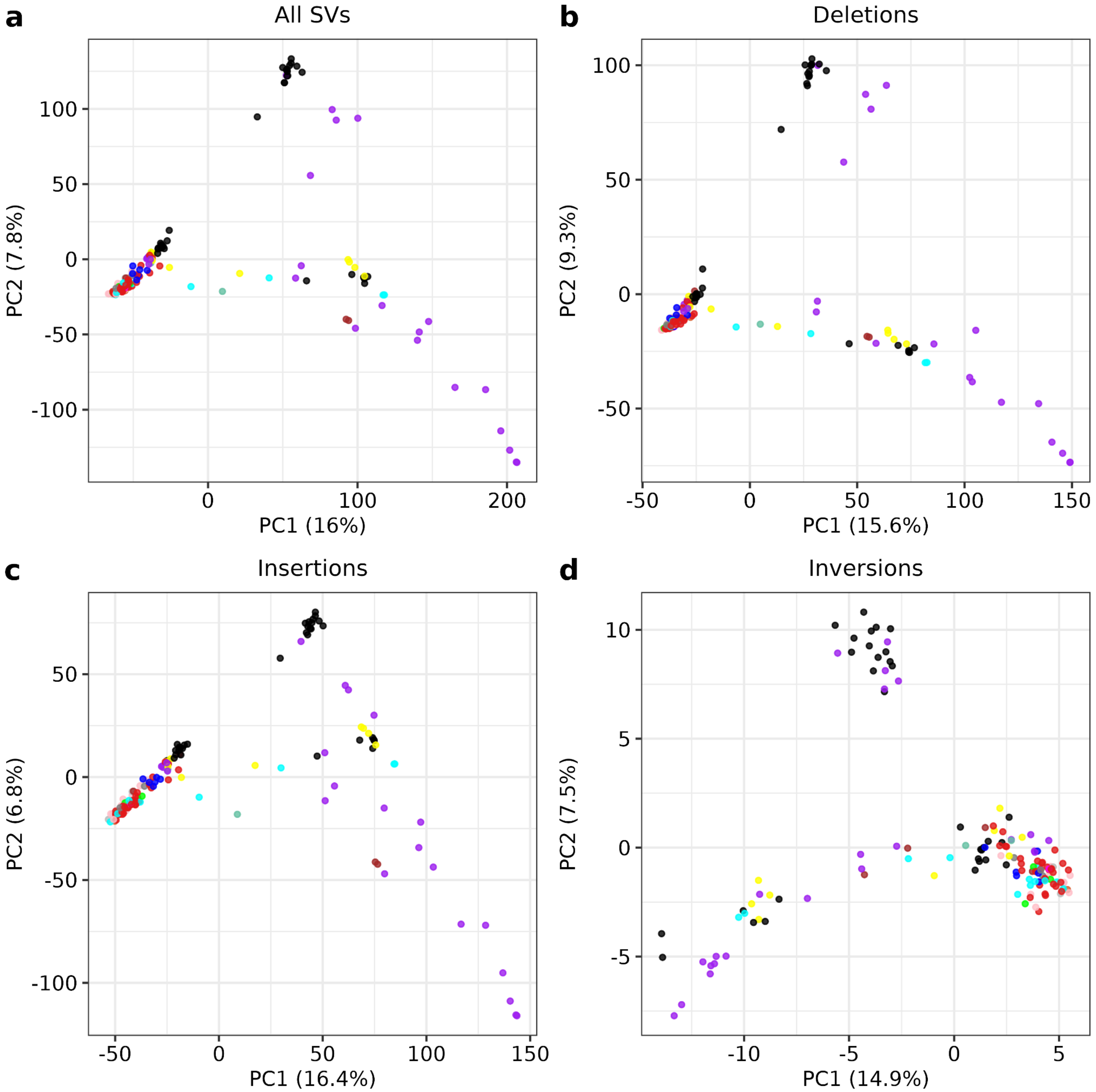


**Supplementary Figure 9. PCA of each SV type.** PCA of all SVs (**a**), deletions (**b**), insertions (**c**), and inversions (**d**). Each point is colored by the sampling location. Black = Big Island; bright green = Africa; blue = Atlantic; orange = CGC1; red = Europe; purple = Kaua'i; yellow = Maui; sage green = Moloka'i; pink = North America; brown = O'ahu; cyan = Oceania; magenta = South America; grey = unknown.

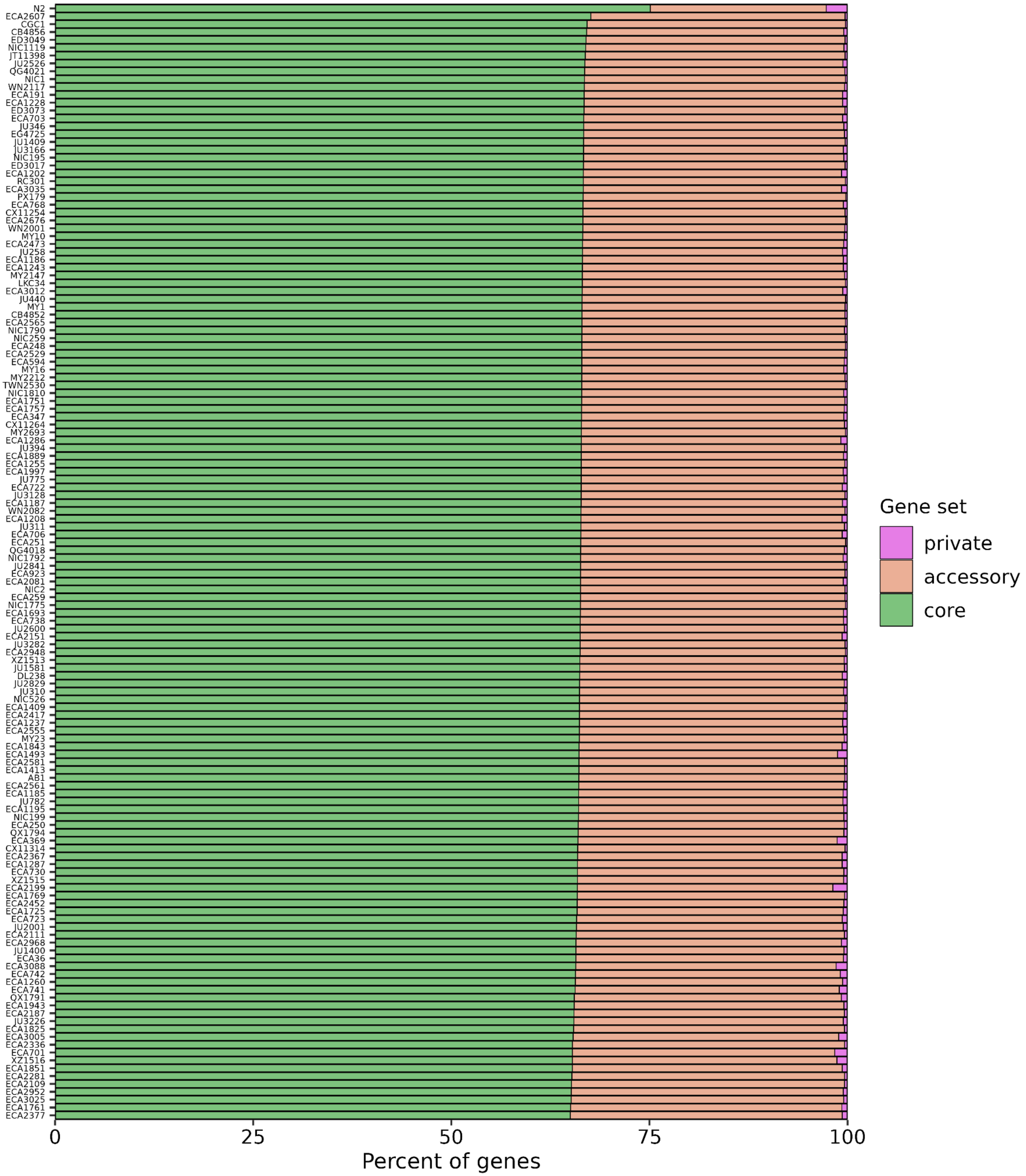


**Supplementary Figure 10. Proportion of genes contributing to each pangenome gene set for every wild strain.** Each row is a strain in the pangenome strain set, with the percent of genes contributing to each gene set displayed on the x-axis.

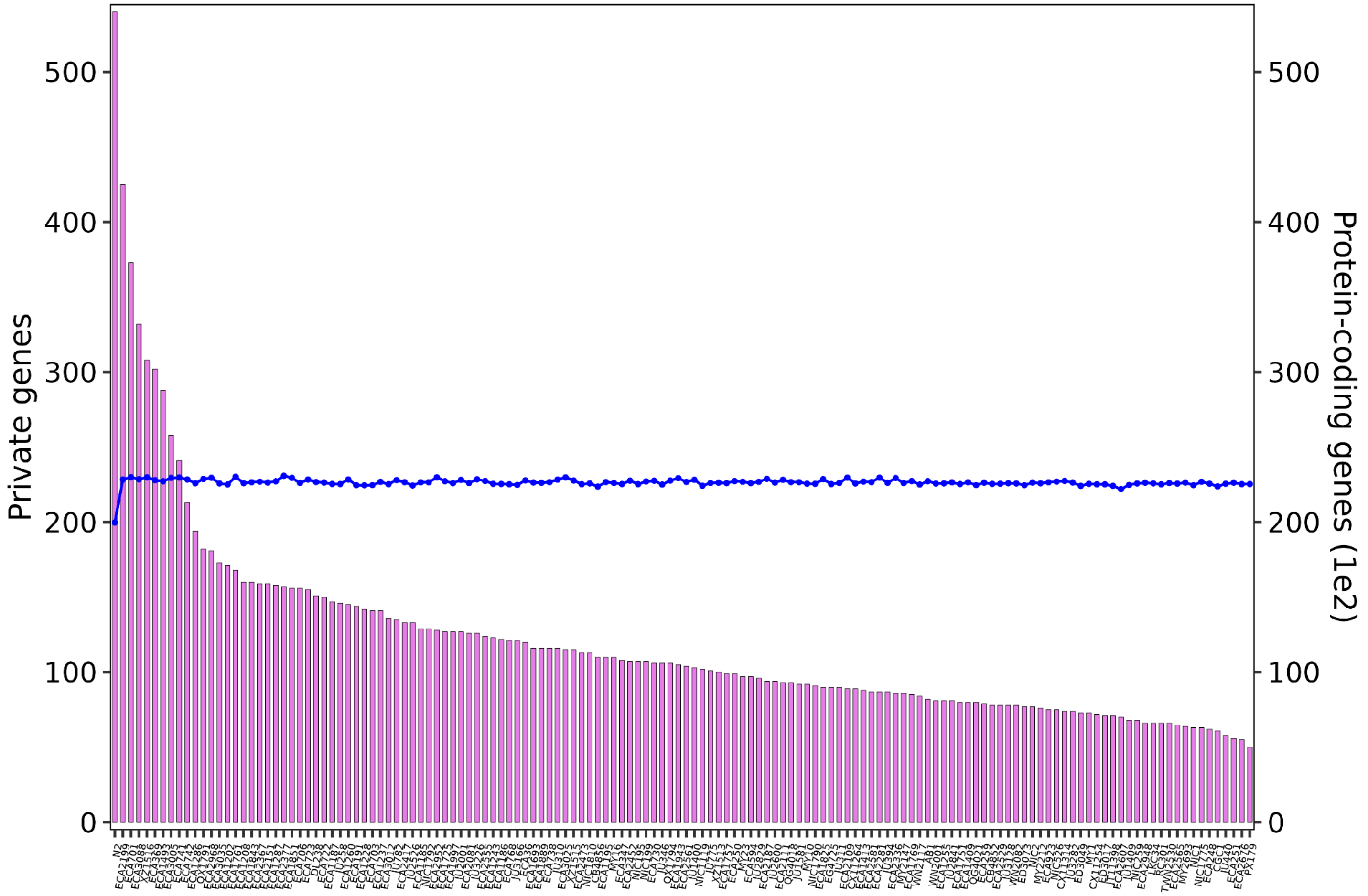


**Supplementary Figure 11. The number of predicted protein-coding genes for all strains and the number that contribute to the private pangenome.** The number of private genes for each wild strain is displayed on the left y-axis, represented by pink bars, and the total number of predicted protein-coding genes is displayed on the right y-axis by blue points.

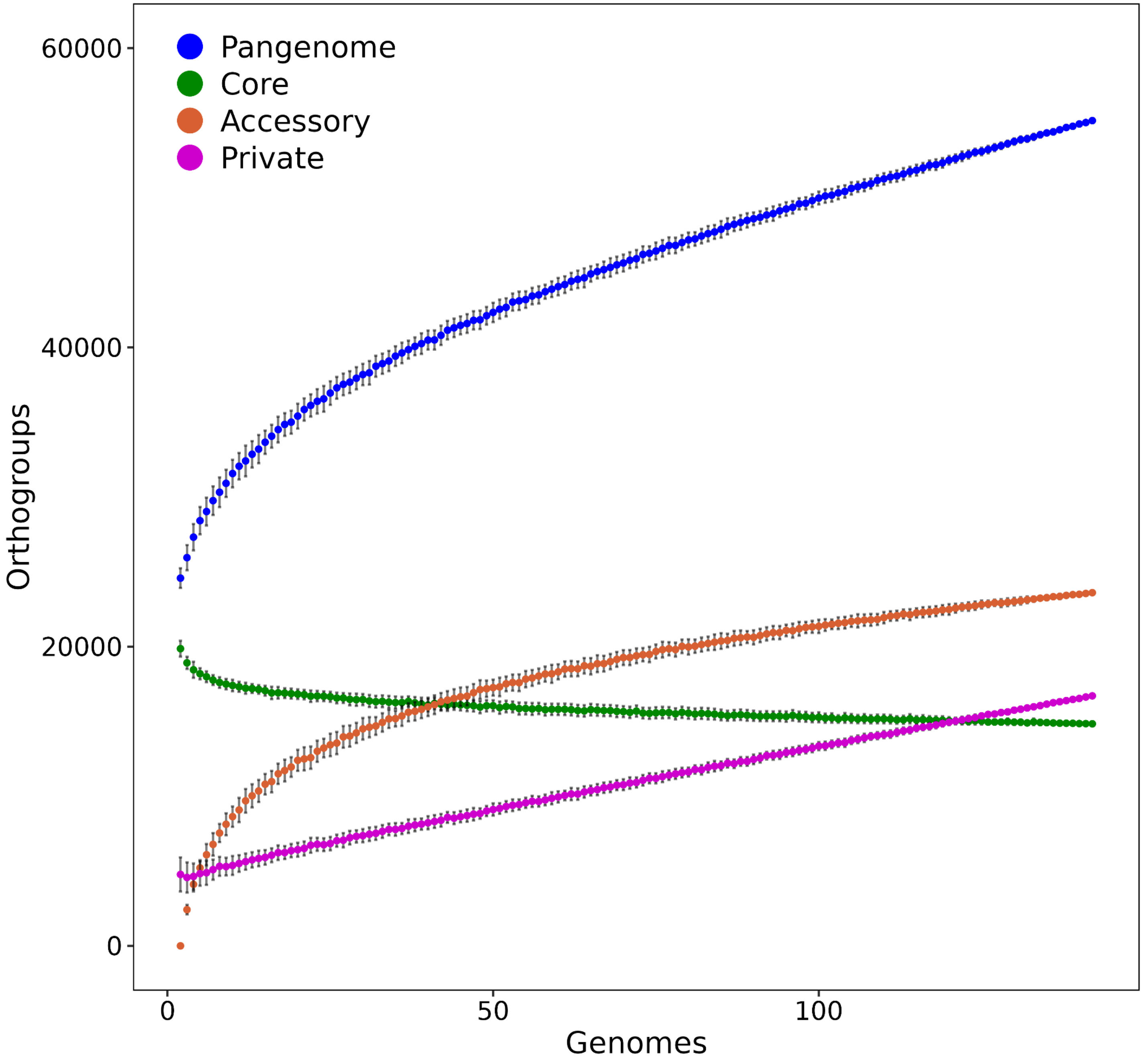


**Supplementary Figure 12. Rarefaction curves of pangenome gene sets.** Rarefaction curves of all orthogroups (pangenome, blue), core orthogroups (green), accessory orthogroups (orange), and private orthogroups (pink). Colored points display the mean value for each subsample of genomes (between 2 - 142) after 100 iterations, and error bars represent one standard deviation from the mean.

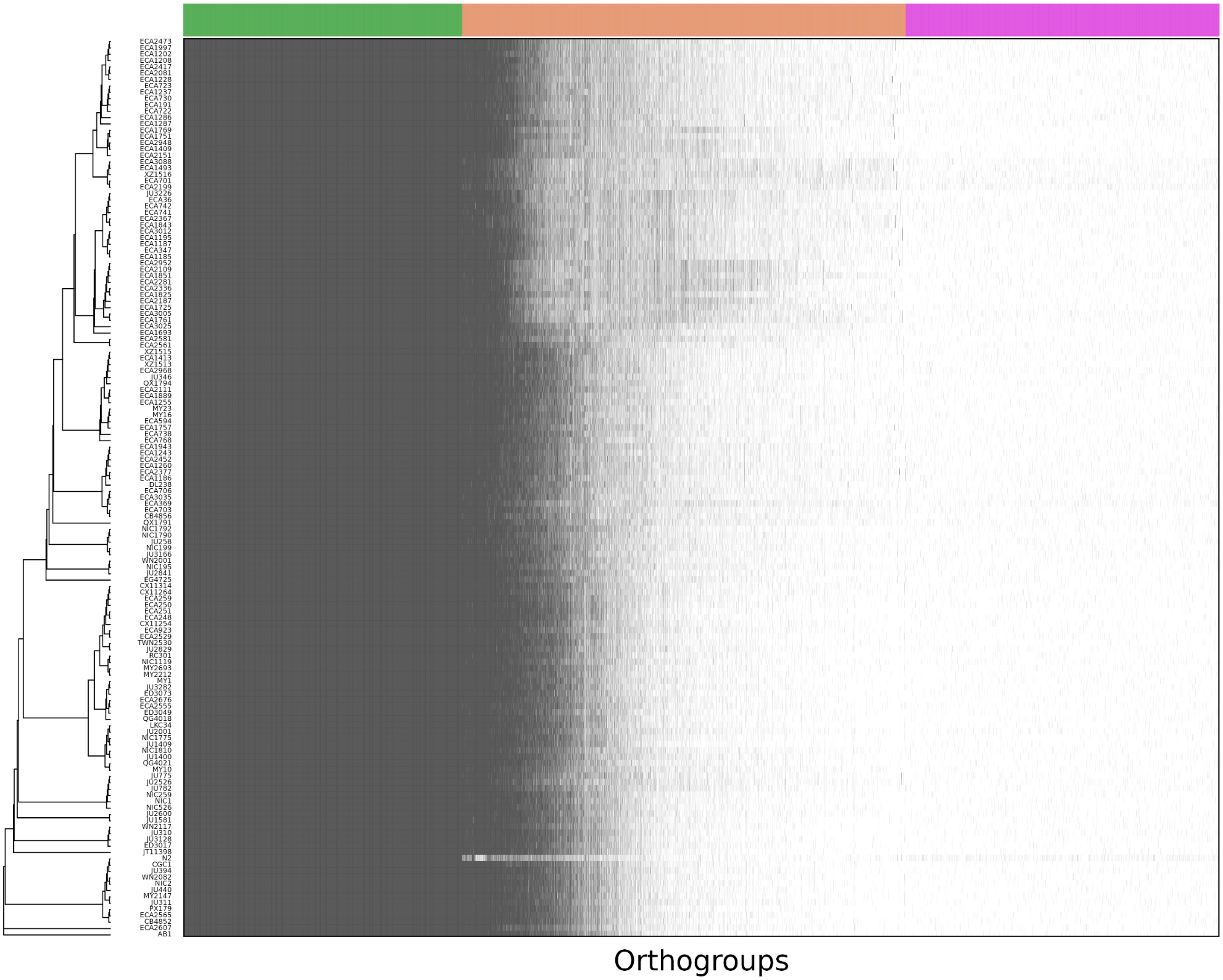


**Supplementary Figure 13. Relatedness structure inferred from BUSCOs poorly explains accessory pangenome variation.** A presence-absence heatmap of orthogroups (x-axis) and all 142 *C. elegans* strains in the pangenome strain set (y-axis). Dark grey rectangles indicate that a wild strain has an ortholog contributing to an orthogroup. Pangenome gene set classifications of orthogroups are represented by the colored rectangle (core orthogroups: green; accessory orthogroups: orange; private orthogroups: pink).

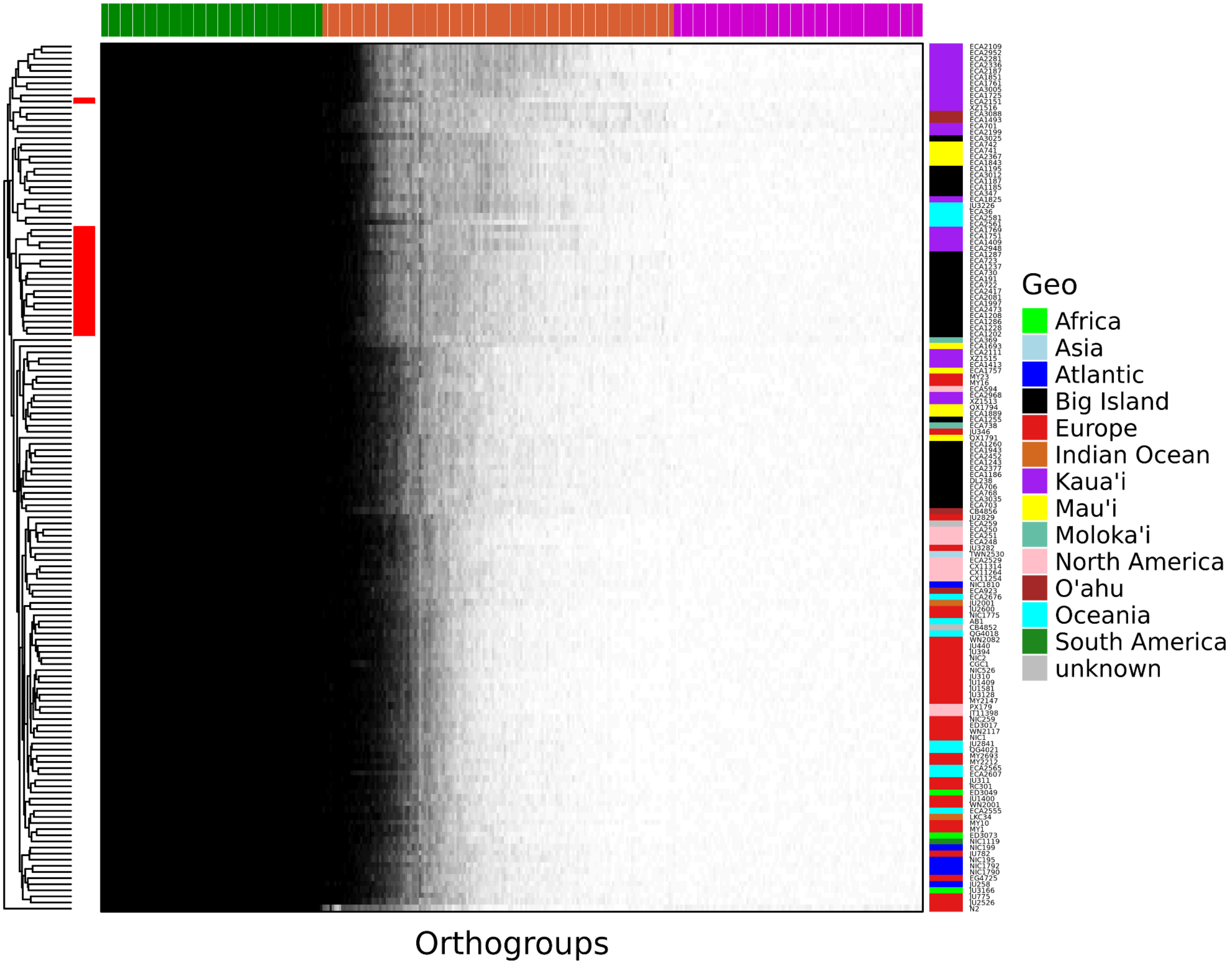


**Supplementary Figure 14. Accessory pangenome clustering reveals relatedness-group-specific clusters.** A presence-absence heatmap of orthogroups (x-axis) and all 142 *C. elegans* strains in the pangenome strain set (right y-axis). Black rectangles indicate that a wild strain has an ortholog contributing to an orthogroup. Pangenome gene set classifications of orthogroups are represented by the colored rectangle (core orthogroups: green; accessory orthogroups: orange; private orthogroups: pink). Strains were clustered based on accessory orthogroup sharing (dendrogram, left y-axis), and 19 Hawaiian strains that cluster in SV PCA also cluster based on accessory pangenome gene content (red bar) (Figure 2b).

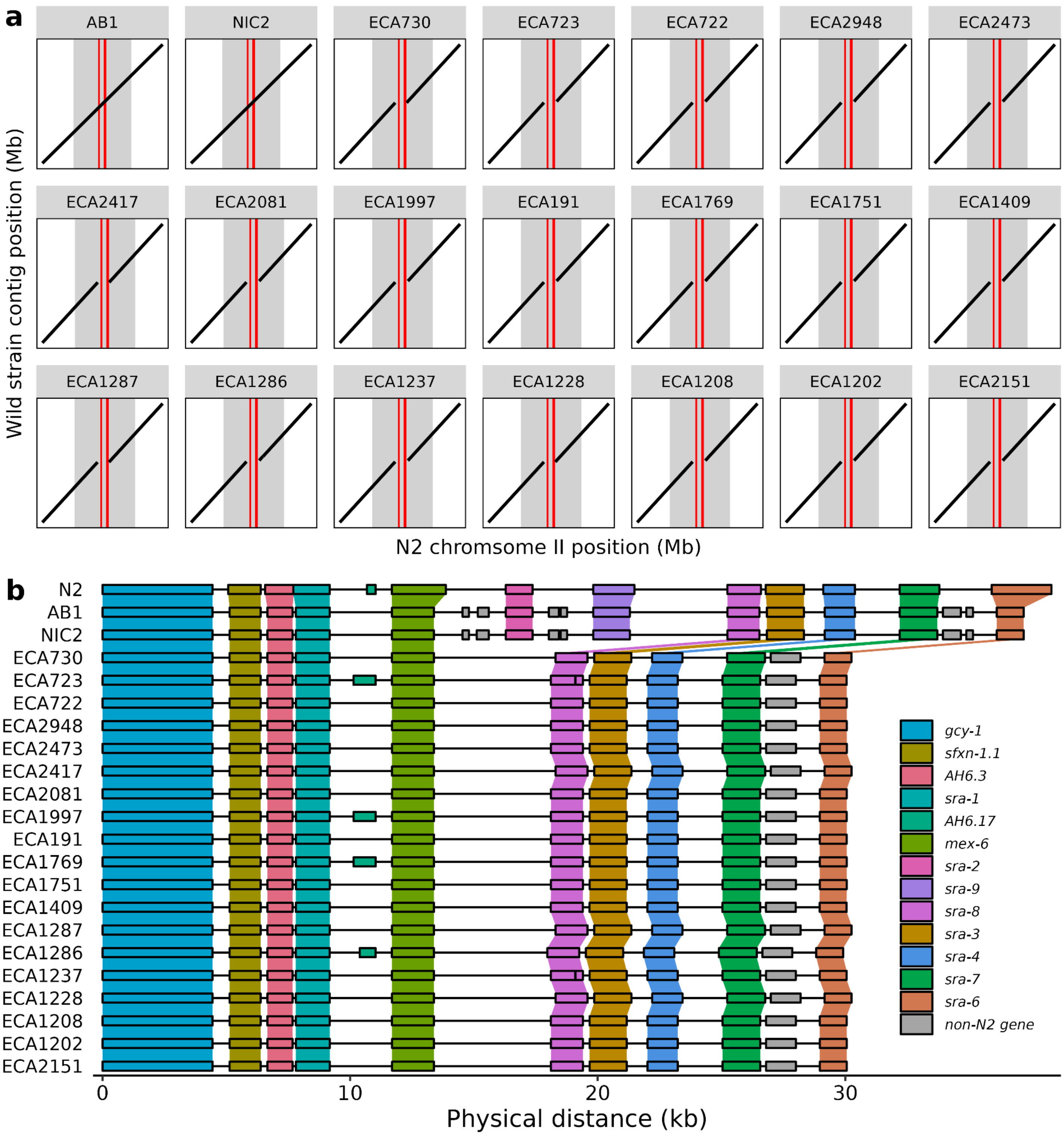


**Supplementary Figure 15. Sub-population-specific loss of *sra-2* and *sra-9* in 18 Hawaiian strains. a)** Genome-genome alignments of 19 Hawaiian strains, AB1, and NIC2 to the N2 reference genome. Wild strains AB1 and NIC2 were chosen for visualization purposes because they are closely related to the N2 reference strain and exhibit the same haplotype at the displayed locus. The vertical grey rectangle represents the genomic region that is visualized in (**b)** and vertical red rectangles represent the loci of *sra-2* and *sra-9*. **b)** Synteny visualization of the N2 reference genome region surrounding *sra-2* and *sra-9*. Orthogroups are colored by N2 reference genes and connected with colored banding. Wild strain genes that do not have an N2 ortholog are grey.

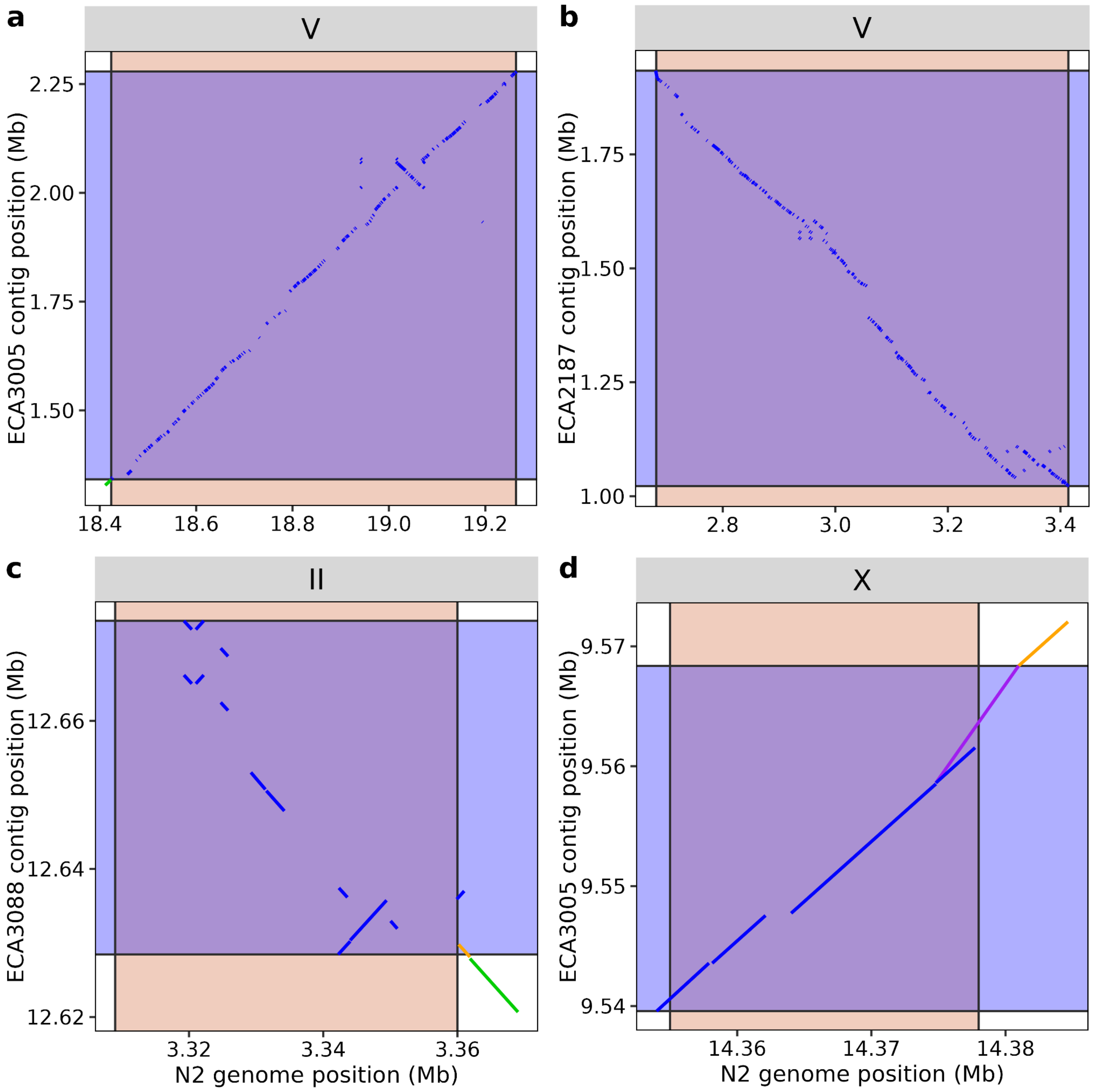


**Supplementary Figure 16. Lift-over of HDRs from N2 reference genome coordinate space to wild strain genome coordinate space.** An HDR is displayed on the x-axis marked by vertical black lines and filled in orange for **a)** ECA3005, **b)** ECA2187, **c)** ECA3088, **d)** and ECA3005 in reference N2 coordinate space. Coordinate lift-over of HDRs to wild strain genome space is marked by horizontal black lines filled in blue (y-axis). Lift-over of HDR boundaries to wild strain coordinate space is performed based on contig alignments spanning N2 HDR boundaries or using the max or min coordinate of the most proximal contig alignment to the N2 HDR boundary (see Methods). Green line = lagging; blue line = overlapping; orange line = leading; purple line = extension.

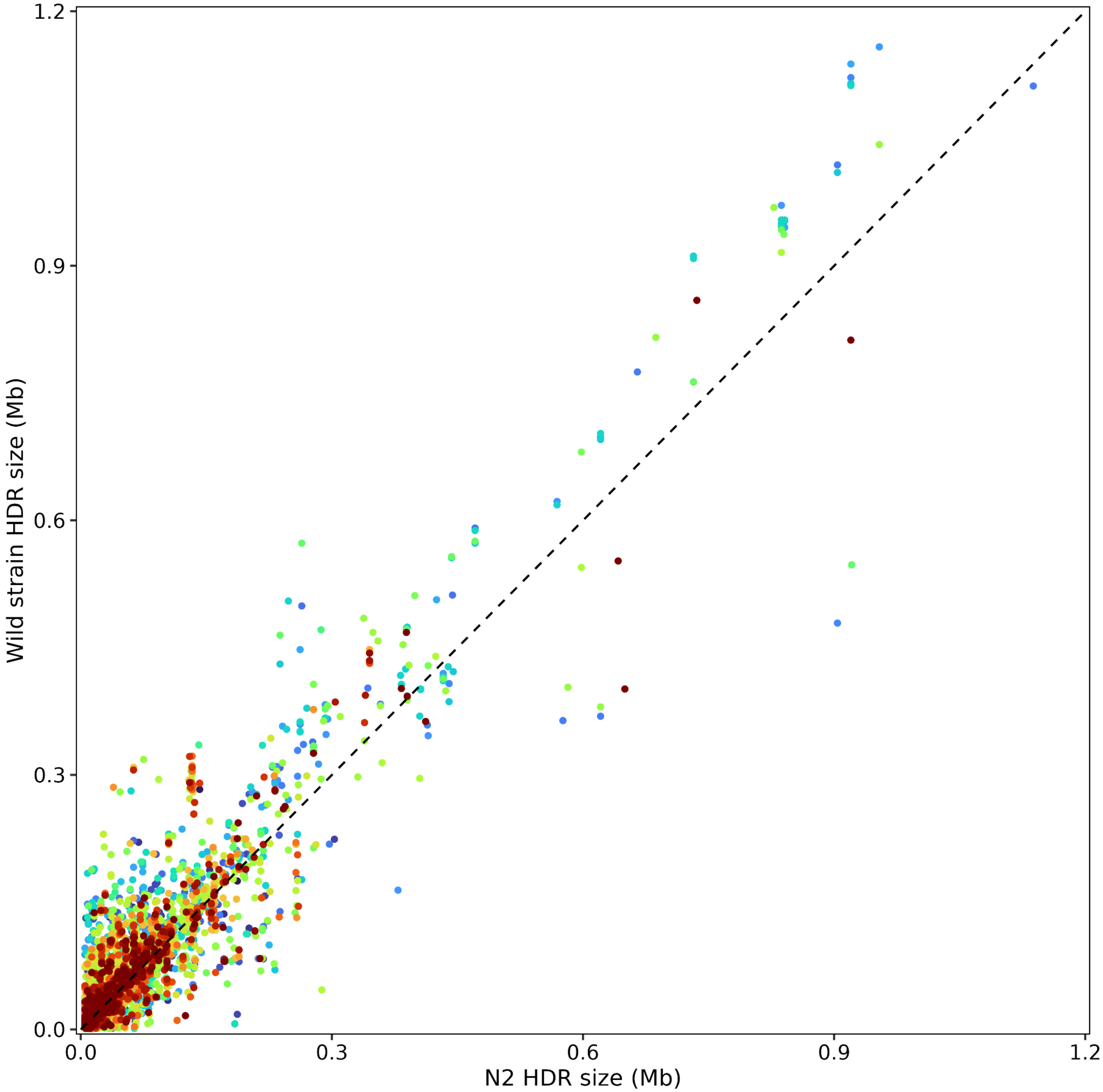


**Supplementary Figure 17. Difference in HDR sizes after wild strain coordinate lift-over.** The sizes of HDRs identified in relation to the N2 reference genome for 140 wild strains are displayed on the x-axis. Each point represents a single HDR and is colored by wild strain identity. Sizes of HDRs in wild strain genomic coordinates after coordinate lift-over are displayed on the y-axis. The dashed line represents the y = x slope.

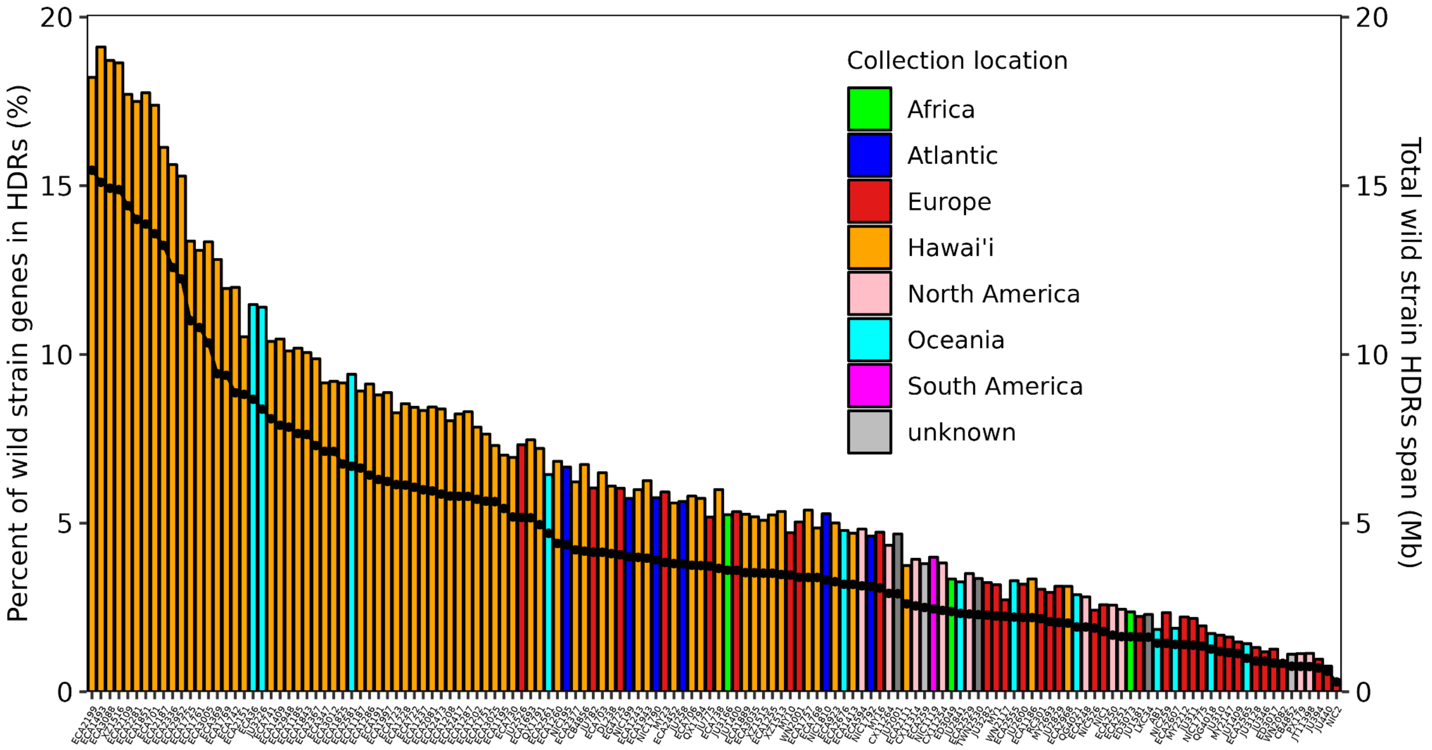


**Supplementary Figure 18. The span and percent of genes in wild strain HDRs.** The percent of genes in HDRs for every wild strain (left y-axis, bar plot) and the cumulative span of HDRs (right y-axis, line plot). Strains are colored by their sampling location.

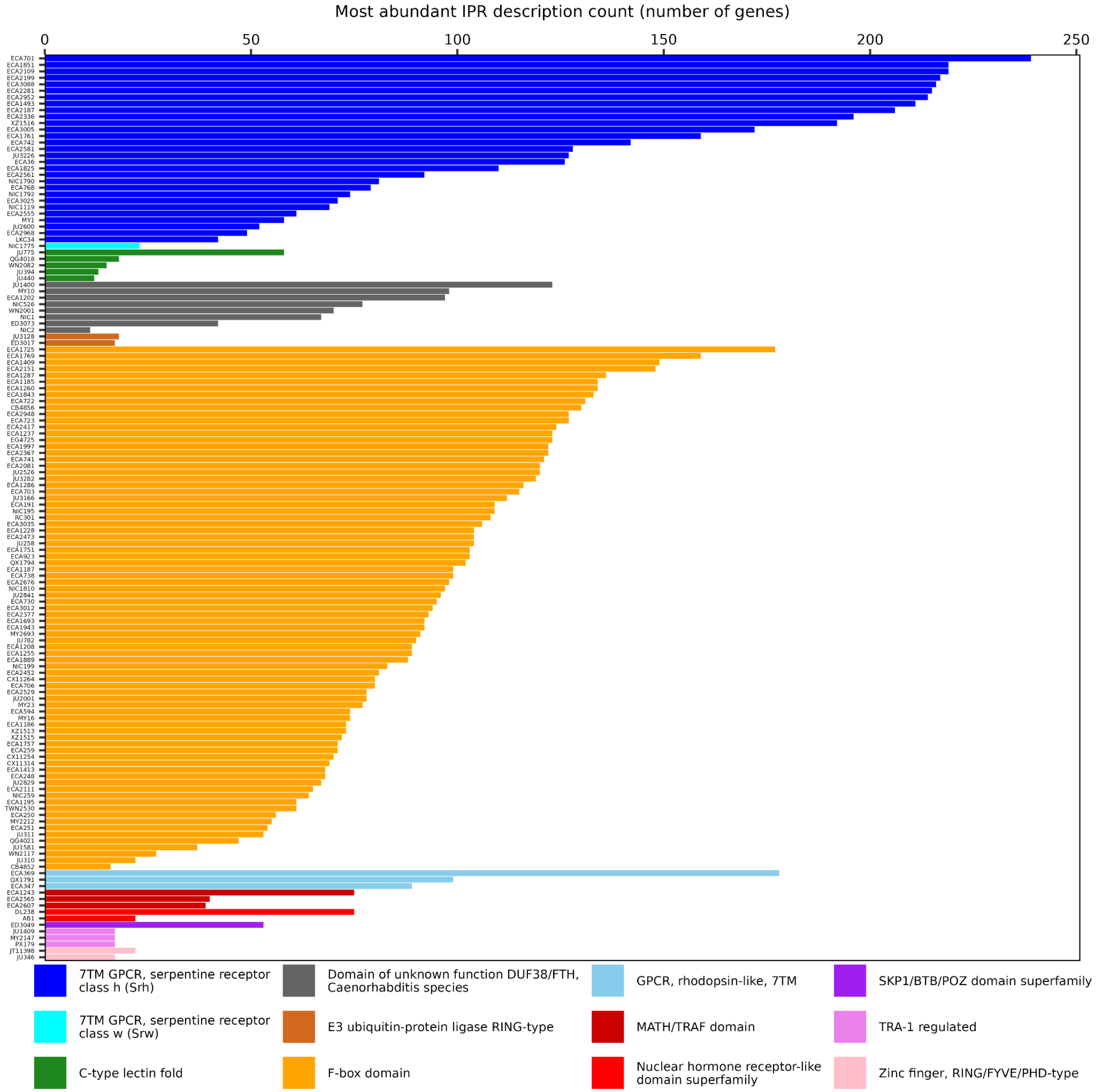


**Supplementary Figure 19. The most abundant IPR terms in wild strain HDRs.** For every wild strain (x-axis), the most abundant IPR description (*i.e.*, the most abundant protein) for genes in HDRs is shown. Colors correspond to IPR descriptions.

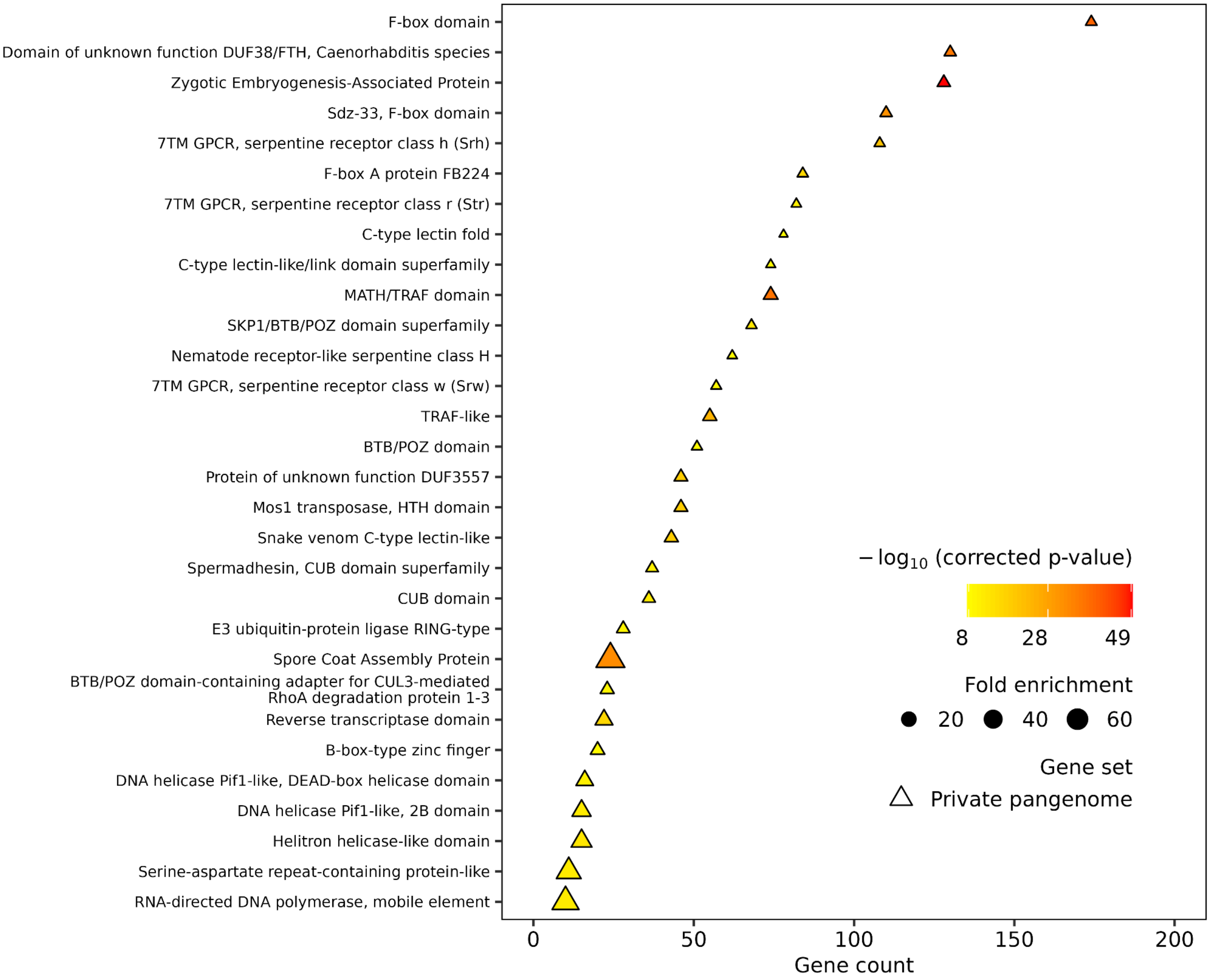


**Supplementary Figure 20. Wild strain HDRs are enriched for private environmental sensing and response genes.** IPR functional enrichment analysis using a hypergeometric test of wild strain private genes in HDRs using all genes in the pangenome as the background set. Corrected p-values are FDR-adjusted using Benjamini-Hochberg. Displayed are the top 30 most significantly enriched IPR annotations in the private HDR pangenome. Data points are shaped by the pangenome gene set, filled by statistical significance, and sized by fold enrichment in the corresponding pangenome gene set.

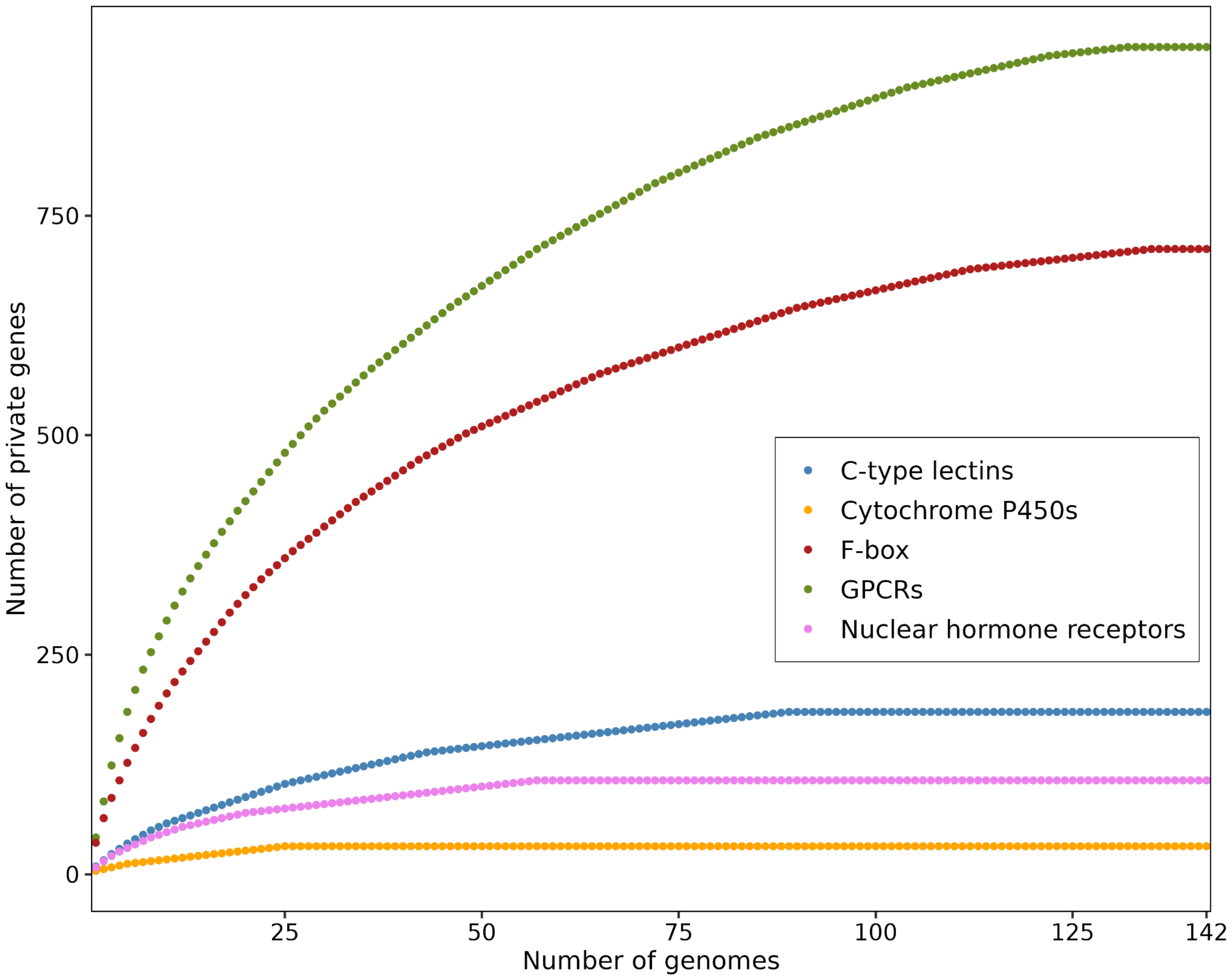


**Supplementary Figure 21. Private gene class rarefaction.** Strains were ordered on the x-axis by decreasing count of private genes (y-axis)for each gene class, indicated by color. All rarefaction curves level off (slope = 0), indicating that not all wild strains have private genes of the displayed gene classes.

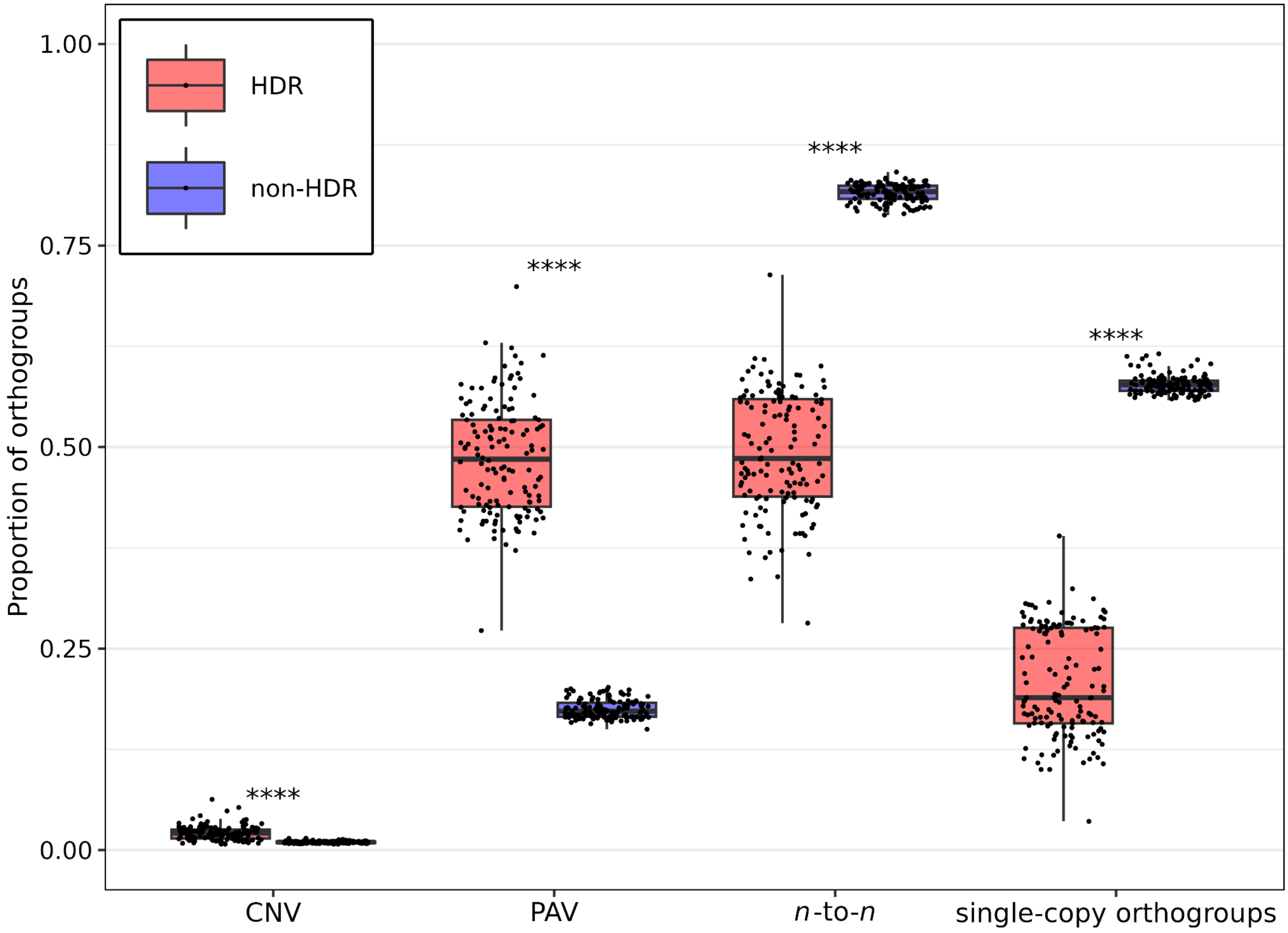


**Supplementary Figure 22. Enrichment of gene copy-number variation in HDRs.** Each point represents the proportion of orthogroups for a single strain. Each point is normalized by the total number of orthogroups inside or outside of HDRs for that strain, where the proportion of CNVs, PAVs, and *n*-to-*n*'s add up to one for each strain. The proportion of single-copy orthogroups (conserved among all 142 strains) is calculated independently and represents the proportion of orthogroups that are single-copy orthogroups out of all orthogroups for that strain (inside or outside of HDRs). Statistical significance was determined using the Wilcoxon signed-rank test with Benjamini-Hochberg correction. ****$p < 0.0001$.

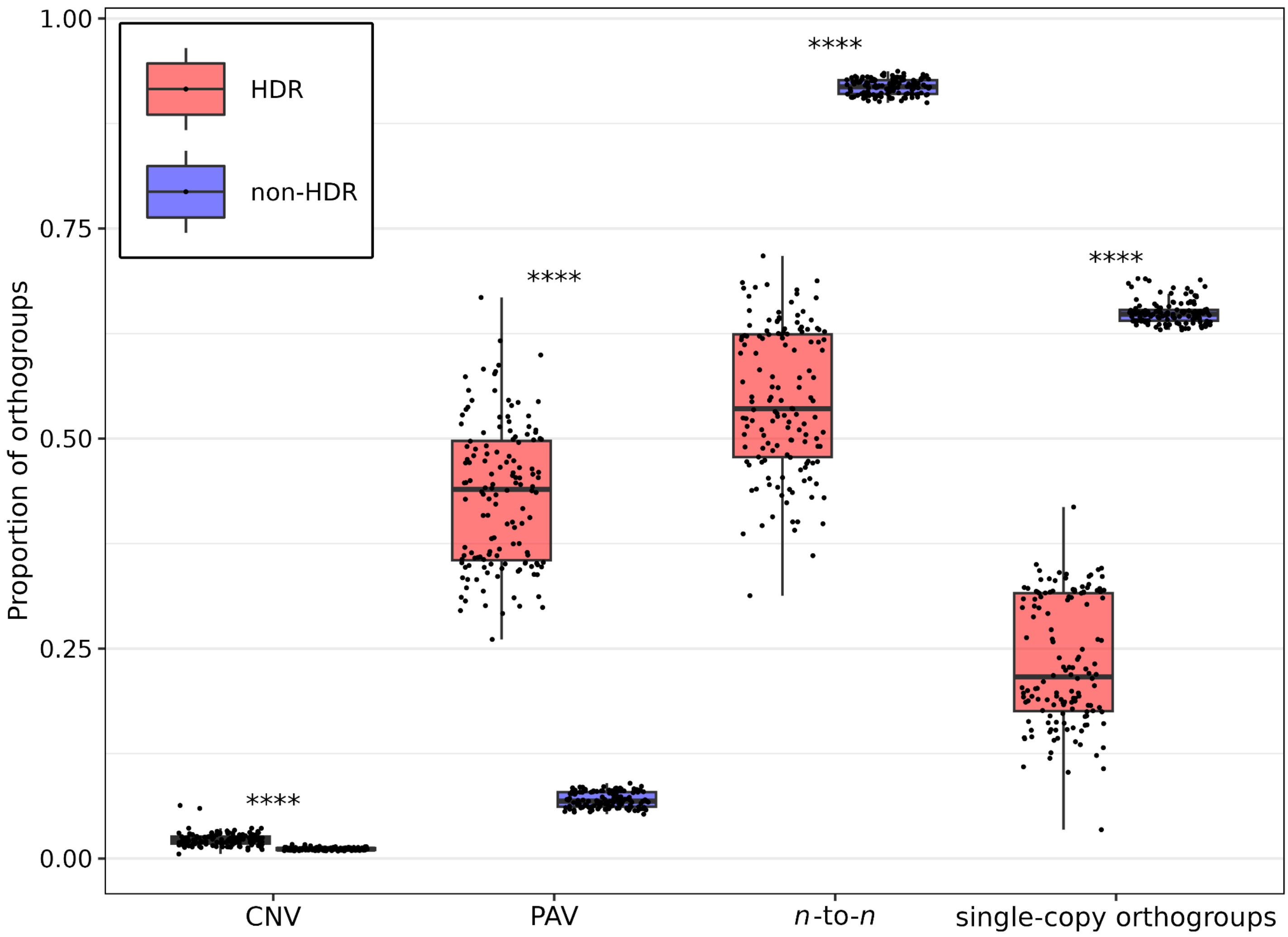


**Supplementary Figure 23. Enrichment of N2 gene copy-number variation in HDRs.** Each point represents the proportion of N2 orthogroups with HDRs classified for each of the 140 wild strains. Each point is normalized by the total number of orthogroups inside or outside of HDRs, where the proportion of CNVs, PAVs, and *n*-to-*n*'s will add up to one. The proportion of single-copy orthogroups (conserved among all 142 strains) is calculated independently and represents the proportion of orthogroups that are single-copy orthogroups out of all orthogroups for N2 (inside and outside of HDRs for each wild strain). Box and whisker plots display the median value for each metric on the x-axis. The upper and lower whiskers correspond to the first and third quartiles, respectively. Statistical significance was determined using the Wilcoxon signed-rank test with Benjamini-Hochberg correction. ****$p < 0.0001$.

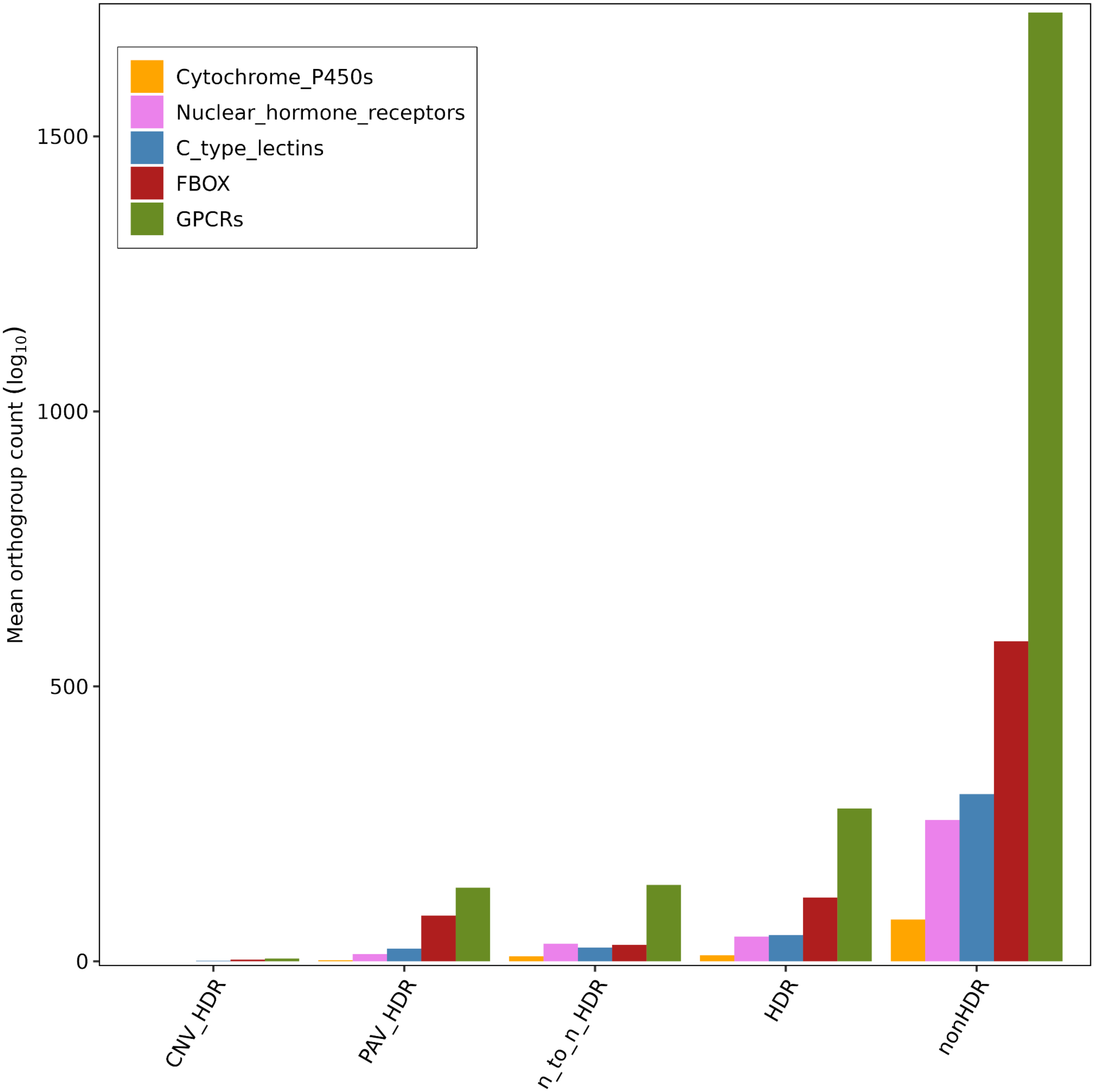


**Supplementary Figure 24. Average number of orthogroups among 140 wild strains for gene families enriched in HDRs.** Orthogroups are classified as CNVs, PAVs, or *n*-to-*n* based on their orthologous relationship with the N2 strain (see Methods) and then further classified as being inside (HDR) or outside of HDRs (nonHDR).

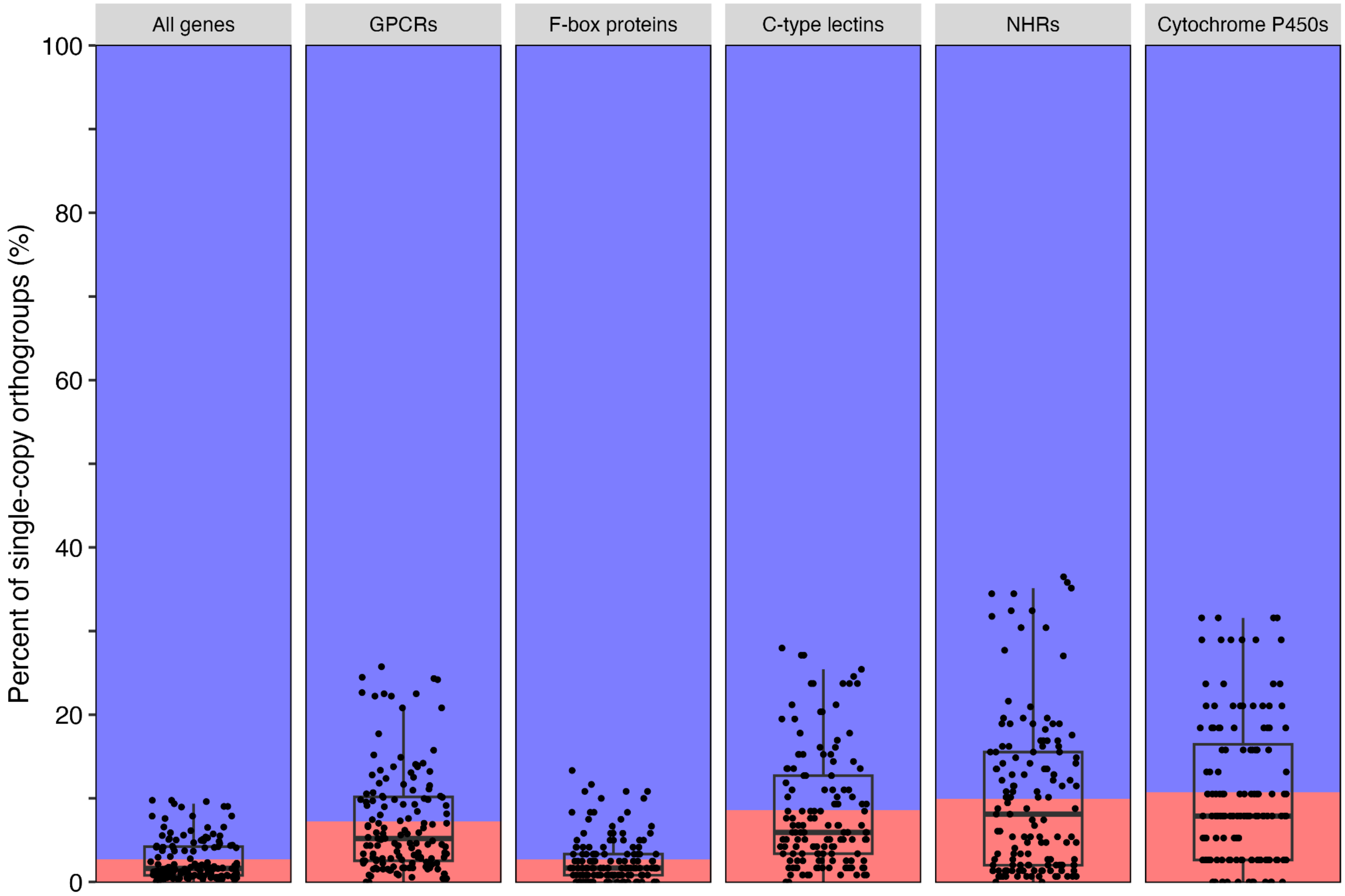


**Supplementary Figure 25. Percent of single-copy orthogroups found in HDRs.** The mean percent of single-copy orthogroups found in HDRs among all wild strains is colored in red, and the mean percent found outside of HDRs is colored in blue. Each point represents the percent for a wild strain with box and whisker plots displaying the median value for each facet, and the upper and lower whiskers correspond to the first and third quartiles, respectively.

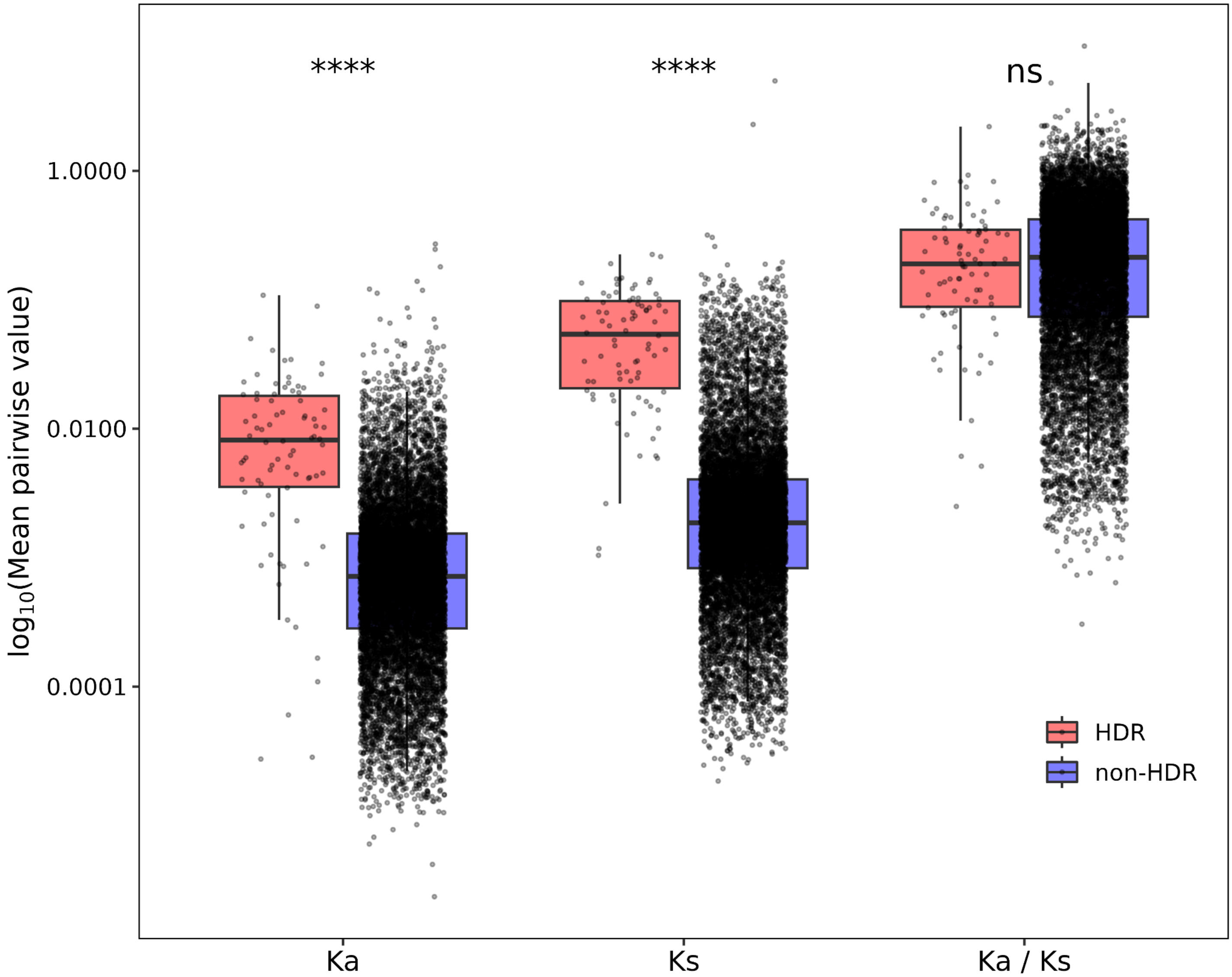


**Supplementary Figure 26. Single-copy ortholog sequence divergence is elevated in HDRs.** Synonymous ($K_s$), non-synonymous ($K_a$), and $K_a$ / $K_s$ rate among single-copy orthologs found in HDRs (red) or outside HDRs (blue). Box and whisker plots display the median value for each metric on the x-axis. The upper and lower whiskers correspond to the first and third quartiles, respectively. Wilcoxon ranked-sum statistical text with Benjamini-Hochberg FDR-adjusted correction. ****$p$ < 0.0001, ns = not significant.

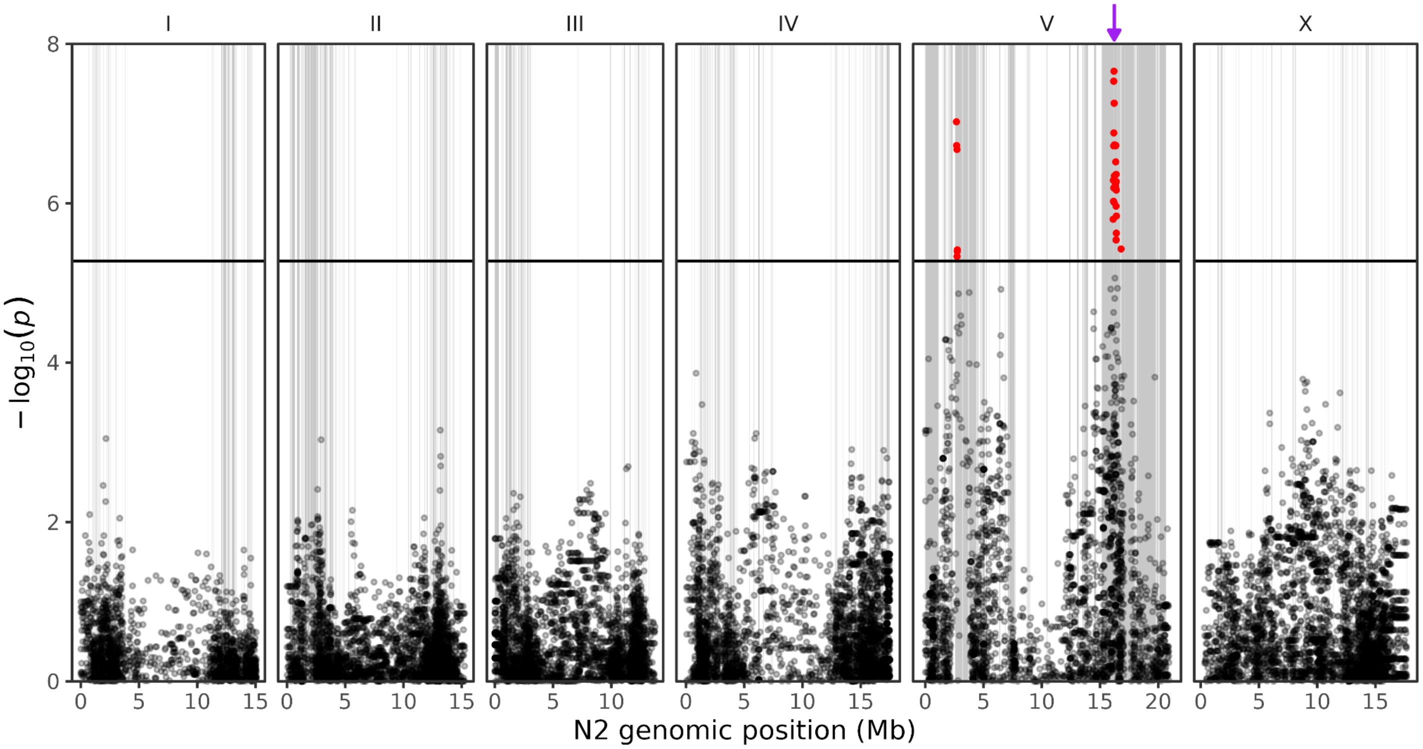


**Supplementary Figure 27. Genome-wide association of SNVs with abamectin response variation.** Manhattan plot of abamectin q90 time of flight responses from larval development assays (Supplementary Tables 11 and 12). Genomic position (x-axis) is plotted against the -log10(*p*) value (y-axis) significance value for each SNV (Supplementary Table 13). SNVs are colored red if they pass the Bonferroni-corrected significance threshold designated by the solid black line. Collapsed HDRs among 237 wild strains that were used in the GWAS are represented in grey, and the location of *glc-1* is represented by the purple arrow.

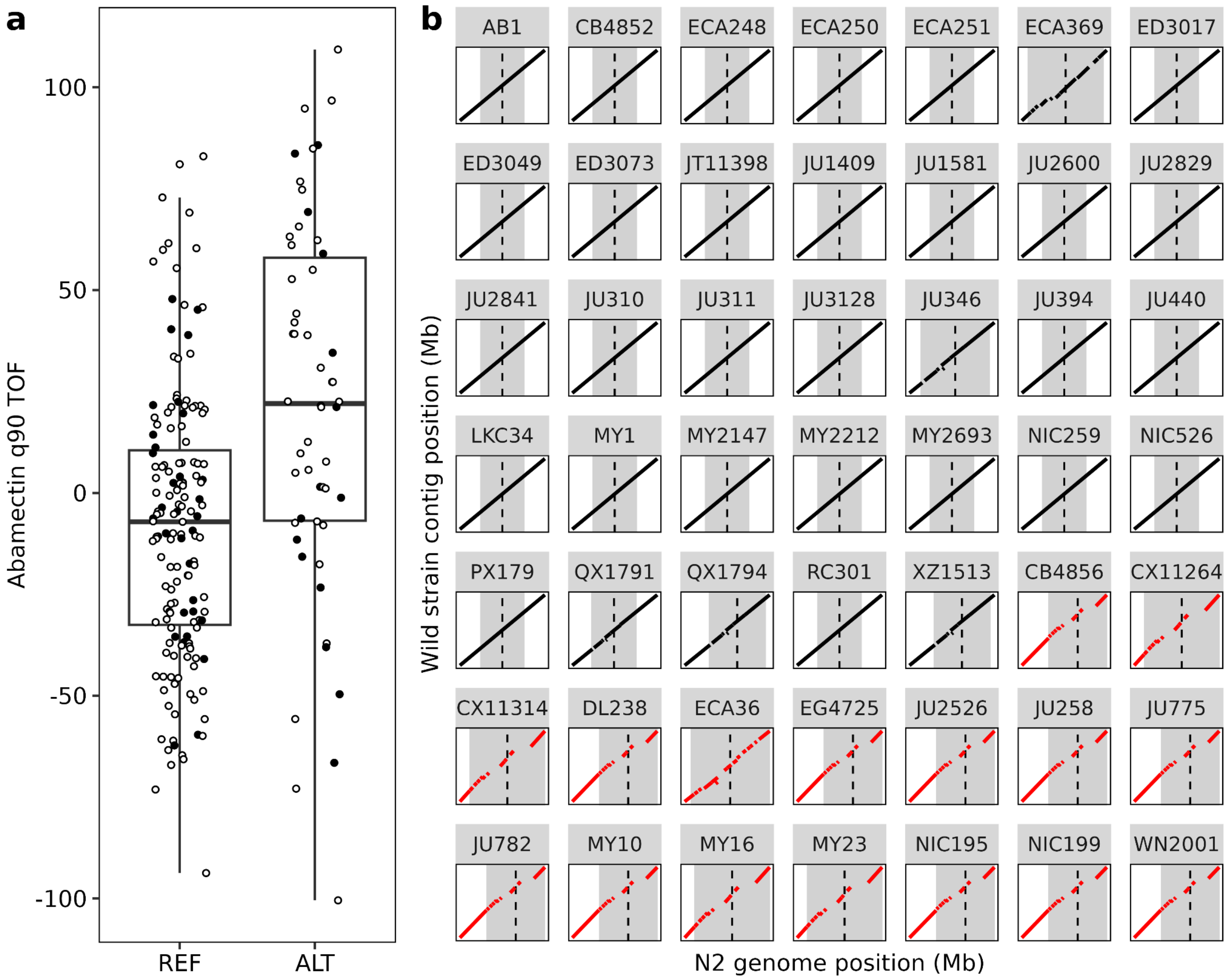


**Supplementary Figure 28. Response to abamectin is correlated with haplotype. a)** Phenotype distribution of abamectin q90 time of flight for wild strains that have the reference (REF) or alternative (ALT) SNV peak marker allele for the QTL on the right arm of chromosome V. Points that are black are wild strains that are in the pangenome. Box and whisker plots display the median value for each metric on the x-axis. The upper and lower whiskers correspond to the first and third quartiles, respectively. **b)** Genome-genome alignments of pangenome strains (black points in **a**)) surrounding the peak marker (vertical dashed line, gray rectangles are 22 kb upstream and 32 kb downstream). REF strains are in black, and ALT strains are in red.

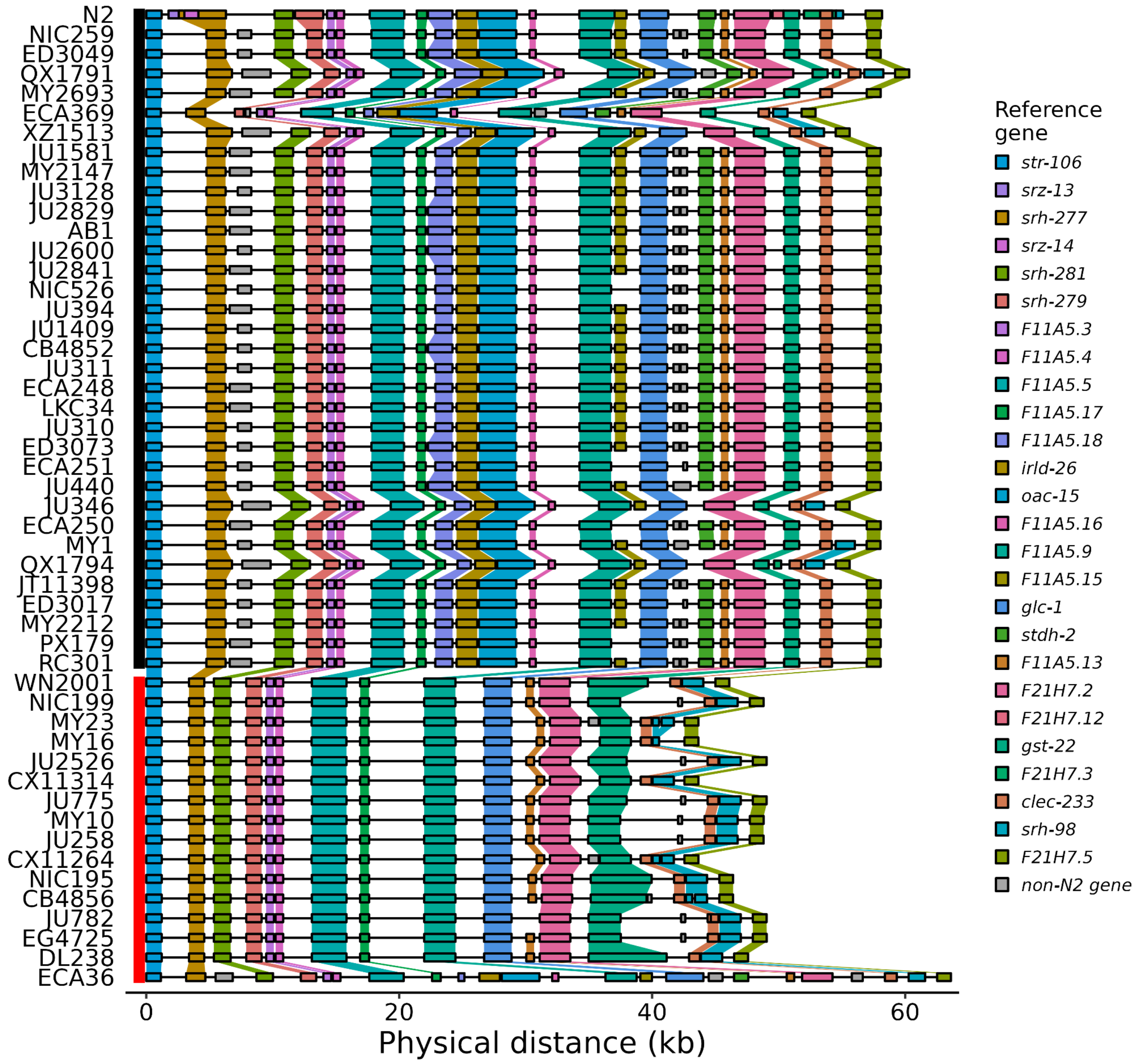


**Supplementary Figure 29. Haplotype visualization of wild strains that are sensitive or resistant to abamectin.** Vertical rectangles represent the peak marker genotype (red = ALT, black = REF). Strains are ordered by increasing abamectin resistance (q90 TOF) within each genotype. Genes are shown as horizontal rectangles, colored by N2 gene identity. Orthologous genes are connected by colored polygons.

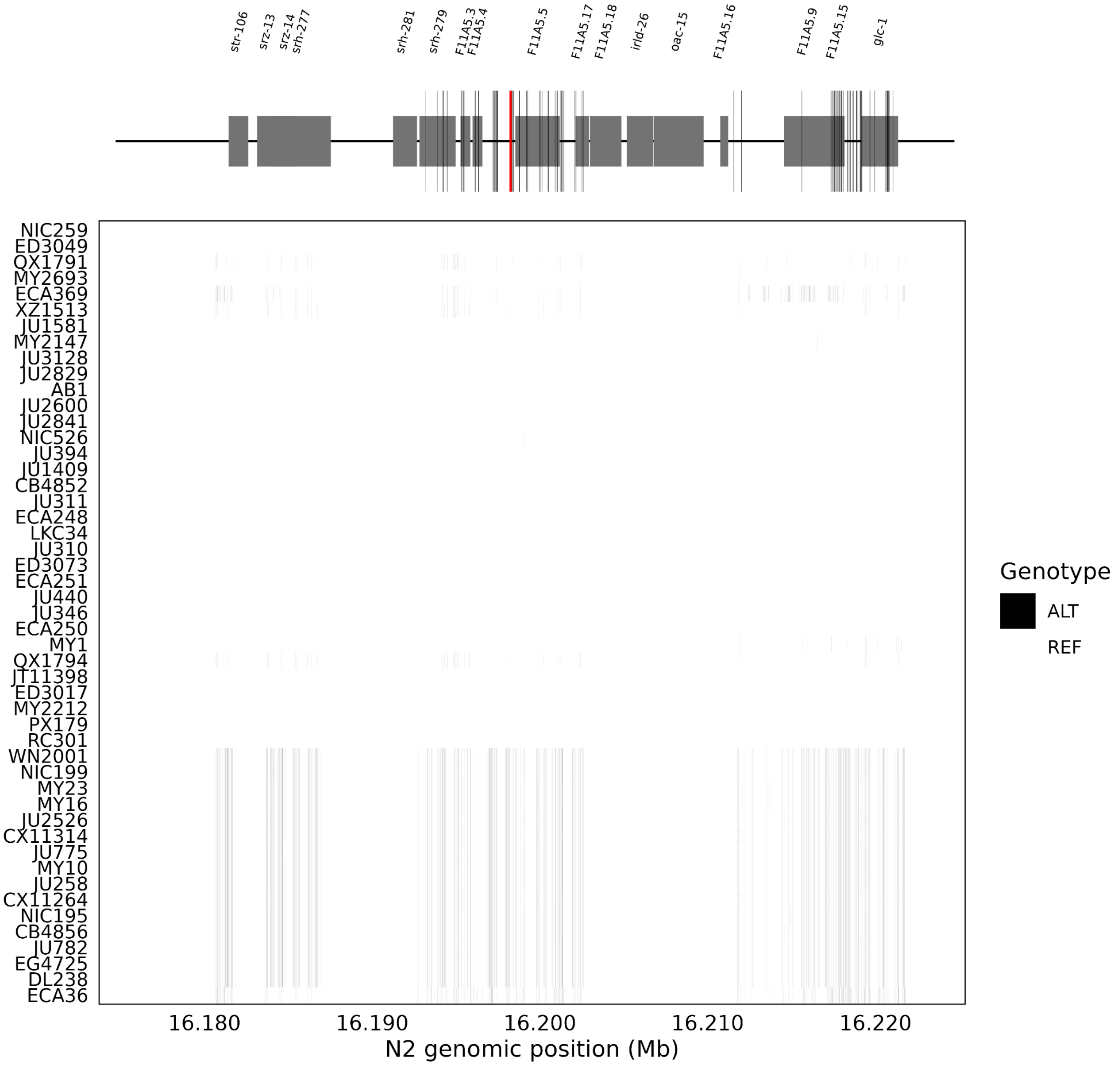


**Supplementary Figure 30. SNVs shared among abamectin resistant strains are concentrated in *F11A5.15* and *glc-1*.** Biallelic SNVs are shown as vertical black tiles. The peak SNV associated with abamectin resistance is indicated by a red vertical line on the gene model plot. SNVs that are unique to abamectin resistance strains are indicated by black vertical lines on the gene model plot.

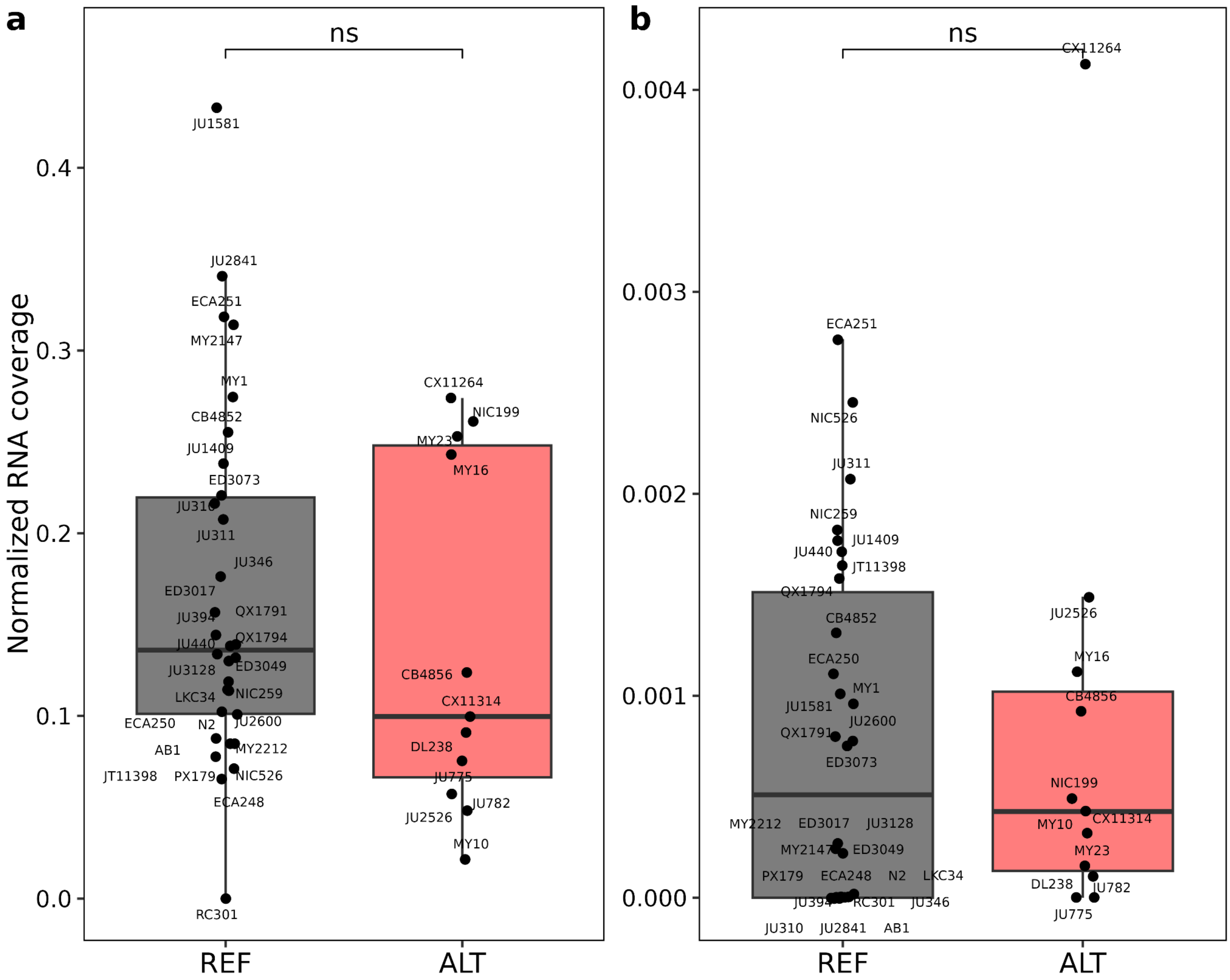


**Supplementary Figure 31. Differences in RNA coverage of *glc-1* and *F11A5.5* among abamectin susceptible and resistant strains.** Average RNA coverage of *glc-1* **(a)** and *F11A5.5* **(b)** normalized by the average exome-wide coverage for 40 of the 50 phenotyped wild strains, faceted by genotype for peak SNV associated with abamectin resistance (Supplementary Figure 28). An unpaired Wilcoxon statistical test with an alpha of 0.05 was used to determine statistical significance.

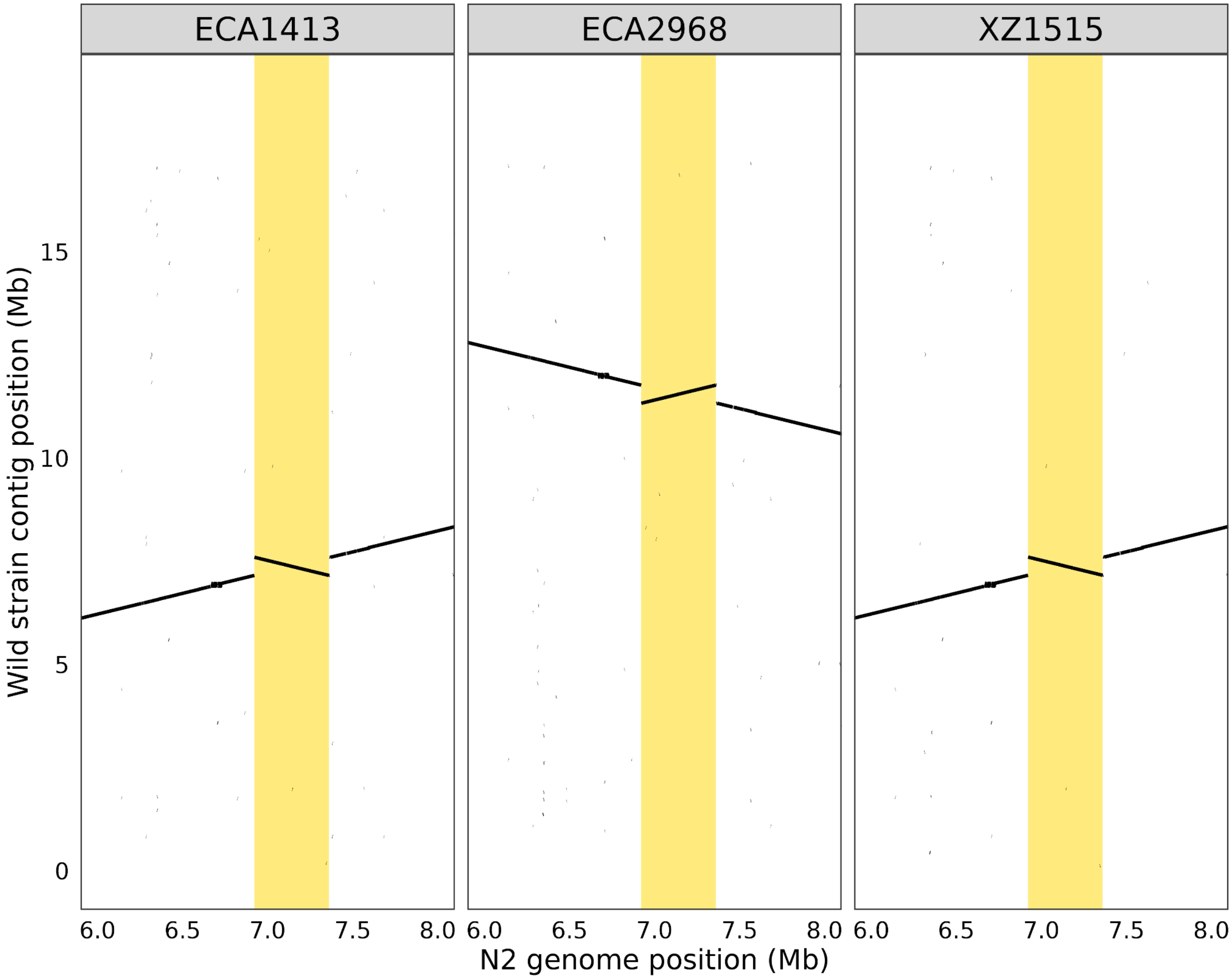


**Supplementary Figure 32. The largest PAV-identified inversion on chromosome IV shared among three Kaua'i strains.** A 438 kb inversion displayed using a dotplot of genome-genome alignments between a wild strain (facet) and N2 on chromosome IV.

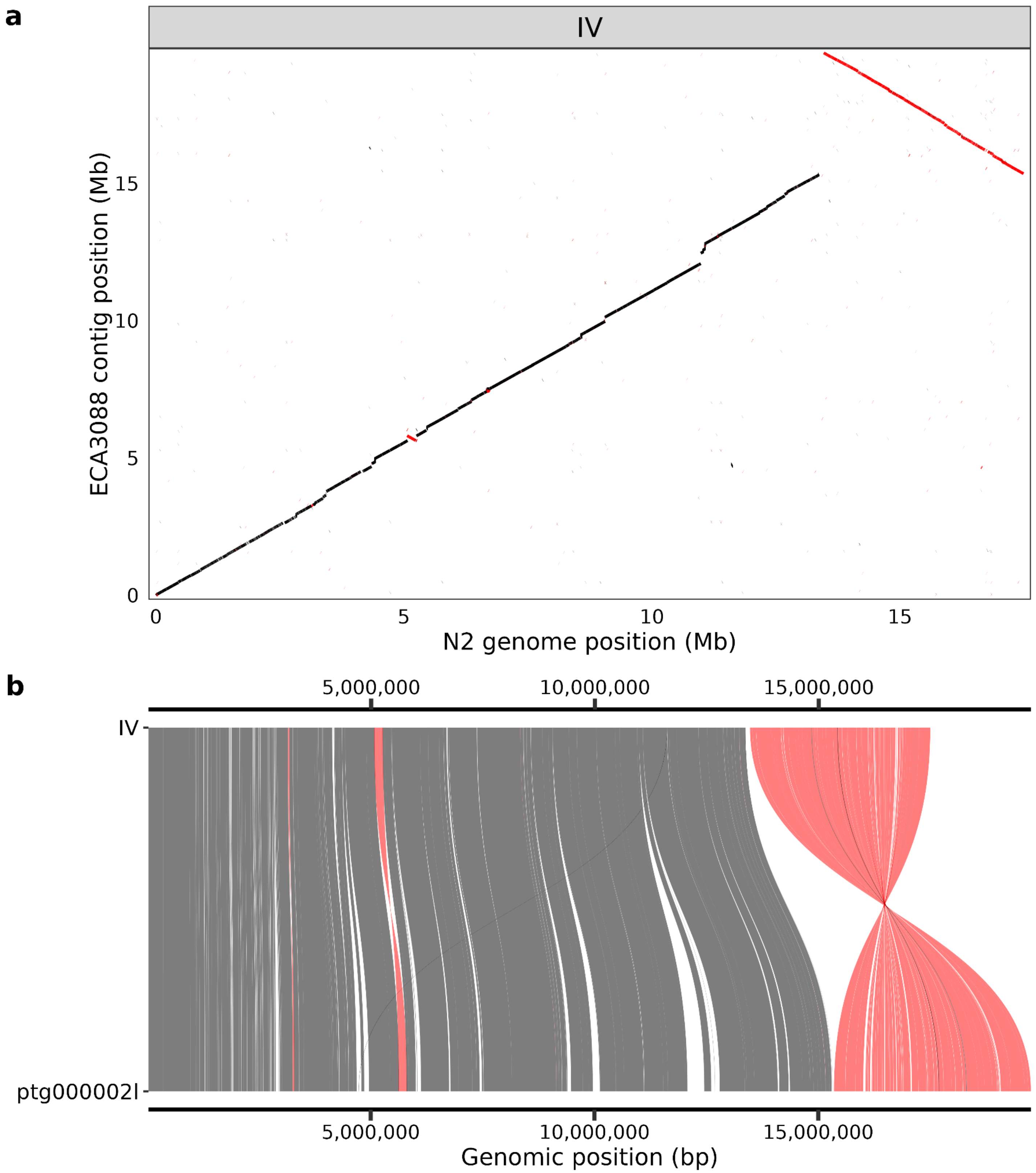


**Supplementary Figure 33. Large inversion on the right end of chromosome IV in ECA3088**. **a)** Genome-genome alignments of ECA3088 to N2 chromosome IV. Alignments in the reverse orientation are displayed in red to highlight inverted alignments. **b)** An SVbyEye[3] plot displaying shared sequence between N2 chromosome IV (top) and ECA3088 contig ptg000002l (bottom) with sequence in the opposite orientation displayed in red.

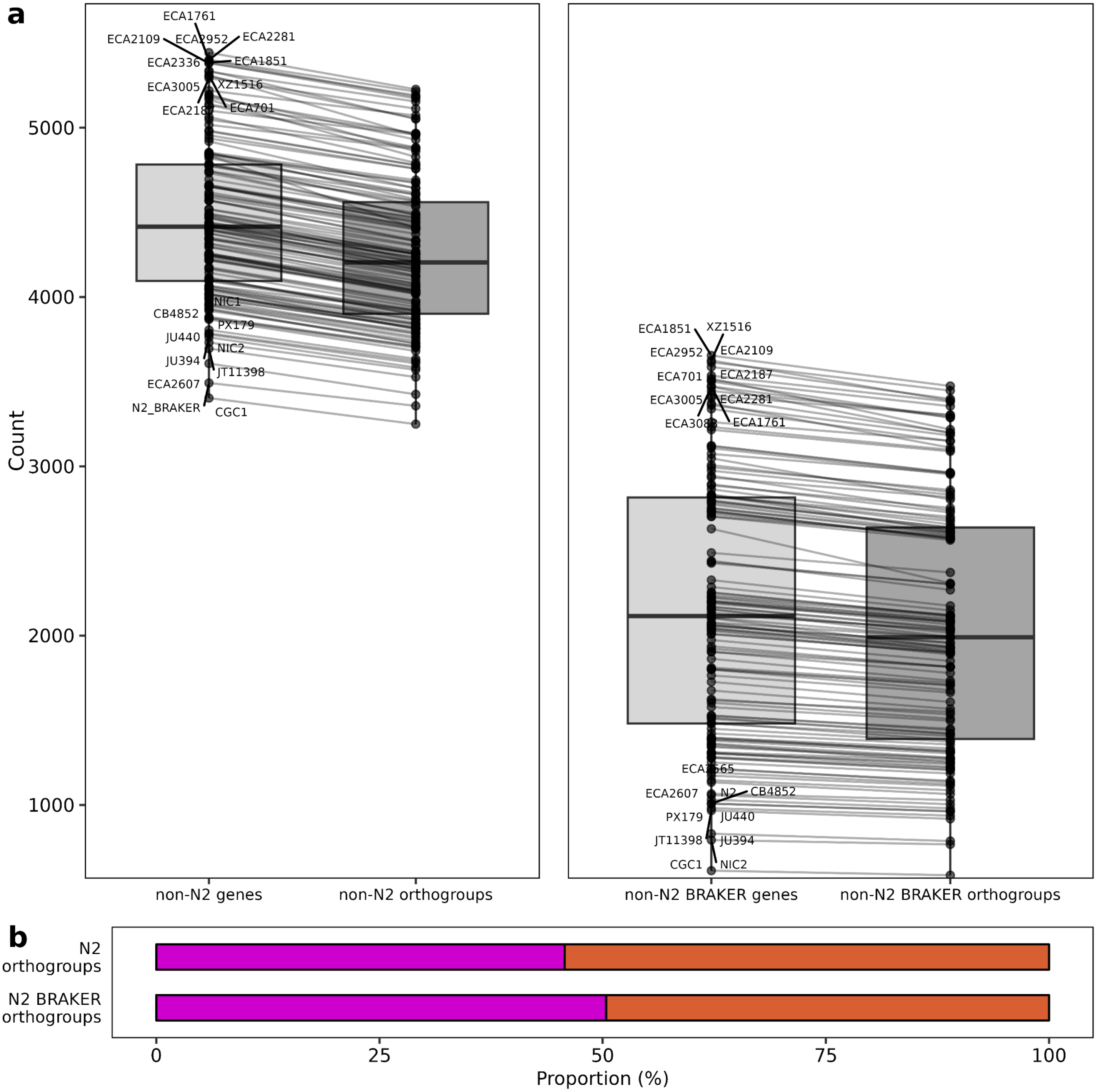


**Supplementary Figure 34. Wild strains have many non-reference genes. a)** The number of orthogroups and genes that are wild-strain-specific or absent in N2 WormBase (WS283) gene annotations (left) or N2 BRAKER3 gene annotations (right). Box and whisker plots display the median value for each metric on the x-axis. The upper and lower whiskers correspond to the first and third quartiles, respectively. **b)** Among all orthogroups that do not have an N2 ortholog for each wild strain, the average proportion of gene sets to which these orthogroups contribute are displayed in pink (private orthogroups) and orange (accessory orthogroups).

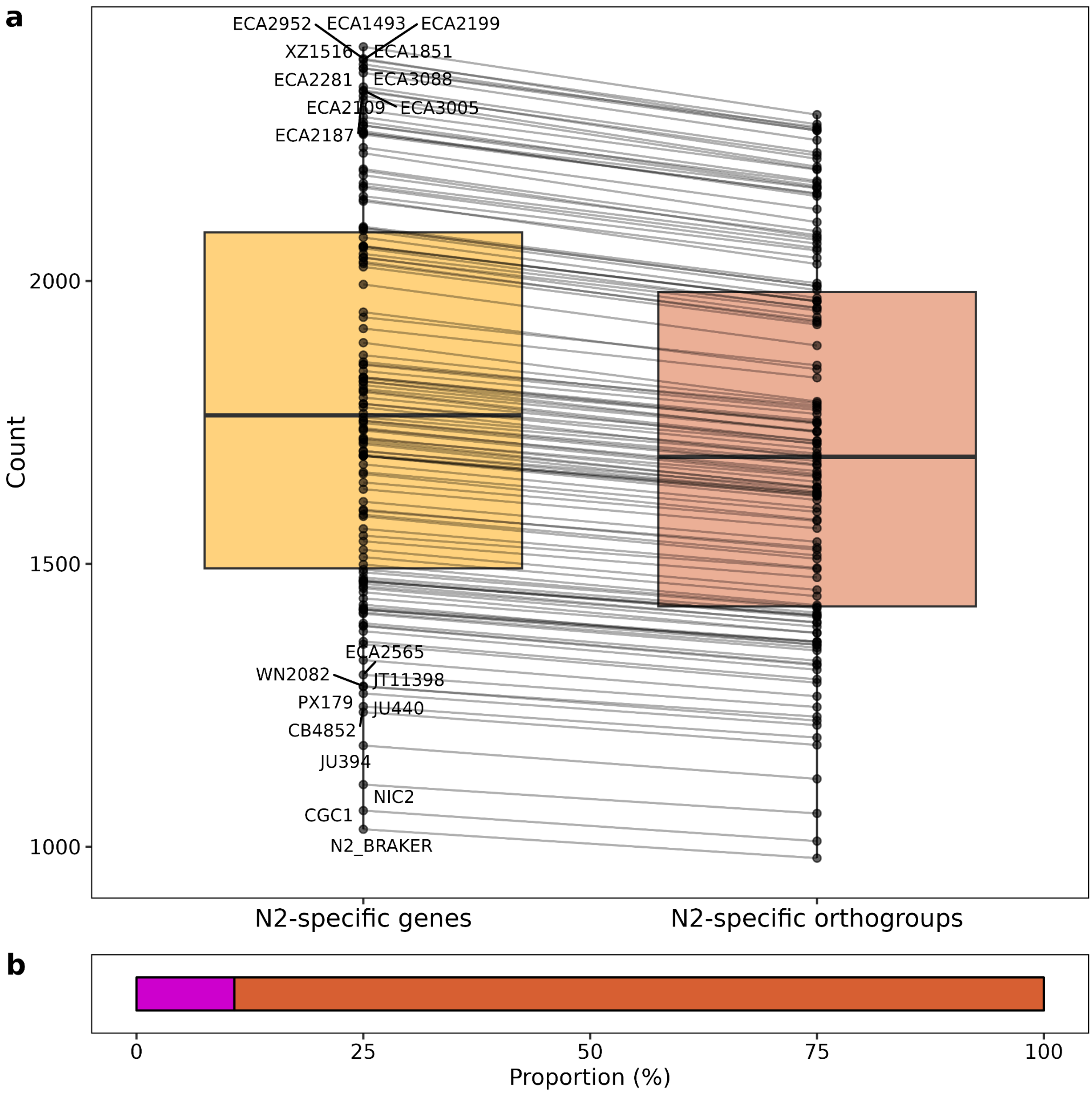


**Supplementary Figure 35. N2 has many genes not found in wild strains. a)** The number of orthogroups and genes that are N2-specific and absent in a given wild strain. Box and whisker plots display the median value for each metric on the x-axis. The upper and lower whiskers correspond to the first and third quartiles, respectively. **b)** Among all orthogroups that do not have a wild strain ortholog for N2, the average proportion of gene sets to which these orthogroups contribute are displayed in pink (private orthogroups) and orange (accessory orthogroups).
